# Aryl amino acetamides prevent the development of *Plasmodium falciparum* rings via inhibition of the lipid transfer protein PfSTART1

**DOI:** 10.1101/2023.11.02.565411

**Authors:** Madeline G. Dans, Coralie Boulet, Gabrielle M. Watson, William Nguyen, Jerzy M. Dziekan, Cindy Evelyn, Kitsanapong Reaksudsan, Somya Mehra, Zahra Razook, Niall D. Geoghegan, Michael J. Mlodzianoski, Christopher Dean Goodman, Dawson B. Ling, Thorey K. Jonsdottir, Joshua Tong, Mufuliat Toyin Famodimu, Betty Kouskousis, Michael J. Delves, Geoffrey I. McFadden, Alyssa E. Barry, Brendan S. Crabb, Tania F. de Koning-Ward, Kelly L. Rogers, Alan F. Cowman, Wai-Hong Tham, Brad E. Sleebs, Paul R. Gilson

## Abstract

With resistance to most antimalarials increasing, it is imperative that new antimalarial drugs are developed to replace or complement front-line artemisinin therapies. We previously identified an aryl acetamide compound, MMV006833 (M-833), that inhibited ring development of newly invaded merozoites. Here, we selected parasites resistant to M-833 and identified independent mutations arising in the START lipid transfer protein (PF3D7_0104200, PfSTART1). Introduction of the identified PfSTART1 mutations into wildtype parasites reproduced resistance to both M-833 and highly potent analogues, confirming PfSTART1 mutations were sufficient to confer resistance. The analogues bound to recombinant PfSTART1 with nanomolar affinity. We also demonstrated selective PfSTART1 engagement by the analogues using organic solvent-based Proteome Integral Solubility Alteration (Solvent PISA) assay for the first time in *Plasmodium.* Imaging of newly invaded merozoites showed the inhibitors prevented the conversion into larger amoeboid ring-stage parasites potentially through the inhibition of phospholipid transfer from the parasite to the encasing parasitophorous vacuole membrane (PVM) and/or within the parasite. We show that these PfSTART1 inhibitors also block transmission. With multiple stages of the parasite’s lifecycle being targeted by PfSTART1 inhibitors, this protein therefore represents a novel drug target with a new mechanism of action.

## Introduction

There were an estimated 247 million cases of malaria in 2021 resulting in approximately 619,000 deaths, mainly in sub-Saharan Africa (World Health Organization, 2022). These numbers remain similar to 2020 when there was >10% increase over previous years due to SARS-CoV-2 related service disruptions. Artemisinin combination therapies (ACTs) remain the frontline treatments for malaria infections by eliminating the causative *Plasmodium* parasites from the body. Of concern is that mutations in the parasite’s *kelch13* gene, a marker of partial resistance to ACTs, have now emerged in many regions around the world and portents to increasing treatment failures (Bergmann et al., 2021; Dondorp et al., 2009; Eastman and Fidock, 2009; Lautu-Gumal et al., 2021; Yoshida et al., 2021). For this reason, new small molecule inhibitors with novel mechanisms of action that can kill various lifecycle stages of the parasite need to enter the developmental pipeline.

A prime process to target is invasion by the free form merozoite into host red blood cells (RBCs) (Burns et al., 2019). This process occurs rapidly in less than a minute (Gilson and Crabb, 2009; Weiss et al., 2016) and requires many protein-protein interactions to successfully occur in a precise, carefully choreographed order (Weiss et al., 2015). To identify inhibitors with novel mechanisms of action against RBC invasion, we previously performed a phenotypic screen of the 400 compound Medicines for Malaria Venture (MMV) Pathogen Box (Dans et al., 2020). Two hits (MMV020291 and MMV006833) displayed promising traits for invasion inhibition (Dans et al., 2020). Recently we described the mode of action of MMV020291, finding that it targeted parasite actin-profilin interactions and prevented merozoites from penetrating the RBC (Dans et al., 2023). In contrast, compound MMV006833 (M-833) had a novel effect whereby the merozoite entered its target RBC but failed to differentiate into an amoeboid ring-stage parasite during the formation of the parasitophorous vacuole membrane (PVM) around the parasite (Dans et al., 2020). Here, we generated M-833 resistant parasites and whole genome sequencing revealed mutations in the *Plasmodium falciparum* Steroidogenic Acute Regulatory protein-related lipid Transfer (START) domain-containing phospholipid transfer protein (PF3D7_0104200, PFA0210c).

START-domain proteins can ferry lipids between membranes and are defined by their START region of ∼210 amino acids, that form a hydrophobic lipid-binding pocket. START domains typically comprise an α/β helix-grip fold with antiparallel β-sheets, flanked by N- and C-terminal α-helices (Iyer et al., 2001). The human START domain family encompasses 15 members (reviewed by (Clark, 2020)), whilst *Plasmodium spp.* In contrast have only five known START-domain proteins (Ressurreição and van Ooij, 2021). Of those five, PFA0210c is the most studied and the focus of this article, which we will refer to as PfSTART1. PfSTART1 shares sequence similarities of 20% with the human StarD7, which is a phosphatidylcholine transfer protein involved in maintaining mitochondrial membrane integrity. However, PfSTART1 is structurally closer to human StarD2, another phosphatidylcholine transfer protein involved in lipid droplets metabolism (Ressurreição and van Ooij, 2021; van Ooij et al., 2013). PfSTART1 can transfer a wide range of phospholipids *in vitro*, with a preference for phosphatidylcholine and phosphatidylinositol (van Ooij et al., 2013). An unusual C-terminal extension has been reported to negatively regulate PfSTART1 activity, and this regulation was essential *in vivo* (Hill et al., 2016). PfSTART1 appears to be vital during the blood stages and most strongly expressed during schizogony (van Ooij et al., 2013). Despite containing a PEXEL motif, PfSTART1 does not appear to be exported to the RBC, but instead remains in the PV (Ressurreição et al., 2023; van Ooij et al., 2013). Overall, PfSTART1 is hypothesised to play a role in forming the PVM upon merozoite invasion by transferring lipids from the RBC or parasite membranes into the nascent PVM, and therefore represents a promising novel drug target (Hill et al., 2016; van Ooij et al., 2013).

Here, we report the M-833 series as the first inhibitors of PfSTART1. In this study, we identify mutations in the START domain of PfSTART1 (N309K, N330K and I224F) in parasites resistant to M-833. Introduction of the mutations responsible for the most resistant phenotypes (N309K and N330K) into 3D7 wildtype parasites conferred resistance to M-833 and its highly potent analogues. We further investigated how the M- 833 series inhibited PfSTART1 function and how this blocks merozoite development into ring-stage parasites.

## Results

### Selection for resistance to MMV006833 followed by whole genome sequencing identified PfSTART1 as a possible target

To select for resistance to MMV006833 (Fig 1A) hereafter called M-833, five populations (labelled A-E) of 10^8^ asexual blood stage 3D7 parasites were cycled on and off with 3 µM of M-833 (corresponding to 10 x EC50 provided by MMV). After three cycles of drug treatment, three parasite populations were identified as less susceptible to the compound and their growth in serially diluted M-833 was measured by lactate dehydrogenase (LDH) activity assays after 72 h. The EC50 of the drug-treated parasites was 2.18 to >10-fold higher than 3D7 wild-type parasites, demonstrating resistance had been generated (Fig S1A). The two most resistant populations (PopD and PopE) were further cloned out by limiting dilution. Growth assays demonstrated the resistance against M-833 was heritable, with a 17-fold increase in EC50 compared to 3D7 parasites (Fig 1B). To identify the target of M-833, genomic DNA was extracted from the clonal lines of PopD and PopE and subjected to whole genome sequencing using MinION technology and compared to clonal 3D7 parental parasites (Dans et al., 2023). This identified 14 non-synonymous single nucleotide polymorphisms (SNP) across nine different genes when compared to the 3D7 parental strain (Table S1). The two genes which had SNPs present across all resistant clones were the *P. falciparum liver stage antigen 1* gene (PF3D7_1036400) and *P. falciparum stAR-related lipid transfer protein* (PF3D7_0104200, PFA0210c; PfSTART1). The former is not expressed in the blood stage (Bozdech et al., 2003; Llinás et al., 2006; Mikolajczak et al., 2011) and comprises a large central repeat region which can be problematic for variant calling. Therefore, we focused on PfSTART1: two of the three sequenced PopD clones (PopD-C6 and -F7) had a SNP (AAC-AAA) which caused a mutation N309K in PfSTART1 (Fig 1C, Table S1). The third PopD clone (PopD-E3) also contained the N309K mutation, although this had been removed during quality filtration as it did not meet the minimum depth of 10x coverage. While we did not sequence the genome of PopD-D7, it also contained the N309K mutation, as demonstrated by PCR amplification and sequencing (Fig S1B). Both PopE clone genomes had a SNP (AAC- AAG) within PfSTART1 that resulted in a mutation N330K (Fig 1C, Table S1). To test whether the N309K and N330K mutations caused a growth defect, PopD-D7 and PopE- F10 were synchronised, and their growth measured with an LDH readout every 48 h. Both lines containing mutations in PfSTART1 had significantly reduced growth compared to 3D7 parasites (Fig 1D). Finally, PCR-amplification and sequencing of PfSTART1 of PopC showed it contained an I224F mutation (Fig S1C). Collectively, all the identified mutations (N309K, I224F, N330K) were found within the central conserved START domain and predicted to face the lipid binding pocket (Fig 1C).

**Figure 1.**
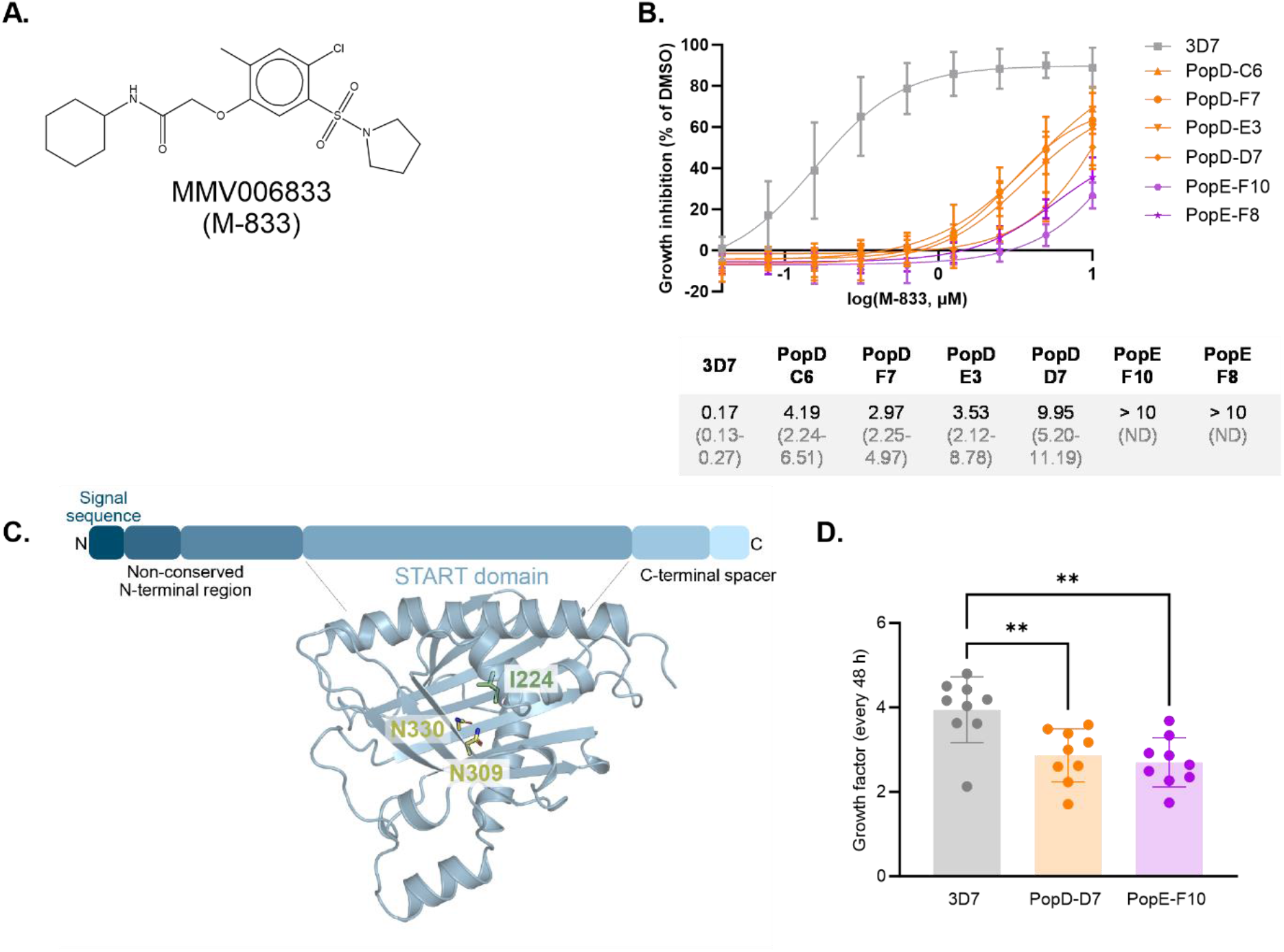
M-833 resistant parasites contain mutations in the *P. falciparum* START1 protein. **(A)** Structure of MMV006833 (M-833). **(B)** Parasites populations (Pop) resistant to M-833 were generated by exposing 3D7 *P. falciparum* parasites to 10 x EC50 of M-833 three times. These parasites were cloned, and growth inhibition assays performed in the presence of 10 µM to 0.04 µM M-833. Growth inhibition was normalised to DMSO controls. EC50 values and 95% confidence intervals are indicated; n=3 biological replicates. **(C)** Key domains and cartoon representation of the AlphaFold predicted structure of the StAR-related lipid transfer protein, PfSTART1 (PF3D7_0104200) (Jumper et al., 2021; Varadi et al., 2022). Mutations that had been identified in the M-833 resistant parasite populations are shown as sticks (yellow = N309K, N330K; green = I224) **(D)** The growth of 3D7, PopD-D7 (N309K mutant) and PopE-F10 (N330K mutant) was measured with an LDH read-out every 48 h (n=9 cycles, over 5 biological replicates). Each dot represents a cycle (LDH48h/LDH0h). Statistical test: Ordinary one-way ANOVA with Tukey’s multiple comparisons test. **: p<0.005.

### Introduction of N309K and N330K mutations into PfSTART1 gene increases resistance to M-833

To test whether PfSTART1 was the target of the M-833 series, the N309K and N330K mutations were first introduced into the *pfstart1* gene using selection linked integration (SLI) (Birnbaum et al., 2017). This construct was designed to integrate via single crossover homologous recombination, inserting the full plasmid into the chromosome (Fig S2A). The native 5’ half of *pfstart1* was used as a homology block, followed by a recodonised 3’ half of the gene (either wild-type (WT), or with the N309K or the N330K mutation), followed by an haemagglutinin (HA) tag at the C-terminus of the protein. A P2A-skip peptide followed by a neomycin resistance gene allowed for selection of parasites that successfully integrated the construct within their genomes. The construct also contained a *glm*S riboswitch to allow for protein knock-down after addition of glucosamine (GlcN). The plasmid had a hDHFR cassette for initial selection of successfully transfected parasites with WR99210. After clonal lines of the WT, N309K and N330K had been obtained, integration was confirmed by PCR (Fig S2B), and the modified *pfstart1* loci were sequenced to ensure the mutations were correct (Fig S2C). The clonal parasites obtained with the SLI method are thereafter called SLI-WT, SLI- N309K and SLI-N330K.

We challenged SLI-WT, SLI-N309K and SLI-N330K with M-833 to determine the effect of the mutations on M-833 treatment (Fig 2A). SLI-N309K and SLI-N330K parasites were 10- and 64-fold more resistant than the 3D7 parental parasites respectively. Unexpectedly, we found that the SLI-WT parasites in which the recodonised WT *pfstart1* sequence was used to replace endogenous coding sequence, were 7-fold more sensitive to M-833 compared to the parental 3D7 parasites.

**Figure 2.**
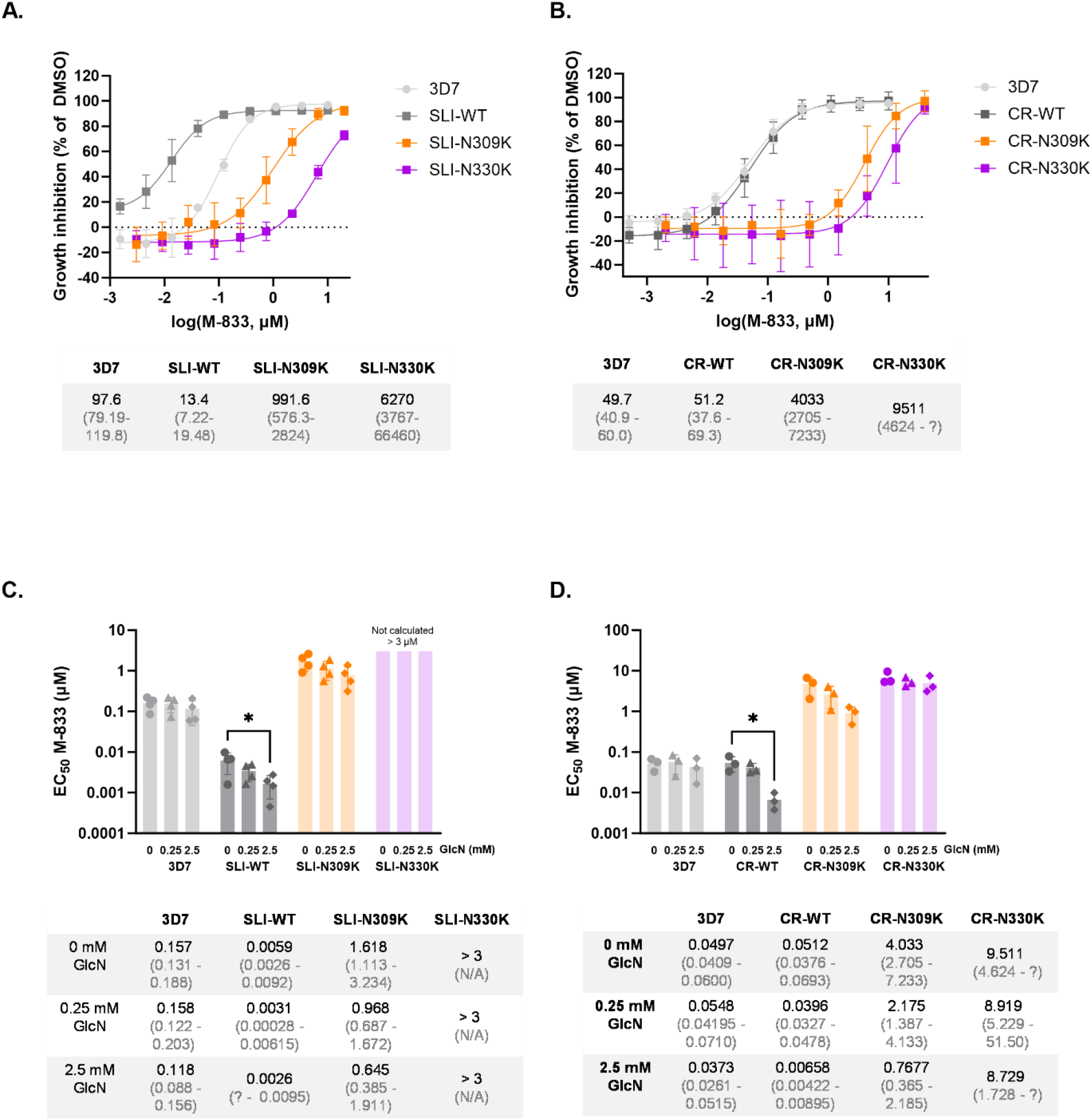
Engineering mutations N309K and N330K in PfSTART1 confers resistance to M-833 and knocking-down PfSTART1 sensitises parasites to M-833. Wild-type (WT) or mutant (N309K, N330K) *pfstart1* genes were introduced in drug-sensitive 3D7 parasites using a Selection-Linked Integration (SLI) method or using CRISPR-Cas9 (CR). **(A, B)** Growth inhibition curves of M-833 on 3D7, **(A)** SLI-parasites, or **(B)** CRISPR- parasites. Parasites were incubated in various concentrations of M-833 and growth was assessed 72 h later using a Lactate Dehydrogenase (LDH) activity assay. EC50 values and 95% confidence intervals are indicated; n=3 biological replicates. **(C, D)** EC50 values (and confidence intervals) of M-833 on 3D7, **(C)** SLI- parasites, or **(D)** CRISPR-parasites in the presence of 0, 0.25 or 2.5 mM glucosamine (GlcN). Engineered parasites contained a riboswitch *glm*S system: when GlcN is added, START is knocked-down (see Fig S2 and S3). n=4 (SLI) and n=3 (CRISPR) biological replicates. Ordinary one-way ANOVA with Dunnett’s multiple comparison test. * p<0.05.

We hypothesised that the addition of the HA tag and P2A skip peptide interfered with PfSTART1 function and caused the sensitisation of SLI-WT parasites to M-833. Therefore, a different cloning strategy was used, whereby the introduction of the mutations in *pfstart1* gene did not include HA and P2A, using CRISPR-Cas9 (McHugh et al., 2021). In comparison to the SLI plasmids, the CRISPR constructs contained an additional 3’ homology block designed to enable double crossover and limited plasmid insertion (Fig S3A). Correct integration was confirmed by PCR (Fig S3B) and the modified *pfstart1* loci was sequenced (Fig S3C). These CRISPR parasite clones are thereafter called CR-WT, CR-N309K and CR-N330K. CR-N309K and CR-N330K parasites exhibited resistance to M-833 to similar extent as the original resistant parasites (over 81- fold resistance compared to CR-WT, Fig 2B). In addition, the CR-WT parasites displayed an EC50 against M-833 comparable to 3D7 (51.2 and 49.7 nM respectively). Overall, these results demonstrate that the mutations N309K and N330K in PfSTART1 confer resistance to M-833.

### Knockdown of PfSTART1 sensitises parasites to M-833

We wanted to examine if knocking down PfSTART1 would impact the EC50 of M-833. First, we tested the knock-down efficiency exerted by the *glm*S riboswitch integrated downstream of the *pfstart1* genes (Fig S2A and S3A).

Full length PfSTART1 is expected to be about 54 kDa, but upon cleavage of the PEXEL motif by plasmepsin V, it appears at ∼48 kDa on western blots (Ressurreição et al., 2023). Western blots of the SLI parasites with an anti-HA antibody labelled a ∼60 kDa band that was larger than the ∼48 kDa native PfSTART1 band detected in 3D7 with an anti-PfSTART1 antibody (Fig S2D). In SLI-N309K there was a smaller band detected that was similar in size to the native protein and contained no HA-tag. The source of this smaller PfSTART1 protein is unknown: the line was cloned again, and all derived parasite lines also contained the two PfSTART1 bands; diagnostic PCRs of the SLI-N309K parasites indicated the native gene was tagged (Fig S2B). Western blots of the CRISPR tagged parasites using the anti-PfSTART1 antibody detected proteins of the expected size (Fig S3D).

Upon addition of 2.5 mM glucosamine (GlcN) for 48 h, PfSTART1 was successfully knocked down in schizonts in both the SLI and the CRISPR parasites (Fig S2D and S3D). The knockdown was stronger in the CRISPR constructs (75% for CR-WT, 81% for CR- N309K, 84% for CR-N330K) compared to the SLI parasites (60% for SLI-WT, 43% for SLI-N309K, 70% for SLI-N330K). This was expected as the *glm*S riboswitch is located further downstream of the *pfstart1* coding sequence in the SLI constructs compared to the CRISPR constructs.

To measure the impact of PfSTART1 knockdown on parasite growth, ring stage parasites (3D7, SLI- and CR- WT, N309K, N330K) were exposed to 0, 0.25 or 2.5 mM GlcN for 72 h, and their growth was assessed by LDH activity (Fig S2E and S3E). In the presence of 2.5 mM GlcN, all SLI- and CR- parasites grew significantly less than in the absence of GlcN: approximately 40% less for most lines, and down to 50% less in the case of CR-N309K. 3D7 parasites were slightly impacted by the presence of 2.5 mM GlcN, but to a lesser extent, with ∼5% reduced growth. The data therefore indicates that reduction in expression of PfSTART1 reduces parasite growth.

To determine if knockdown of PfSTART1 also increased the sensitivity of the parasites to M-833, a growth inhibition assay was performed in the presence of 0, 0.25 or 2.5 mM GlcN, and the EC50 calculated (Fig 2C and 2D). In both the SLI-WT and the CR-WT parasites, the addition of 2.5 mM GlcN significantly reduced the EC50 values: by 2.3-fold for more sensitive SLI-WT; 7.8-fold for CR-WT. In the case of the N309K mutation, the M-833 EC50 values were reduced by similar factors to the WT parasites in the presence of 2.5 mM GlcN (2.5-fold reduction for SLI-N309K, 5.2-fold reduction for CR-N309K), although this did not reach statistical significance. Regarding the N330K mutants, the addition of GlcN did not impact the EC50 of M-833 in CR-N330K parasites. As expected, the addition of GlcN did not change the EC50 of control 3D7 parasites. Together, these results confirm that M-833 targets PfSTART1, and that PfSTART1 is important for parasite growth.

### M-833 resistant parasites are also resistant to highly potent analogues

To create more potent probes to investigate the target of M-833 as well as to explore the therapeutic potential of the compounds, we explored the structure activity relationship of the M-833 series. It was observed that M-833 shared structural similarities with the aminoacetamide compound **2** (Norcross et al., 2019) (Fig 3A), which is developed from a Tres Cantos Antimalarial Set (TCAMS) hit. The structural similarities include a substituted aryl sulfonamide, acetamide core and aryl amido substitution in both series. As the mechanism of action for the Norcross *et al*. series has not been determined, its similarities to M-833 suggests it may share the same target. Compound **2** was assessed for activity against M-833 resistant populations and was shown to be cross-resistant (Fig 3B), indicating compound **2** is likely to have the same molecular target as M-833. In the optimisation of M-833 it was found that there was crossover with the structure activity relationship between the two structurally related hit compounds (Norcross et al., 2019). Installation of a gem-dimethyl on the alpha position of the acetamide generated the compound **W-991** (WEHI-991), resulting in a 20-fold improvement in parasite activity (EC50 = 7 nM) (Fig 3A). This change also led to a small increase in human HepG2 cell activity (CC50 = 29 µM). This activity is consistent with that previously reported (Norcross et al., 2019). Replacing the phenolic group (**3**) in M-833 with an anilino group in **W-991** resulted in a 20-fold improvement in parasite activity (EC50 = 150 nM), highlighting the preference for a secondary amine in that position. Methylation of the amido group resulted in an inactive analogue (compound **4**) (EC50 > 10 µM) which was used as a negative control for our studies (Fig 3A).

**Figure 3.**
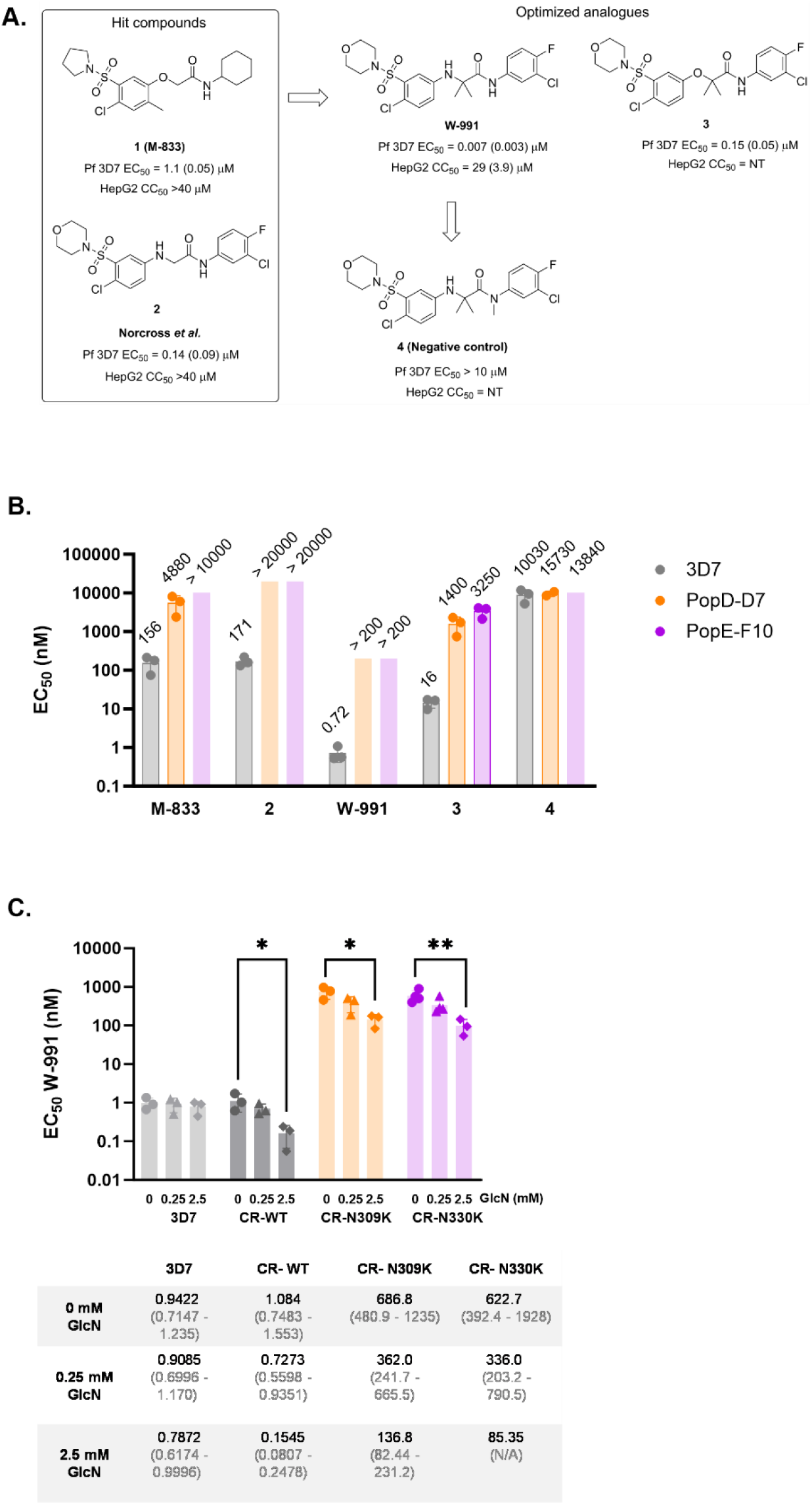
Analogues of M-833 are resisted by M-833 resistant lines, SLI- and CRISPR-mutants. **(A)** Optimised analogues of M-833: their structures, EC50 (SD) on 3D7 parasites, toxicity (CC50 (SD)) on HepG2 cells. NT = not tested. **(B)** EC50 values of M-833 and analogues on 3D7 and M-833 resistant parasites PopD-D7 and PopE-F10, containing the PfSTART1 mutations N309K and N330K, respectively. n=3 biological replicates. EC50 values indicated above the bar, as well as in Table S2. Growth curves available in Fig S4. **(C)** EC50 values (and 95% confidence intervals) of W-991 on 3D7, CR-WT, CR-N309K and CR-N330K parasites, in the presence of 0, 0.25 or 2.5 mM glucosamine (GlcN). n=3 biological replicates. Ordinary one-way ANOVA with Dunnett’s multiple comparison test. *: p<0.05. **: p<0.005.

These newly synthesised analogues were tested against the original M-833 resistant parasites PopD-D7 and PopE-F10 which contain the PfSTART1 N309K and N330K mutations respectively. Both resistant clones were highly resistant to the active analogues (compound **2**, **W-991**, and compound **3**) (Fig 3B, Fig S4 and Table S2). No change in EC50s was observed between the resistant lines and 3D7 for the inactive analogue **4**. Growth inhibition assays with GlcN were also carried out on the CRISPR parasites, using the potent analogue W-991 (Fig 3C). The first observation was that the CR-N309K and CR-N330K parasites were around 600-fold more resistant than the 3D7 and the CR-WT controls. This demonstrates that the PfSTART1 mutations confer resistance to W-991. Despite this impressive resistance, it should be noted that the mutant parasites had reasonably low EC50 (both below 700 nM). When PfSTART1 was knocked down with the addition of GlcN, all CRISPR parasites became significantly sensitised to W-991 (7-fold for CR-WT, 5- fold for CR-N309K, 7.3-fold for CR-N330K), while the addition of GlcN did not change the EC50 on control 3D7 parasites. These results show that we successfully developed potent analogues of M-833, that all appear to target PfSTART1.

### M-833-series binds to recombinant PfSTART1 protein and PfSTART1 in parasites, but not to the mutant PfSTART1(N330K) protein

To independently confirm that PfSTART1 was the target for the M-833 series, we expressed recombinant wildtype PfSTART1(WT) (I149-V394) and measured binding to M-833, W-991 and negative control compound **4** by isothermal titration calorimetry (ITC) (Fig 4A, Fig S5A-C). M-833 bound PfSTART1(WT) with nanomolar affinity (KD = 42 ± 12 nM), and was both enthalpically and entropically favoured, which is an optimal thermodynamic signature for inhibitor/target interactions (Fig 4A.ii). The optimised inhibitors, W-991, had improved binding affinity for PfSTART1 (KD = 10 ± 7 nM) compared to M-833. This improved binding affinity can be largely attributed to an enhanced ΔH, suggesting improved hydrogen bond contributions. PfSTART1(WT) did not interact with the chemically related but inactive analogue **4**. To investigate the effect of the resistant mutations to compound binding we also recombinantly expressed PfSTART1(N309K) and PfSTART1(N330K) (I149-V394) (Fig S5A,B). Since the former had poor yield, only PfSTART1(N330K) was evaluated in ITC binding experiments. This revealed no interaction with potent analogue W-991 (Fig 4A, Fig S5C), indicating the lysine substitute at the 330 position was the mechanism of parasite resistance in PopE parasites.

**Figure 4.**
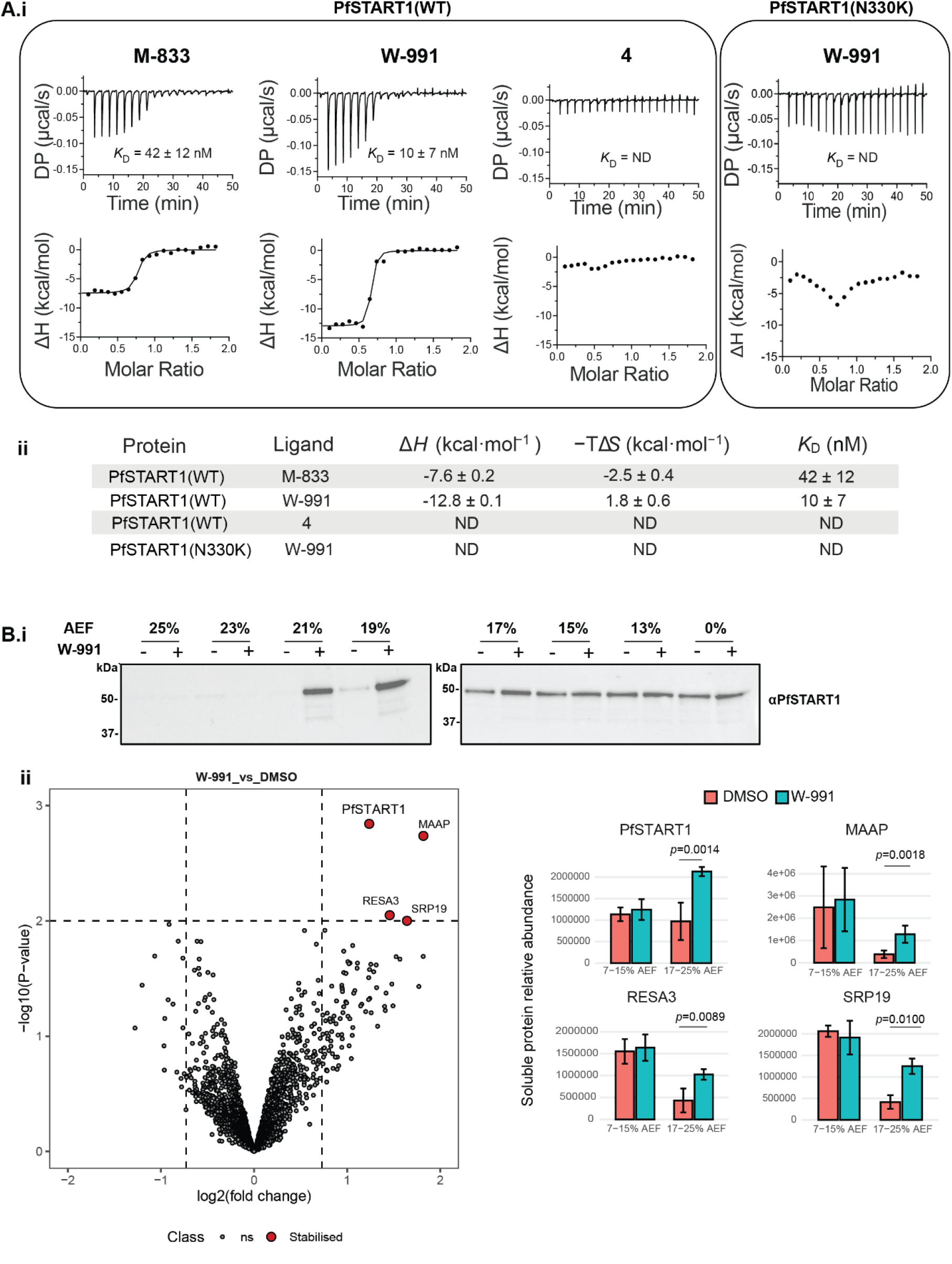
The M-833 series demonstrate target engagement for PfSTART1 while the mutant form prevents compound binding. **(A)** Isothermal titration calorimetry (ITC) analysis of M-833 series and recombinant PfSTART(WT) or PfSTART(N330K). **i.** Representative thermograms of 90 µM PfSTART1 titrated into 10 µM M-833, W-991 and negative control **4**. The bottom panel comprises the data after integration of the peaks and a fitted offset applied. The binding curve shows the fit to a single-site binding model. DP = differential power. **ii.** Summary of the mean affinity, enthalpy and entropy obtained for M-833, W-991 and **4** binding to PfSTART1 from n=2 experiments. Error represents standard deviation. ND = not determined. The second replicate and values for individual parameters are provided in Fig S5. **(B)** Solvent proteome profiling assays. **(i)** Parasite lysate was treated with DMSO or 10 µM W-991, challenged with an acetic acid/ethanol/formic acid mixture (AEF) and soluble fractions extracted and analysed via western blots probed with anti-PfSTART1. Western blot replicates are shown in Fig S5B. **(ii)** Volcano plots depict differential soluble protein abundance analysis of parasite lysate treated with W-991 (100 µM) or the DMSO control after solvent-induced protein precipitation (7-25% AEF). Non-significant (ns) proteins are plotted in grey, significant stabilised proteins in red. Hit selection cut-offs of 0.73 log2 fold-change and p < 0.01 are indicated with dashed lines. Significant stabilisation hits are shown in the bar graphs representing relative soluble protein abundance in DMSO- or W-991-treated parasite lysate samples after solvent-induced protein precipitation, plotted for Gradient 1 ‘G1’ (7- 15% AEF) and Gradient 2 ‘G2’ (17-25% AEF). Error bars represent the standard deviation of three biological replicates.

Next, in order to assess the specificity of the M-883 series we carried out proteome-wide drug-target engagement profiling. A routine method that is now used widely for this in the antimalarial field are thermal shift assays (Bopp et al., 2023; Dziekan et al., 2020; Dziekan et al., 2019; Favuzza et al., 2020). However, we found that PfSTART was thermally stable up to 75°C in parasite lysate (data not shown), in accordance with a previous proteome study (Dziekan et al., 2019). Instead, we leveraged the principle of Solvent Induced Protein Precipitation (SIPP). SIPP identifies protein-ligand interactions based on differential susceptibility of ligand-bound proteins to denaturation by organic solvents (Van Vranken et al., 2021; Yu et al., 2023; Zhang et al., 2020). Here, soluble parasite protein lysate was exposed to increasing concentrations (0-25%) of a mixture of acetone, ethanol and formic acid (AEF) in the presence of W-991 or DMSO, followed by soluble protein isolation via centrifugation. Soluble proteins were separated out on an SDS-PAGE and probed with an PfSTART1 antibody (Fig 4B.i). This showed that W-991 elicited protection to PfSTART1 aggregation at 19-21% AEF (Fig 4B.i, Fig S5D). To further investigate the target engagement of W-991, we next turned to a global proteome analysis in the SIPP assay. After solvent challenge, soluble protein fractions were subsequently combined into two samples representing low and high denaturation pressure (7-15% and 17-25% AEF, respectively), following a Proteome Integral Solubility Alteration (PISA) experimental format (Gaetani et al., 2019). Relative protein abundance was subsequently determined through Data Independent Acquisition mass spectrometry (DIA-MS) analysis (2,393 *Plasmodium* proteins with ≥3 peptides), followed by differential abundance analysis of drug- and vehicle-treated samples. This ‘Solvent-PISA’ assay revealed four proteins exhibiting significantly (p< 0.01) increased levels in the presence of W-991, of which PfSTART1 was the most significant (Fig 4B.ii, *p*=0.0014). The three other drug-stabilized proteins included merozoites-associated armadillo repeats protein (MAAP, PF3D7_1035900), ring-infected erythrocyte surface antigen (RESA3, PF3D7_1149200) and signal recognition particle subunit (SRP19, PF3D7_1216300) (Fig 4B.ii). The abundance/stability of the other two START-domain containing *P. falciparum* proteins detected in the assay was not affected by W-991 treatment (Fig S5E). Taken together, the ITC and Solvent-PISA experiments strongly support PfSTART1 as the principal molecular target of the M-833 inhibitor series.

### PfSTART1 inhibitors block ring development but their effect is reversible

M-833 treated parasites have been observed to invade normally but to stall before ring development (Dans et al., 2020). To investigate whether parasites could recover from PfSTART1 inhibition, parasites were treated similarly as in the Dans *et al*. study, and then followed-up for 3 days. Late-stage schizonts were treated for 4 h (during the egress/invasion window) with M-833, W-991, ML10 (reversible egress inhibitor), E64 (irreversible egress inhibitor), and heparin (invasion blocker), after which the compounds were removed, non-egressed schizonts eliminated with a sorbitol treatment, and parasite growth and phenotype monitored over 3 days (Fig 5A). Growth was assessed every 24 h via luminescence of nanoluciferase (Nluc) in parasites expressing Hyp1-Nluc (Azevedo et al., 2014; Barnes et al., 2022). All treatments significantly reduced parasite growth compared to DMSO (Fig 5A.ii), with M-833, W-991 and heparin displaying the most striking difference, with virtually no recovery after three days (72 h). A short 4 h treatment was also sufficient to affect the morphology of the treated parasites for days: at 72 h, M- 833 and W-991-treated parasites still resembled the dysmorphic ring-stage parasites seen previously in both chemical inhibition by the M-833 series and genetic knockout of PfSTART1 (Fig 5A.i) (Dans et al., 2020; Hill et al., 2016).

**Figure 5.**
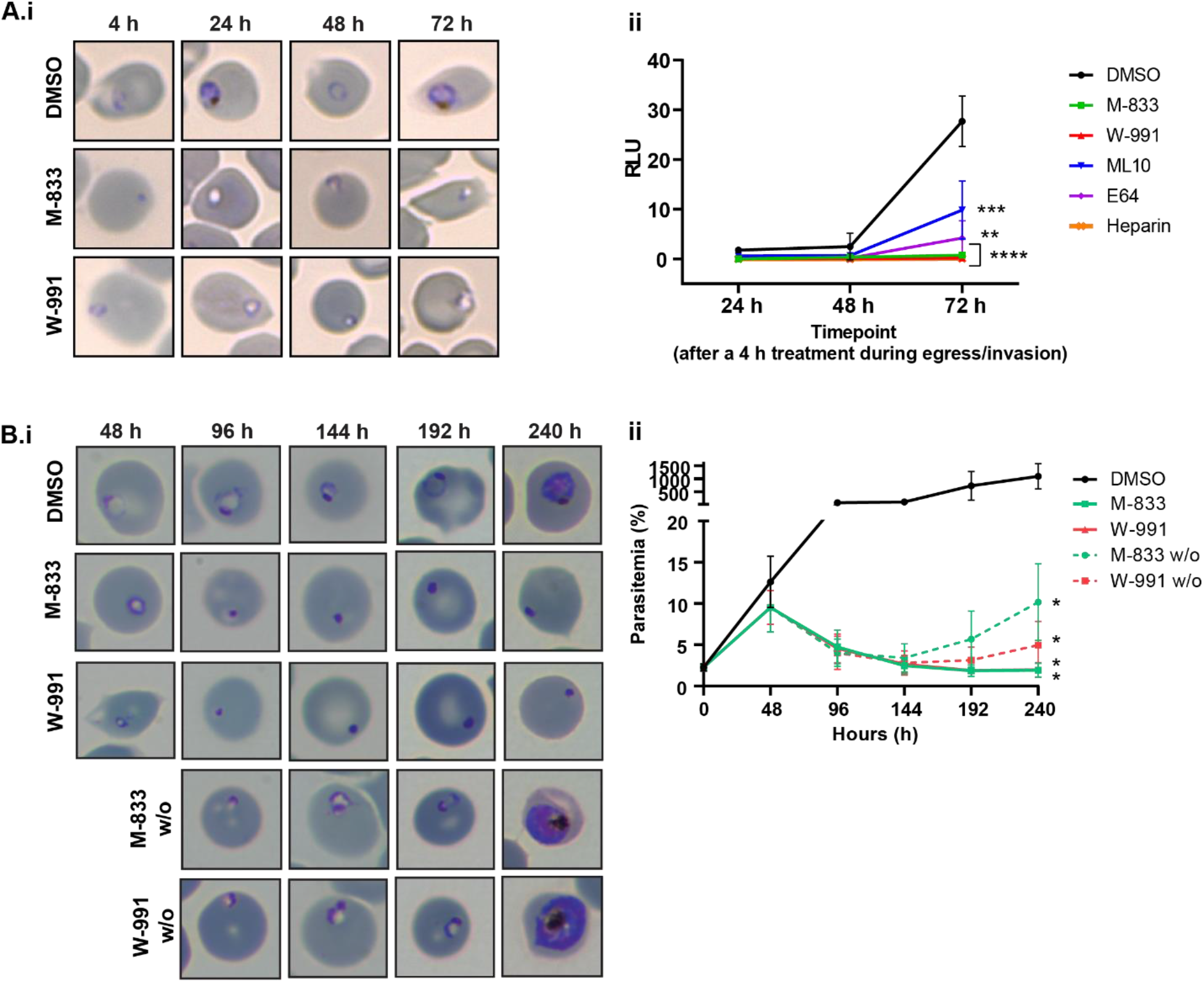
PfSTART1 inhibitors block ring development but their effect is reversible. **(A)** Tightly synchronised Hyp1-Nluc schizonts were exposed to the following drugs for 4 h (during the egress/invasion window): M-833 (2µM), W-991 (60 nM), DMSO (0.02%), ML10 (30 nM) (egress inhibitor), E64 (10µM) (irreversible egress inhibitor), and heparin (100 µg/mL) (invasion inhibitor). After 4 h, non-egressed schizonts were eliminated with a sorbitol treatment, the drugs washed off, and parasites were followed over 72 h. Smears were taken at each timepoint **(A.i)** and growth was assessed with a luminescence read-out **(A.ii)**. n=3 biological replicates. Ordinary one-way ANOVA with Tukey’s multiple comparison test conducted on the 72 h timepoint. *: p<0.05 (ML10 vs. Heparin; not shown on the graph). ***: p<0.005 (DMSO vs. ML10 and E64). ****: p<0.0001 (DMSO vs. M-833, W-991 and heparin). **(B)** Highly synchronous ring-stage 3D7 parasites were exposed to M-833 (2 µM), W-991 (60 nM) or DMSO (0.02%). After 48 h, populations of M-833 and W-991-treated parasites were treated with sorbitol, compounds were washed out (w/o) (dotted lines), and morphology visualised by Giemsa-stained thin blood smears **(B.i)** and parasitemia was quantified by SYBR Green staining and flow cytometry **(B.ii)** every 48 h for four following cycles of growth. Error bars indicate standard deviation of two biological replicates. Statistical analyses conducted via one-way ANOVA, comparing DMSO vs. other treatments. *indicates p <0.05.

After confirmation that the M-833 series was blocking normal ring-stage development, we sought to determine (1) whether other intraerythrocytic stages were affected and (2) whether this resulted in parasite death or stasis beyond 72 h. To do this, we treated highly synchronous ring-stage parasites with DMSO, M-833 or W-991 and followed them through a cycle of growth, which did not indicate any impact on already developed rings, trophozoites or schizonts (Fig S6A). The presence of dysmorphic rings in the M-833 and W-991-treatments was confirmed after 48 h (Fig 5B.i). Drug-treated samples were then exposed to sorbitol lysis to remove unruptured schizonts and compounds were washed out. Parasites were then returned to culture alongside their continuously treated counterparts. This revealed that after 6 days post washout (T=192 h), parasites that had been pre-treated with either M-833 or W-991 were able to resume normal growth, which was visible by both the increase in parasitemia as measured by SYBR Green staining (Fig 5B.ii) and Giemsa-stained blood smears (Fig 5B.i), indicating that one cycle of treatment is not sufficient to cause irreversible inhibition of parasite growth. To corroborate this, we performed parasite reduction ratio (PRR) assays whereby the parasites were exposed to W-991 for up to 5 days before the compound was removed and parasites returned to culture for 21 days. This revealed that despite a reduction of viable parasites after an increasing exposure time to W-991, a proportion of parasites remained after a treatment period of 5 days (Fig S6B.i-ii). Taken together, these experiments show that complete inhibition of parasite growth with the PfSTART1 inhibitors requires >5 days of continuous treatment and parasite survival with <5 days of treatment is likely due to stasis in the ring-stage of the asexual lifecycle.

### PfSTART1 inhibitors block differentiation into ring-stage parasites through membrane disruption

It has been previously postulated that the phospholipid transfer activity of PfSTART1 may be required for the expansion of the parasitophorous vacuole (PV) after invasion (van Ooij et al., 2013). Since previous live cell imaging of merozoites treated with M-833 also showed a defect in ring-stage establishment (Dans et al., 2020), we utilised lattice light sheet microscopy to visualise PVM formation directly after merozoite invasion in the presence of our potent PfSTART1 inhibitor. To track these events, the RBC membrane was labelled with a fluorescent steryl dye, Di-4-ANEPPDHQ and merozoites with Mitotracker Deep Red dye (Geoghegan et al., 2021). Consistent with previous reports (Geoghegan et al., 2021; Gilson and Crabb, 2009), several minutes after invasion in the DMSO control, the PVM was observed to become less spherical and more irregularly shaped consistent with the differentiation of the spherical merozoite into an amoeboid ring (Fig 6A.i, Movie S1). In contrast, newly invaded merozoites treated with W-991 remained spherical over this period (Fig 6A.i, Movie S2). Bioinformatic analysis of 4- dimensional images of nascent PVMs indicated the mean sphericity of the PVM of W-991-treated merozoites stayed high during the observation period compared to the DMSO control where the PVM became less spherical consistent with change in shape to accommodate the amoeboid ring (Fig 6A.ii, Fig S7A, p=0.045 at 15 min). The mean vacuole surface area and volume of the PVM were also found to be reduced in W-991 treatments but this was found not to be significant (Fig S7B-C; p=0.109, p=0.129, respectively). Overall quantification of the lattice light sheet microscopy supports our earlier empirical observations that inhibition of PfSTART1 prevents normal development of the parasite plasma membrane and/or PVM to help expand the PV space, thereby reducing the capacity of the merozoite to differentiate into a ring-stage parasite.

**Figure 6.**
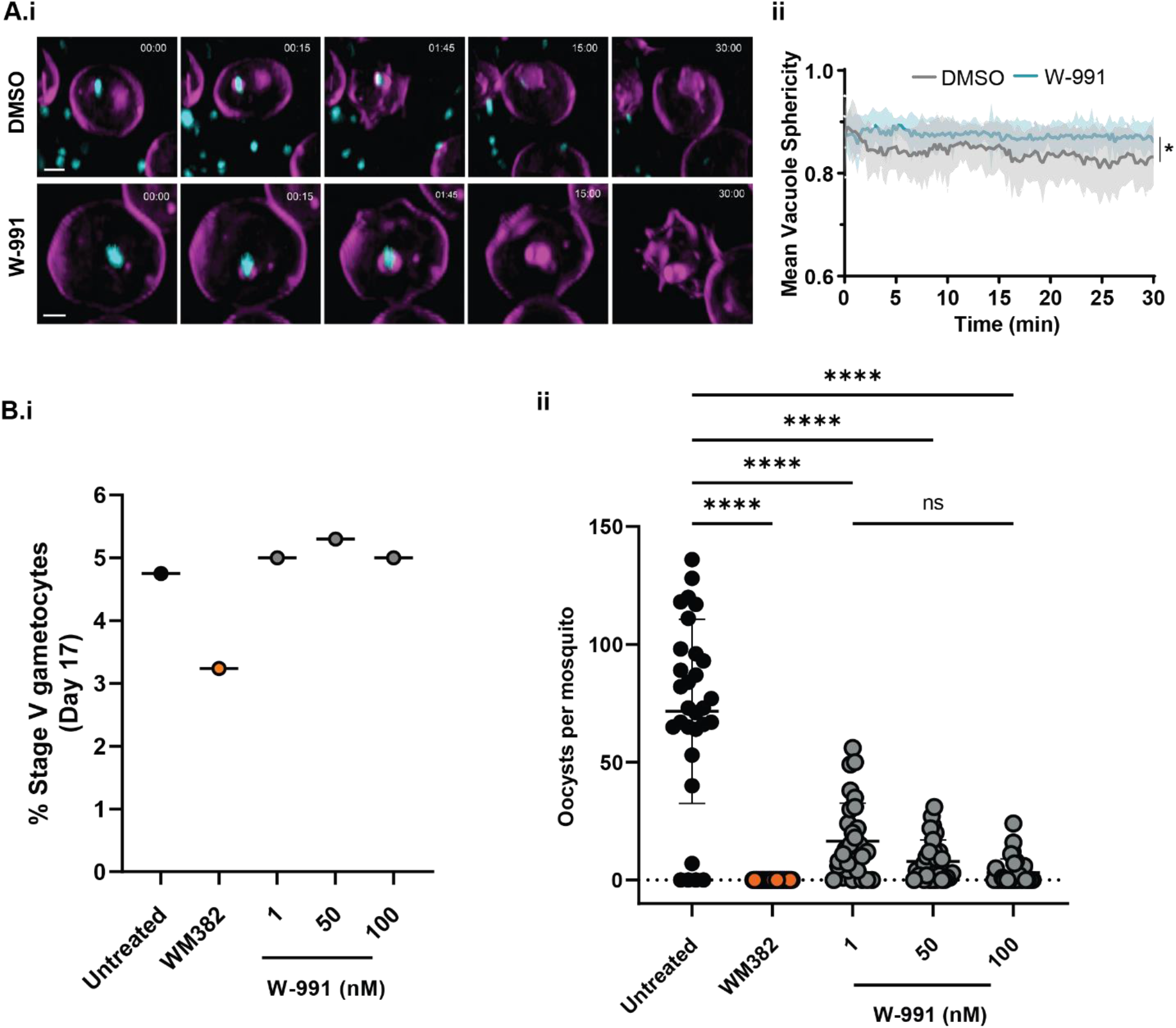
W-991 blocks merozoite transition to ameboid ring in RBCs, and blocks transmission in mosquitoes. **(A.i)** Representative images of 4D Lattice Light Sheet Microscopy of merozoite (cyan) directly after invasion of RBCs (all membranes are magenta). Treatment with W-991 shows disruption to the formation of the parasitophorous vacuole membrane (magenta) compared to vehicle control. **(ii)** The mean vacuole sphericity for W-991 and DMSO treatments across 30 min filming period. Statistical analysis was conducted via a Nested t-test (two-tailed) in GraphPad Prism showing p= 0.716 at T=0 mins and a p=0.045 at T=15 mins (*). Across three independent experiments, there were 20 and 22 events analysed for W-991 (60 nM) and DMSO (0.001%) treatments, respectively (Fig S8 and Movie S1 and S2). **(B.i)** Stage V gametocytes were quantified on day 17 which demonstrates there was no defect in gametocyte development upon W-991 treatment. **(ii)** Oocysts were quantified in mosquito midguts on day 7 after feeding which showed a dose-dependent decrease in oocysts per mosquito with W-991 treatment. Statistical tests (two-way ANOVA) were performed in GraphPad Prism. **** indicates *p*<0.0001. Positive control compound, WM832, was used at 50 nM (Favuzza et al., 2020).

### PfSTART1 inhibitors show no liver stage activity but block parasite transmission to mosquitoes

Since sporozoites are known to form a PV in hepatocytes after invasion (Mota et al., 2001), we evaluated W-991 against sporozoite invasion to determine if PfSTART1 may be required for this process. This revealed that W-991 had no impact against sporozoite invasion (Fig S8A, p>0.05).

Next, we assessed the activity of the M-833 series against the sexual stage of the *P. falciparum* lifecycle, in a *P. falciparum* dual gamete formation assay (DGFA) (Delves et al., 2018). Mature gametocytes were treated for 48 h with 1 µM M-833 or W-991 and then gametogenesis triggered. Exflagellation and surface Pfs25 expression were quantified, which demonstrated that no inhibition of either male or female gamete formation in the presence of M-833 and W-991 had occurred (Fig S8B). To further evaluate this series in the sexual stage we next performed a standard membrane feeding assay (SMFA) where 3 increasing concentrations of W-991 (1, 50, 100 nM) were exposed to stage IV-V gametocytes from days 14-17. On day 17 the percentage of stage V gametocytes were evaluated which showed there was no impact of W-991 affecting gametocyte development (Fig 6B.i). Media containing the compounds was then removed and treated gametocytes were fed to mosquitoes and the number of oocysts present in the midgut of mosquitoes was quantified 7 days later. This showed that W-991 exhibited transmission blocking activity of *P. falciparum* to mosquitoes in a dose-dependent manner, with 100 nM of W-991 reducing the number of oocysts by >20-fold compared to untreated control (Fig 6B.ii, *p*>0.001). Altogether, this indicates that the PfSTART1 inhibitors cause irreversible inhibition of an essential process in the transmission phase of the lifecycle between induction of gametocytogenesis and oocyst formation, thereby showing potential as a transmission blocking antimalarial.

### PfSTART1 expression, processing and localisation in schizonts and merozoites

We wanted to understand the function of the PfSTART1 protein in *P. falciparum* parasites to comprehend the mode of action of M-833. HA-tagged PfSTART1 SLI-WT parasites were tightly synchronised and proteins were extracted in very young rings, older rings, young trophozoites, older trophozoites and schizont stages (Fig 7A, Fig S9A). Following normalisation to the PfHSP70-1 loading control, PfSTART1 was most strongly expressed in schizonts and in very young rings (Fig 7A.ii). In addition to the main ∼60 kDa band (corresponding to PEXEL-cleaved PfSTART1-HA), a minor processed band (around 50 kDa) was also observed. The site of cleavage is not known, but must occur at the N-terminal, since this processed band was also detected with the HA-antibody. The anti-PfSTART1 antibody provided similar results to the anti-HA antibody, with limited off-target signals. Most of the subsequent experiments were therefore conducted on 3D7 parasites, in order to avoid potential effects linked to the additional HA tag on the C-terminus of PfSTART1.

**Figure 7.**
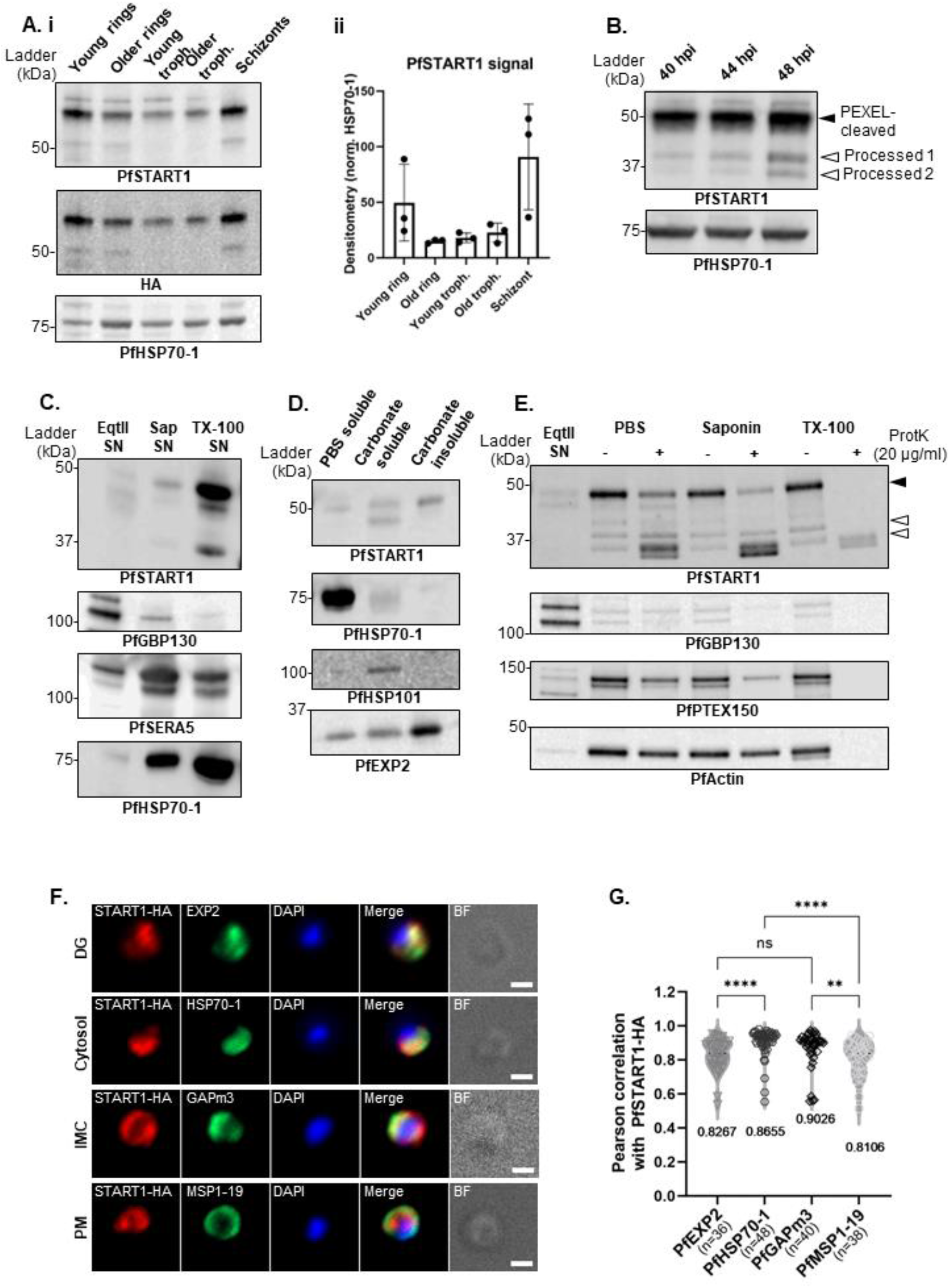
PfSTART1 expression, processing and localisation. **(A.i)** Western blot of SLI-WT parasites along the 48 h erythrocytic cycle. A saponin-lysis was performed on tightly synchronised SLI-WT young rings, older rings, young trophozoites, older trophozoites and schizonts. **(A.ii)** Densitometry of PfSTART1 over three biological replicates (normalised by the corresponding PfHSP70-1 signal). **(B)** Synchronous 3D7 trophozoites were magnet-purified, and whole cells harvested at ∼40 hpi (hour post invasion), ∼44 hpi and ∼48 hpi (this last sample was treated with 10 µM E64 to prevent egress). PEXEL-cleaved PfSTART1 and two further processed forms are indicated by solid and empty arrows respectively. **(C)** To localise PfSTART1, Percoll-purified 3D7 schizonts were sequentially lysed with equinatoxin II (Eqt II), saponin (Sap) and Triton-X100 (TX-100), and the supernatants (SN) were collected to harvest proteins localising in the red blood cell (RBC) cytosol, the parasitophorous vacuole (PV) and the parasite, respectively. PfGBP130 is a protein known to be exported into the RBC cytosol; PfSERA5 localises to the PV in schizonts; PfHSP70-1 is a parasite cytosolic protein. **(D)** To determine the solubility of PfSTART1, saponin-lysed 3D7 schizonts were sequentially lysed in PBS (with five freeze-thaw cycles) and sodium carbonate. The supernatant resulting from the PBS lysis was collected (“PBS soluble”), as well as the supernatant and the pellet of the carbonate lysis (“Carbonate soluble” and “Carbonate insoluble” respectively). Controls are the soluble protein PfHSP70-1, the membrane-associated PfHSP101, and the integral transmembrane protein PfEXP2. **(E)** Proteinase K protection assay was conducted on Percoll-purified 3D7 schizonts. Schizonts were first lysed in EqtII, the supernatant (SN) of which was collected. The remaining parasite and PV were either incubated in PBS (no lysis), saponin (PVM-lysis) or TX-100 (lysis of all membrane) with or without proteinase K. PfGBP130 is RBC cytosolic protein; PfPTEX150 is a PV protein; PfActin-1 is a parasite cytosolic protein. Note that another replicate for each of these experiments is shown in Fig S9E. **(F)** Representative immunofluorescent assay (IFA) images are shown for SLI-WT merozoites probed with antibodies against: HA (red; PfSTART-HA), PfEXP2 (green; dense granules (DG)), PfHSP70-1 (green; parasite cytosol), PfGAPm3 (green; inner membrane complex (IMC)), or PfMSP1-19 (green; merozoite plasma membrane (PM)). Nuclei were stained with DAPI (in blue). Scale bar is 1 μm. **(G)** Pearson correlation coefficient was calculated on individual merozoites between START-HA and PfEXP2 (DG), PfGAPm3 (IMC), PfHSP70-1 (cytosol) and PfMSP1-19 (PM). The number of cells analysed and the average r values are shown on the scatter plot. Kruskal-Wallis with Dunn’s multiple comparisons test. ns: p≥0.05. **: p<0.005. ****: p<0.0001.

To further understand the processing of PfSTART1, proteins of tightly synchronised 3D7 schizonts were harvested at 4 h intervals (E64 was used for the latest timepoint to prevent parasite egress; Fig 7B). This time course demonstrates that while the majority of PfSTART1 remains as a ∼48 kDa band (PEXEL-cleaved), two further processing events occur as schizogony progresses (PfSTARTproc1 ∼39 kDa, PfSTARTproc2 ∼36 kDa). Both processing of PfSTART1 occurred at the N-terminus, as both processed bands could be detected by an HA-antibody on SLI-WT schizonts (Fig S9B.i). To more precisely pinpoint when PfSTART1 was processed along schizogony, we treated schizonts with inhibitors stalling egress at different time points (Fig S9B.ii and iii): 49c is a plasmepsin X (PMX) inhibitor, which prevents activation of SUB1, early in the egress pathway (Pino et al., 2017); compound 1 (C1) is a protein kinase G inhibitor, which prevents exoneme discharge (Gurnett et al., 2002); E64 is a cysteine protease inhibitor which prevents the rupture of the red blood cell membrane, near the very end of egress (Bailly et al., 1992). The densitometry of processed bands was normalised by the PEXEL-cleaved PfSTART1, and the effect of treatment was analysed (Fig S9B.iii). 49c-treated schizonts contained the least amount of PfSTART1proc1 (compared to DMSO and other treatments), and C1- treated schizonts had the lowest levels of PfSTART1proc2. E64-treated schizonts contained the most of both processed bands, with a significant enrichment in PfSTART1proc2 compared to the DMSO control.

PfSTART1 contains an unusual PEXEL motif and was recently described as not being exported to the RBC in trophozoites (Ressurreição et al., 2023). To investigate PfSTART1 localisation in schizonts, 3D7 schizonts were Percoll-purified and sequentially lysed in equinatoxin II (EqtII), saponin (Sap) and Triton X100 (TX100) in order to collect the supernatant (SN) corresponding to the RBC cytosol soluble fraction, the parasitophorous vacuole (PV) soluble fraction, and the parasite fraction, respectively (Fig 7C and Fig S9C). PfGBP130, a known exported soluble protein (Jonsdottir et al., 2023; Perkins, 1988), was detected in the EqtII SN; PfSERA5, a soluble protein which localises in the PV of schizonts (Blackman, 2008; Jonsdottir et al., 2023), was enriched in the Sap SN; parasite cytosolic protein PfHSP70-1 was found mainly in the TX-100 SN. PfSTART1 was almost only detected in the TX100 fraction, indicating that it was either within the parasite, and/or associated with membranes.

We tested whether PfSTART1 was fully soluble (in which case, PfSTART1 should be localised within the parasite), membrane-associated or membrane-bound. To do so, saponin-lysed 3D7 schizonts were sequentially lysed: first in PBS (with freeze-thaw cycles to break the cells), the supernatant of which was described as “PBS soluble”. The pellet was then incubated in sodium carbonate: the supernatant was described as “carbonate soluble” (proteins associated with membranes) and the pellet as “carbonate insoluble” (integral membrane proteins) (Fig 7D and Fig S9D). Controls included PfHSP70-1, PfHSP101 and PfEXP2, which were detected in the PBS soluble fraction, in the carbonate soluble fraction, and the carbonate insoluble fractions respectively. PfSTART1 was mainly detected in the carbonate soluble and insoluble fractions in accordance with previous findings (van Ooij et al., 2013).

We then performed a proteinase K protection assay to determine PfSTART1 localisation: different membranes were lysed (the RBC membrane with EqtII, the PVM with saponin, and all membranes with TX100) with or without proteinase K (Fig 7E, Fig S9E). If a protein was protected from proteinase K degradation by the presence of a membrane, this protein would be localised within the bounds of this membrane. PfGBP130 was localised in the EqtII supernatant as expected. Parasite PfActin-1 was protected from proteinase K degradation unless all the membranes were lysed. PVM-associated PfPTEX150 was mainly degraded upon addition of saponin, and was further degraded after addition of TX100, although some degradation was already observed when only the RBC membrane was lysed. A similar pattern was observed for PfSTART1, suggesting that the PEXEL-cleaved protein is also mainly localised in the PV, whilst PfSTART1proc1 and PfSTART1 proc2 appear to be inside the parasite and protected from proteinase K cleavage.

We conducted a series of immunofluorescence assays (IFAs) to localise PfSTART1 in schizonts, merozoites and rings (Fig 7F and S10). We used markers of invasion-specific secretion organelles (micronemes, dense granules, rhoptries), of the endoplasmic reticulum, of the parasite cytoplasm, inner-membrane complex, parasite plasma membrane and RBC membrane. From these assays, it appears that PfSTART1 is not restricted to a specific secretory organelle. It appears to be distributed widely throughout the merozoite, within its plasma membrane, with a particularly good co-localisation with HSP70-1 (cytosol), GAPm3 (inner membrane complex) and EXP2 (dense granules), as measured by Pearson correlation coefficient (r) (Fig 7G). In young rings, it seems likely PfSTART1 remains within the parasite or in the PV (Fig S10C).

## Discussion

In this study, we explored the activity of an anti-*Plasmodium* compound series whereby parasites resistant to the hit compound M-833 were generated. Whole genome sequencing revealed point mutations in PfSTART1 (N309K, N330K, I224F). Through structure activity relationship studies, we achieved analogues that remained on-target for PfSTART1, whilst significantly increasing the potency to achieve an EC50 against wildtype parasites in the low nanomolar range. Engineering the resistance mutations N309K and N330K into drug sensitive parasites conferred resistance to M-833 and to the potent analogue W-991. Furthermore, knocking down PfSTART1 sensitised parasites to both M- 833 and W-991. We demonstrated direct binding of M-833 and analogues to PfSTART1, both using recombinant proteins, as well as parasite lysate. Overall, this demonstrates that M-833 and analogues exert their antiparasitic activity via the inhibition of PfSTART1.

Mammalian START proteins are known to have pathological roles in several human diseases (*e.g.*, STARD3 overexpression in cancer (Asif et al., 2021)), including infectious diseases (*e.g.*, human STARD11 is hijacked by intracellular *Chlamydia trachomatis* bacteria (Derré et al., 2011)). Due to this, several inhibitors of human START proteins have been identified: STARD1 inhibitors C1-C6 (Akula et al., 2012); STARD11 inhibitors HPA-12 (Yasuda et al., 2001) and SC1 plus analogues (Nakao et al., 2019); STARD3 (predicted) inhibitor D(-)-tartaric acid (Chitrala and Yeguvapalli, 2014) and STARD3 inhibitor VS1 (low potency) (Lapillo et al., 2019). The M-833 series does not show structural similarity to the aforementioned human START inhibitors and considering that the M-833 series had CC50 > 29 µM against HepG2 cells (a cell line known to express START proteins (Garbarino et al., 2012)), this indicates that our compounds do not target human START proteins. Follow up binding experiments with the M-833 series and human START proteins are required to confirm this. Additionally, Solvent PISA profiling suggests target specificity of M-833 series for PfSTART over other *P. falciparum* START proteins (PF3D7_1351000 and PF3D7_1463500) detected in the assay.

Amongst the 5 START domain containing proteins in *P. falciparum*, PfSTART1 (PFA0210c) is the most well-characterised (Hill et al., 2016; Ressurreição et al., 2023; van Ooij et al., 2013). Solvent PISA profiling described in this study represents novel unbiassed strategy for antimalarial drug-target identification. In addition to representing means of orthogonal protein-engagement validation for target-candidates derived from parallel approaches, such as TPP, it enables detection of drug interactions with proteins not susceptible to temperature-induced aggregation. PfSTART1 represents one of such highly thermostable protein (Dziekan et al., 2019), and its interaction with the drug in parasite lysate cannot be identified based on stability shift <75°C.

Further, the Solvent PISA strategy introduced here involves major improvements over mass spectrometry-based target identification workflows established for *Plasmodium* (Bopp et al., 2023; Dziekan et al., 2020; Dziekan et al., 2019; Favuzza et al., 2020). Incorporation of Proteome Integral Solubility Alteration (PISA) assay format (Zhang et al., 2022) and Data Independent Acquisition mass spectrometry (DIA-MS) (Doerr, 2015) into the solvent assays significantly reduces the required MS-analysis time and profiling cost, as well reducing sampling bias by profiling the entire proteome across the solvent gradient. Due to the sensitive nature of this assay, we were able to identify three other proteins (PF3D7_1035900, PF3D7_1149200, PF3D7_1216300) other than PfSTART1 in the *P. falciparum* proteome that were significantly stabilised upon W-991 treatment. This will allow follow-up studies to determine if the PfSTART1 inhibitors have additional modes of action that are linked to these proteins.

It was initially encouraging to observe that a short 4 h treatment stopped parasite development for three days post compound removal. However, a longer follow-up of M- 833 and W-991 treated (and removed) parasites indicated that they could recover over a longer period, which was confirmed by the parasite reduction ratio assay. Due to the essentiality of PfSTART1 in the blood stage (Hill et al., 2016), it was somewhat unexpected that parasites could recover after being subjected to 10xEC50 of W-991 for up to five days. One possible explanation of this is that the EC50 value derived from a standard 72 h growth assay could underestimate the concentration required to effectively kill parasites due it being unable to differentiate between failed merozoite conversion into rings or dead parasites. It is therefore plausible that conducting a longer growth assay (>72 h) would assist in determining a dose that would cause irreversible death. Alternatively, it is also possible that inhibiting PfSTART1 does not lead to death but rather the parasites enter a dormant mode in which they can survive until compound levels have decreased. This is reminiscent of artemisinin resistant parasites whereby prolonged ring-stage has been associated with treatment survival (Teuscher et al., 2010; Witkowski et al., 2010). Interestingly, PfSTART1 has been recently found to be significantly upregulated in both lab-adapted and field-derived artemisinin resistant strains that contain mutations in Kelch13 (Dd2^C580Y^ and Cam3.II^R539T^) (Mok et al., 2021). Additionally, artemisinin resistance has also recently been shown to be present in male gametocyte activation whereby parasites resistant to artemisinin are activated under artemisinin treatment, whilst sensitive parasites remain inactivated (Witmer et al., 2020). It would, therefore, be interesting to conduct combination experiments with our PfSTART1 inhibitors and artemisinin against artemisinin resistant strains in both the asexual blood stage and mosquito stages to determine if inhibiting PfSTART1 could sensitise resistant parasites to artemisinin. This could be an attractive property of PfSTART1 inhibitors and is an avenue to be explored for drug development potential.

Beyond its consideration as a potential drug target, another clinically relevant aspect of PfSTART1 is as a vaccine candidate. Indeed, antibodies to the *P. vivax* StAR-related lipid transfer protein (PVX_081550) (∼50% sequence identity to PfSTART1), have been found and associated with clinical protection in a cohort of children in Papua New Guinea (Mazhari et al., 2023) and has been suggested as a possible vaccine candidate (Müller-Sienerth et al., 2020; Zenonos et al., 2014). It will be interesting to assess the invasion (and ring conversion) inhibition of PfSTART1 antibodies and nanobodies (Dietrich et al., 2022).

The activity profile of the PfSTART1 inhibitors was found to be variable in other stages of the lifecycle outside the blood stage with no drug effect against liver stage invasion, stage V gametocyte development and gametogenesis. The lack of activity against the liver stage of infection was surprising given that sporozoites grow and replicate in a PV, similar to the asexual blood stages (Meis et al., 1983; Risco-Castillo et al., 2015). The apparent inactivity of the PfSTART1 inhibitors could be due to the experimental design: indeed, sporozoites and W-991 were added to human liver cells and evaluated only 2 h post invasion. It is possible we did not capture the developmental inhibition of the PV formation. Further experiments where sporozoites are added to liver cells with the compounds for a longer period are required to definitively rule out liver activity of this series.

We found that PfSTART1 inhibitors were active in a standard membrane feeding assay. Whilst the functionality of PfSTART1 has been previously explored in the asexual blood stage (Hill et al., 2016; Margarida et al., 2023; van Ooij et al., 2013), there is little known about PfSTART1 in the mosquito stage of infection. Transcriptomic studies have shown that PfSTART1 is expressed in salivary gland sporozoites (Gómez-Díaz et al., 2017; Lindner et al., 2019) and ookinetes (López-Barragán et al., 2011). Since there is no PV formation in ookinetes (Garnham et al., 1962; Meis and Ponnudurai, 1987; Zieler and Dvorak, 2000), it is possible that PfSTART1 is utilised for another lipid dependent process during this stage. Sporozoites, however, are known to form a PV during the invagination of the epithelial cell membrane in the mosquito salivary glands (Ghosh and Jacobs-Lorena, 2009; Pimenta et al., 1994; Rodriguez and Hernández-Hernández, 2004) which could suggest that the PfSTART1 inhibitors could act through similar mode of action to that we see in the asexual blood stage but further investigations are required to confirm this.

To study the function of PfSTART1 in the blood stage, we tagged the C-terminal region of the protein with a 3xHA using the selection linked integration (SLI) system. Unexpectedly, we saw an increased sensitivity of the SLI-WT parasites which did not occur in the CRISPR WT parasites that did not include a tag. This may be due to the addition of the 3x HA and P2A peptide (additional 55 amino acids) to PfSTART1, which could be interfering with its normal functioning and thereby sensitising the SLI-WT parasites to M-833. The C-terminus of PfSTART1 has been shown to be important for the regulation of lipid transfer (Hill et al., 2016), with the last 30 amino acids of *Plasmodium* START orthologues more conserved than the upstream spacer region. The unusual C-terminal extension is absent in most mammalian START proteins and is predicted to form an alpha helix (Alphafold) (Jumper et al., 2021; Varadi et al., 2022).

Consistent with previous reports, we found that PfSTART1 is predominantly expressed in schizonts in the asexual blood stage, is membrane-associated and is not exported into the surrounding RBC (despite its cleaved PEXEL motif) (Ressurreição et al., 2023; van Ooij et al., 2013). The close association with membranes is probably mediated by one or more interacting partner(s), considering that PfSTART1 does not contain a transmembrane domain. PfSTART1 localises partly to the PV and partly within the parasites in schizonts and has a broad distribution within merozoites (not colocalising clearly with any specific organelle). This suggests that PfSTART1 may have several functions, transferring lipids in several locations.

PfSTART1 appears to be processed multiple times at its N-terminus. Ressurreição and colleagues demonstrated that the PEXEL motif of PfSTART1 is cleaved by plasmepsin V, but that PfSTART1 is not exported (Ressurreição et al., 2023). Therefore, the main PfSTART1 signal observed on western blots at ∼48 kDa likely corresponds to the PEXEL-cleaved protein. Of note, using protein molecular mass calculator, the full-length PfSTART1 sequence is predicted to be 54 kDa, 51 kDa once the signal sequence is removed, and 46 kDa once the PEXEL motif is cleaved (close to the ∼48 kDa band observed). In addition, we demonstrated that two further processing events occur as schizogony progresses. First, the processing to PfSTART1proc1 (∼39 kDa) seems to occur early in schizogony, shortly after SUB1 activation by plasmepsin X, because plasmepsin X inhibition reduced the levels of PfSTART1proc1. Interestingly, PfSTART1 contains a plasmepsin X cleavage site (SDIQ (Mukherjee et al., 2022)), although the resulting theoretical products should be 20 kDa and 26 kDa (each far from the 39 kDa or 36 kDa bands observed). No SUB1 or plasmepsin IX cleavage sites were identified in PfSTART1 (Favuzza et al., 2020; Lidumniece et al., 2022). The second processing event to PfSTARTproc2 (∼36 kDa) appears to occur after protein kinase G activation (because inhibiting protein kinase G reduced the levels of PfSTARTproc2). PfSTARTproc2 appears to accumulate considerably in the presence of cysteine inhibitor E64 (which prevents the rupture of the RBC membrane and the release of merozoites). Overall, it is possible that these processed forms of PfSTART1 are important in merozoites for the upcoming invasion and establishment of the PVM. Both PfSTART1proc1 and PfSTART1proc2 show complete resistance to proteinase K after saponin lysis, indicating they are not located within the PV like the PEXEL-cleaved protein. However, both PfSTART1proc1 and PfSTART1proc2 only represent a small proportion of the total PfSTART1 (less than 40% of the PEXEL-cleaved form). Therefore, the functional relevance of these processed PfSTART1 remains to be investigated.

Inhibition of PfSTART1 in the blood stage led to the abnormal development of the membranes surrounding the merozoite directly after invasion as visualised with lattice light sheet microscopy. Whilst this phenotype was distinct, it remains unknown as to the composition of these lipids, in addition to the direction and location of transport. In *Plasmodium,* the molecular mechanisms underpinning PVM formation are still unknown (Goldberg and Zimmerberg, 2020; Spielmann et al., 2012) but it has recently been shown that the PVM is comprised of mostly host red blood cell lipids, rather than parasite derived material (Geoghegan et al., 2021). Overall, as previously proposed (Hill et al., 2016), our data supports the model whereby PfSTART1 aids in expanding the PVM using lipids from the parasite plasma membrane (Fig 8). How PfSTART1 is delivered to the PVM upon invasion remains to be uncovered. A simple explanation would be that it resides in secretory organelles and is released to the developing PV space upon invasion (Fig 8), similarly to dense granule protein, EXP2. Whilst we saw some co-localisation between EXP2 and PfSTART1 in merozoites, it was not confined to a specific organelle, in accordance with previous studies (Ressurreição et al., 2023). Indeed, we also saw considerable overlap between PfSTART1 and the parasite cytosol and the inner membrane complex. Therefore, in addition to expanding the PVM, PfSTART1 could also play a role in moving lipids within the parasite: for instance, PfSTART1 might assist in the rapid disassembly of the inner membrane complex, therefore providing lipids for the PPM and PVM expansion. Further studies utilising expansion microscopy and lattice light sheet imaging of fluorescently tagged PfSTART1 would help to resolve the localisation of PfSTART1 during and directly following invasion. Coupling these experiments with the now confirmed PfSTART1 inhibitors would shed further light on the mechanism of action of this novel antimalarial series and assist in probing the function of this lipid transferase in *Plasmodium*.

**Figure 8.**
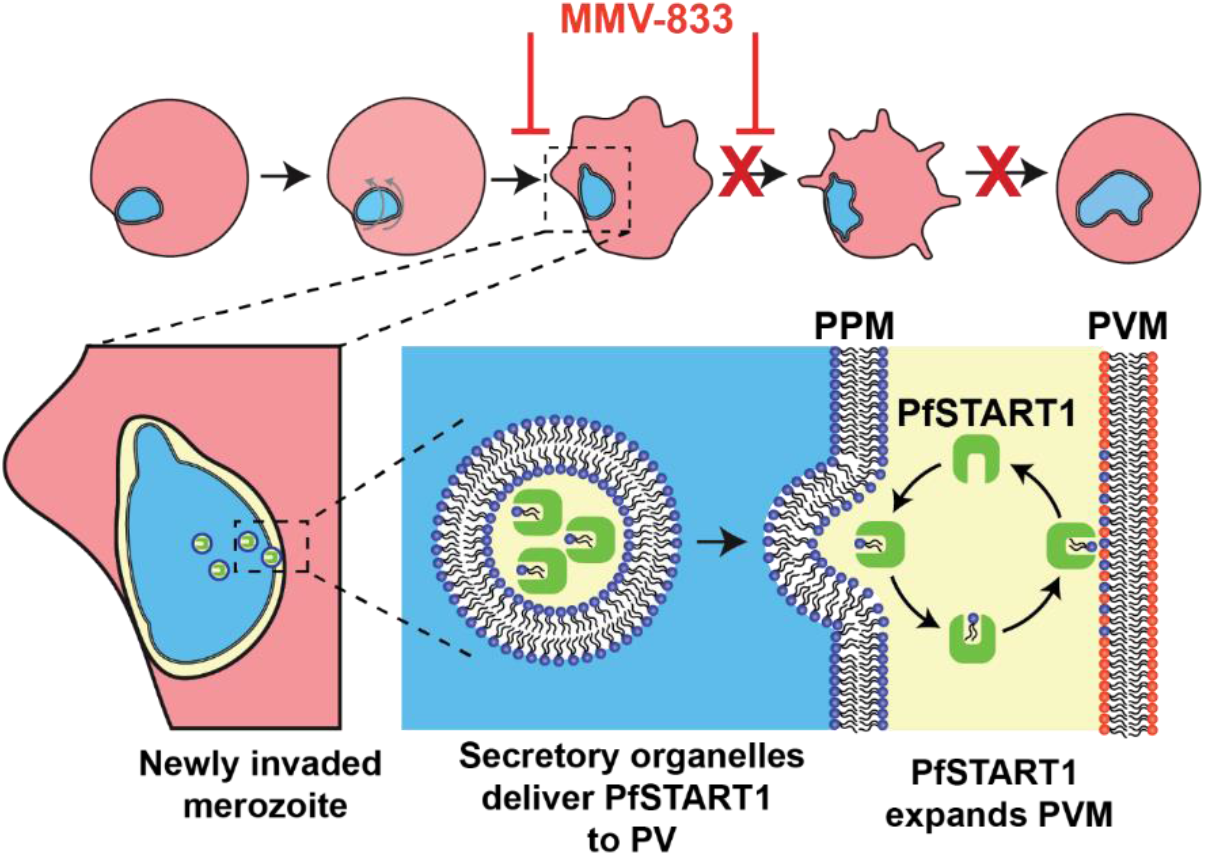
The M-833 series blocks conversion of newly invaded merozoites into ring-stage parasites by inhibiting the PfSTART1 protein and remodeling of the parasitophorous vacuole membrane (PVM). **(Top)** After invasion, the merozoite begins to spin and produce pseudopodia before differentiating into an amoeboid ring-stage parasite. (Bottom) After invasion PfSTART1, is delivered into the vacuole space possibly via secretory organelles fusing with the parasite plasma membrane (PPM). Here PfSTART1 could transfer phospholipids from the PPM to the PVM to allow the latter to expand to accommodate the growing ring-stage parasite. The M-833 compound series tightly binds to PfSTART1, probably displacing the phospholipids and thereby preventing the expansion of the PVM and arresting parasite growth.

## Materials and Methods

### *P. falciparum* culture

*Plasmodium falciparum* 3D7 parasites were cultured as described previously (Trager and Jensen, 1976). They were grown in human red blood cells provided by the Australian Red Cross Blood Bank, at 4% haematocrit, maintained at 37°C in a special gas mixture (1% O2, 5% CO2, 94% N2). The medium used was complete RPMI medium: RPMI-1640 (Sigma), 25 mM HEPES (GIBCO), 0.37 mM hypoxanthine (Sigma), 31.25 μg/ml gentamicin (GIBCO), 0.2% NaHCO3 (Thermo Scientific), 0.5% AlbuMAX II (GIBCO).

### Generating resistance

Resistant parasites were generated as previously described (Dans et al., 2023). A clonal population of 3D7 parasites (10^8^ parasites in five replicates, A to E) were exposed to 10 x EC50 = 3 μM of MMV006833 (M-833), until most parasites died. The drug was then removed, and parasites were allowed to recover. Another cycle of drug treatment (3 μM M-833) was then resumed. The resistant lines were cloned out by limiting dilution (diluting the culture into a 96-well plate to achieve an average of ∼0.3 parasite per well), prior to growth inhibition assays and genomic DNA (gDNA) extraction (DNeasy Blood and Tissue kit (Qiagen).

### Whole genome sequencing and genome reconstruction

Sequencing of M833- resistant parasites and parental 3D7 line was performed as previously described (Dans et al., 2020). Genomic sequencing data is available from European Nucleotide Archive; accession number PRJEB65444.

### Molecular biology and transfection of *P. falciparum*: Selection Linked Integration method

To introduce the PfSTART1 mutations, the second half of the coding sequence of PfSTART1 (PF3D7_0104200), was recodonised without introns to the bias of *Saccharomyces cerevisiae* containing either WT, N309K or N330K mutations, and synthesised as gBlock fragments (Integrated DNA Technologies). The first 819 bp of the 5′ native sequence of PfSTART1 was amplified from gDNA, to form the 5’ homology block (5’HB). The 3′ recodonised gBlock fragment comprising of the last 552 bp of either the WT, N309K or N330K PfSTART1 to form the ‘recodon’ block (primers listed in Table S3). The 5’HB and recodon products were sewn together by overlapping PCR to reconstitute the full coding sequence of PfSTART1. These blocks (WT/N309K/N330K) were digested with *Bgl*II/*Pst*I, and inserted into the parasite vector p-HA-2A-Neo-glmS (Jonsdottir et al., 2021). This plasmid contains an in-frame haemagglutinin (HA) epitope for protein identification and a P2A peptide (2A) to skip continuous peptide bond synthesis of a downstream neomycin resistance gene (NeoR).

Parasites containing the transfected DNA were selected using 2.5 nM WR99210 (Jacobus). To select for parasites in which single crossover integration had occurred into the *pfstart1* locus, parasites were treated with 400 μg/mL G418 (geniticin, Sigma Aldrich) for 14 days: only parasites that have integrated the construct into their genome expressed the neomycin resistance gene together with the PfSTART1 protein. Parasite lines were cloned using limiting dilution. These clones were then tested on Western Blot (for correct HA-tagging and protein size), and correct integration and sequence were tested by PCR analysis: PCR on the gDNA was conducted using the primers START_5’UTR.F and START_Seq.R (Table S3), and sequencing was performed at the Micromon Genomics platform of Monash University.

### Molecular biology and transfection of *P. falciparum*: CRISPR method

The full 5’ homology region of PfSTART1 (either WT, N309K or N330K) was amplified from pPfSTART1-HA-2A-Neo-glmS with the same 5’ *Bgl*II primer and a new 3’ primer that replaced with *Pst*I with *Spe*I and introduced a stop codon. The PCR product was then digested with *Bgl*II and *Spe*I and inserted into similarly digested p1.2 parasite vector (Dans et al., 2023). The p1.2 vector had previously been inserted with a homology region amplified from the 3’ end of the *pfstart1* gene (3’HB) to form a second homology block for double recombination. The primers used to amplify 3’HB are listed primers listed in Table S3.

60 μg of p1.2-PfSTART1(WT/N309K/N330K) plasmid was linearised with *Kas*I, precipitated in 3 M sodium acetate and ethanol (70% final concentration) overnight at - 20°C. The DNA pellet was washed in cold 70% ethanol and resuspended in cytomix. 50 μM of crRNA (Table S3) and 50 μM of tracRNA (Integrated DNA Technologies) were allowed to form a duplex (5 min at 95°C), and Cas9 enzyme (Integrated DNA Technologies) was added, and incubated for 20 min at room temperature. This crRNA- tracRNA-Cas9 mix was added to each DNA preparation and transfected to 3D7 ring parasites using the Bio-Rad Gene Pulser at 310 V and 950 μF.

Parasites containing the transfected DNA were selected using 2.5 nM WR99210 (Jacobus). Parasites lines were cloned using limiting dilution. Correct integration was tested by PCR (using primer START_5’UTR.F and START_Int.R1, Table S3), and sequencing was done using the primers START_3F and START_Int.R2 (Table S3).

### General Chemistry Methods

NMR spectra were recorded on a Bruker Ascend^TM^ 300. Chemical shifts are reported in ppm on the δ scale and referenced to the appropriate solvent peak. Acetone-d6, and CDCl3 contain H2O. Chromatography was performed with silica gel 60 (particle size 0.040-0.063 µm) using an automated CombiFlash Rf Purification System. LCMS were recorded on an Agilent LCMS system comprised of an Agilent G6120B Mass Detector, 1260 Infinity G1312B Binary pump, 1260 Infinity G1367E HiPALS autosampler and 1260 Infinity G4212B Diode Array Detector (Method B). Conditions for LCMS Method A were as follows, column: Luna® Omega 3 µm PS C18 100 Å, LC Column 50 × 2.1 mm at 20 °C, injection volume 2 µL, gradient: 5-100% B over 3 min (solvent A: H2O 0.1% formic acid; solvent B: ACN 0.1% formic acid), flow rate: 1.5 mL/min, detection: 100-600 nm, acquisition time: 4.3 min. Conditions for LCMS Method B were as follows, column: Poroshell 120 EC-C18, 2.1 × 30 mm 2.7 Micron at 30 °C, injection volume 2 µL, gradient: 5-100% B over 3 min (solvent A: H2O 0.1% formic acid; solvent B: ACN 0.1% formic acid), flow rate: 0.8 mL/min, detection: 254 nm, acquisition time: 4.1 min. Unless otherwise noted, all compounds were found to be >95% pure by this method. HRMS were acquired through The Bio21 Mass Spectrometry and Proteomics Facility using a Thermo Scientific™ nano-LC Q Exactive™ Plus Mass spectrometer.

### Chemistry procedures

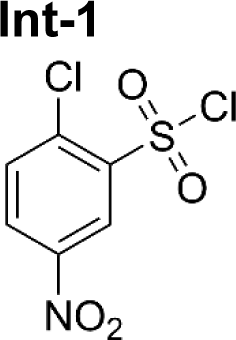

#### *2-Chloro-5-nitro-benzenesulfonyl chloride* (Int-1)

1-Chloronitrobenzene (1 g, 6.3 mmol) was dissolved in neat chlorosulfonic acid (4.2 mL, 63.4 mmol) and stirred at 120°C for 48 h. The mixture was then slowly pipetted into ice water (20 mL). The resulting precipitate was then collected by vacuum filtration to afford **Int-1** as a brown solid (1.1 g, 68%). ^1^H NMR (300 MHz, CDCl3): δ 9.02 (d, *J* 2.6 Hz, 1H) 8.52 (dd, *J* 8.7, 2.6 Hz, 1H) 7.89 (d, *J* 8.8 Hz, 1H).

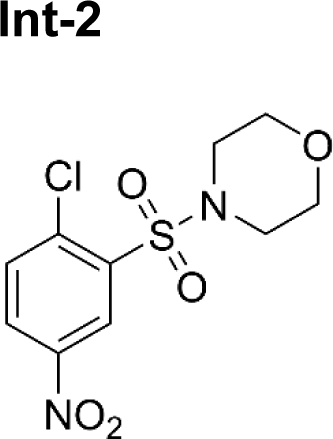

#### *4-(2-Chloro-5-nitro-phenyl)sulfonylmorpholine* (Int-2)

Morpholine (0.405 mL, 4.69 mmol) was added dropwise to a stirred solution of triethylamine (1.09 mL, 7.81 mmol) and **Int-1** (1.0 g, 3.9 mmol) in DCM (20 mL) at 0°C. The reaction was stirred at this temperature for 3 h. The reaction was then washed with 2M HCl (3 × 20 mL) and the organics dried with anhydrous Na2SO4, filtered and concentrated. The crude was then purified by column chromatography eluting with 100% CyHex to 50% EtOAc/CyHex to obtain **Int-2** as a solid (816 mg, 68%). ^1^H NMR (300 MHz, CDCl3): δ 8.90 (d, *J* 2.7 Hz, 1 H), 8.36 (dd, *J* 8.7, 2.7 Hz, 1 H), 7.76 (d, *J* 8.7 Hz, 1 H), 3.71 - 3.79 (m, 4H), 3.32 - 3.43 (m, 4H). LCMS *m/z* 307.2 [M+1].

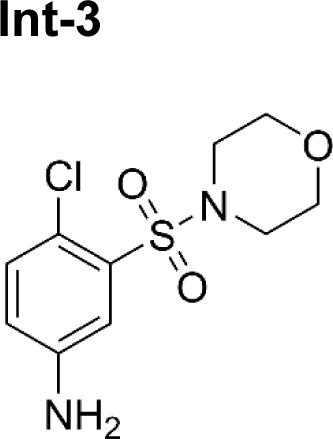

#### *4-Chloro-3-morpholinosulfonyl-aniline* (Int-3)

To a solution of **Int-2** (816 mg, 2.66 mmol) in EtOH (5 mL) and saturated NH4Cl (5 mL) added zinc (1.7 g, 27 mmol). The mixture was then stirred at 40°C for 3 h. The mixture was then filtered through diatomaceous earth which was washed with EtOH (3 × 10 mL). The filtrate was concentrated, and the resulting residue was taken up into EtOAc (30 mL) and washed successively with saturated NaHCO3 (20 mL), H2O (20 mL) and brine (20 mL). The organic layer was then dried over anhydrous Na2SO4, filtered and concentrated to afford **Int-3** as a solid (480 mg, 65%). ^1^H NMR (300 MHz, CDCl3): δ 7.35 (d, *J* 2.9 Hz, 1H) 7.32 – 7.27 (m, 1H) 6.79 (dd, *J* 8.5, 2.8 Hz, 1H), 3.71 - 3.78 (m, 4H) 3.26 - 3.36 (m, 4H). LCMS *m/z* 277.2 [M+1].

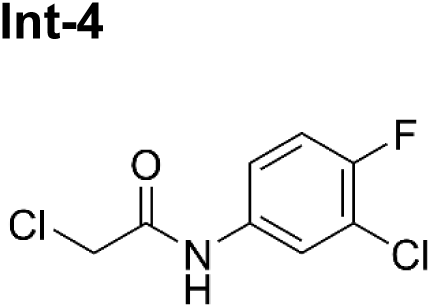

#### *2-Chloro-N-(3-chloro-4-fluoro-phenyl)acetamide* (Int-4)

2-Chloroacetyl chloride (0.52 mL, 6.6 mmol) was added slowly to a stirred solution of 3-chloro-4-fluoro-aniline (800 mg, 5.5 mmol) and triethylamine (1.5 mL, 11 mmol) in DCM (20 mL) at 20°C. The reaction was then stirred for 16 h at this temperature. The organics were successively washed with 2M HCl (20 ml), saturated NaHCO3 (20 ml) and H2O (20 ml). The organic layer was dried with Na2SO4 and concentrated to afford **Int-4** as a solid (970 mg, 80%). ^1^H NMR (300 MHz, CDCl3): δ 8.23 (br s, 1 H), 7.75 (dd, *J* 6.5, 2.6 Hz, 1H), 7.39 (ddd, *J* 9.0, 4.0, 2.7 Hz, 1H), 7.14 (t, *J* 8.7 Hz, 1H), 4.21 (s, 2H). LCMS *m/z* 222.2 [M+1].

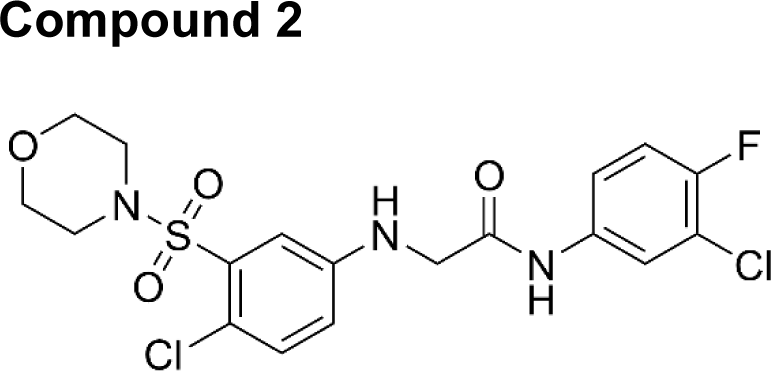

#### *N-(3-Chloro-4-fluoro-phenyl)-2-(4-chloro-3-morpholinosulfonyl-anilino)acetamide* (WEHI-985)

A mixture of **Int-4** (16 mg, 0.072 mmol) in DMF (1 mL) was prepared at 20°C. Potassium iodide (26 mg, 0.16 mmol) was added, and the reaction heated at 45°C for 1 h. The mixture was then cooled to 20°C and **Int-3** (19.9 mg, 0.072 mmol) added in one portion and the mixture heated again to 45°C in a sealed tube for 17 h. The mixture was then diluted with EtOAc (10 mL) and washed with 5% LiCl aq. (3 × 10 mL) and brine (10 mL). The organic layer was dried over anhydrous Na2SO4 and concentrated. The crude material was then purified by reverse phase preparatory HPLC using a gradient of 95% H2O/ACN to 100% ACN to obtain **WEHI-1883985** as a solid (12 mg, 36%). ^1^H NMR (300 MHz, CDCl3): δ 8.23 (s, 1 H), 7.71 (dd, *J* 6.5, 2.7 Hz, 1 H), 7.32 - 7.40 (m, 3 H), 7.10 (t, *J* 8.7 Hz, 1 H), 6.76 (dd, *J* 8.7, 2.92 Hz, 1 H), 4.78 (br s, 1 H), 3.96 (d, *J* 5.1 Hz, 2 H), 3.66 - 3.78 (m, 4H), 3.20 - 3.32 (m, 4H). LCMS *m/z* 462.2 [M+1]. HRMS acquired: (M + H) 462.0454; C18H18Cl2FN3O4S requires (M + H), 462.0452.

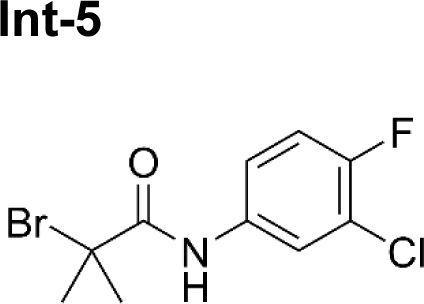

#### *2-Bromo-N-(3-chloro-4-fluoro-phenyl)-2-methyl-propanamide* (Int-5)

2-Bromoisobutyryl bromide (0.64 mL, 5.2 mmol) was added slowly to a stirred solution of 3-chloro-4-fluoro-aniline (500 mg, 3.4 mmol) and triethylamine (0.96 mL, 6.9 mmol) in DCM (15 mL) at 20°C. The reaction was then stirred for 16 h. The reaction mixture was washed with 2M HCl (20 mL), saturated NaHCO3 (20 mL) and H2O (20 mL). The organic layer was dried with anhydrous Na2SO4 and concentrated to afford **Int-5** as a solid (1.0 g, 99%). ^1^H NMR (300 MHz, CDCl3): δ 8.44 (br s, 1H), 7.75 (dd, J 6.5, 2.7 Hz, 1H), 7.32 - 7.40 (m, 1H), 7.13 (t, *J* 8.8 Hz, 1H), 2.05 (s, 6H).

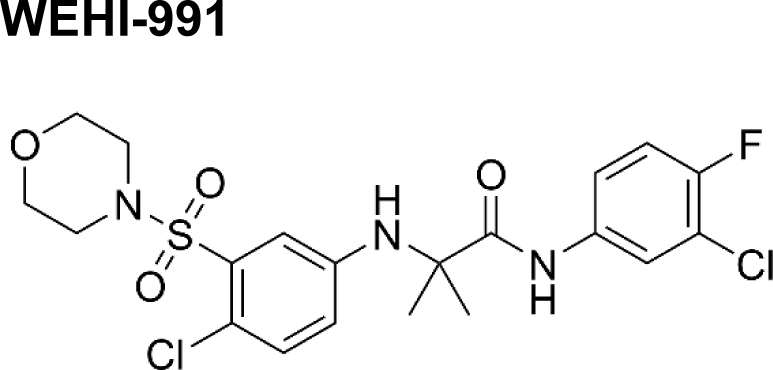

#### *N-(3-Chloro-4-fluoro-phenyl)-2-(4-chloro-3-morpholinosulfonyl-anilino)-2-methylpropanamide* (WEHI-991)

NaH (60% dispersed in mineral oil, 19.5 mg, 0.488 mmol) was added to a stirred solution of **Int-3** (90 mg, 0.33 mmol) in THF (2 mL). This was stirred for a further 10 min and then **Int-5** (150 mg, 0.52 mmol) was added, and the reaction stirred at 40°C for 16 h. The reaction was then quenched with 5% citric acid (1 mL) and concentrated. The crude was then suspended between EtOAc (10 mL) and washed with sat NaHCO3 (10 mL), and brine (10 mL). The organic layer was then dried with anhydrous Na2SO4, filtered and concentrated. The crude was then purified by column chromatography eluting with 100% DCM to 40% EtOAc/DCM to obtain **WEHI-991** as a solid (72 mg, 45%). ^1^H NMR (300 MHz, CDCl3): δ 8.66 (s, 1H), 7.71 (dd, *J* 6.5, 2.7 Hz, 1H), 7.29 - 7.38 (m, 3H), 7.08 (t, *J* 8.7 Hz, 1H), 6.64 (dd, *J* 8.7, 2.9 Hz, 1H), 4.35 (s, 1H) 3.66 - 3.74 (m, 4H), 3.21 - 3.29 (m, 4H), 1.60 (s, 6H). LCMS *m/z* 490.4 [M+1]. HRMS acquired: (M + H) 490.0769; C20H22Cl2FN3O4S requires (M + H), 490.0765.

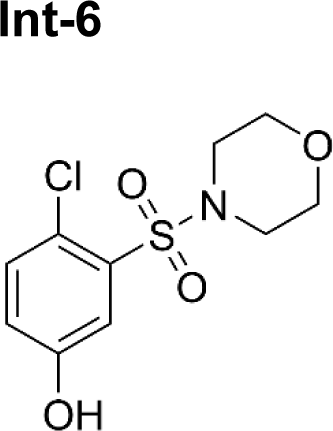

#### *4-Chloro-3-morpholinosulfonyl-phenol* (Int-6)

**Int-3** (310 mg, 1.1 mmol) was dissolved in H2O (2 mL) and the mixture stirred at 20°C. Concentrated sulfuric acid (0.30 mL, 5.7 mmol) was added drop wise through dropping funnel and the reaction mass was stirred for a period of 15 min. The reaction mixture was then cooled 0°C in an ice bath. Sodium nitrite (117 mg, 1.7 mmol) in H2O (2 mL) was then added dropwise to the reaction mixture. The resultant mixture was stirred at 0°C for 1 h. In another mL flask, ice-cold water (1 mL) was taken and added concentrated sulfuric acid (1 mL) to it slowly in 10 min. The mixture was warmed to 65°C and the above diazotised mixture was added dropwise. The resulting mixture was stirred at 100°C for 1 h. The reaction mixture was then poured onto ice water (20 mL) under stirring and the resulting mixture was extracted with EtOAc (3 × 15 mL). The combined organics were then washed with H2O (20 mL) and brine (20 mL), and then dried over anhydrous Na2SO4, filtered and concentrated. The crude was then purified by column chromatography eluting with 100% DCM to 50% EtOAc/DCM to obtain **Int-6** as an oil (125 mg, 40%). ^1^H NMR (300 MHz, CDCl3): δ 7.54 (d, *J* 3.1 Hz, 1 H), 7.40 (d, *J* 8.7 Hz, 1H), 7.00 (dd, *J* 8.7, 2.9 Hz, 1H), 3.71 - 3.78 (m, 4H), 3.28 - 3.33 (m, 4H). LCMS *m/z* 278.2 [M+1].

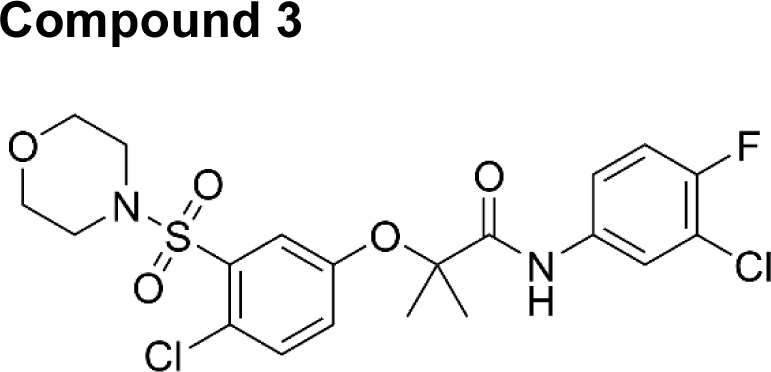

#### *N-(3-Chloro-4-fluoro-phenyl)-2-(4-chloro-3-morpholinosulfonyl-phenoxy)-2-methyl-propanamide* (3)

The procedure used for **WEHI-991** was followed using NaH (60% dispersed in mineral oil, 8.6 mg, 0.22 mmol), **Int-6** (30 mg, 0.11 mmol) and **Int-5** (64 mg, 0.22 mmol) to obtain crude product in 90% purity (39 mg, 66%). 20 mg of this crude was then purified by reverse phase preparatory HPLC using a gradient of 95% H2O/ACN to 100% ACN to obtain **3** as a solid (9.2 mg, 18%). ^1^H NMR (300 MHz, CDCl3): δ 8.37 (s, 1H), 7.78 (dd, *J* 6.5, 2.7 Hz, 1H), 7.69 (d, *J* 2.9 Hz, 1H), 7.48 (d, *J* 8.7 Hz, 1H), 7.36 - 7.43 (m, 1H), 7.07 - 7.17 (m, 2H), 3.66 - 3.79 (m, 4H), 3.23 - 3.36 (m, 4H), 1.61 (s, 6H). LCMS *m/z* 491.2 [M+1].

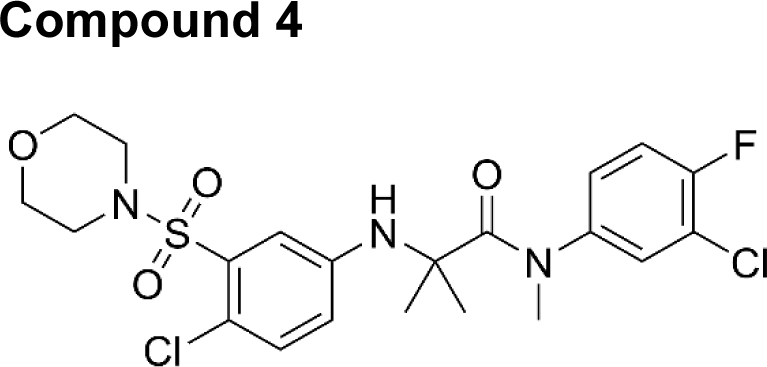

#### *N-(3-Chloro-4-fluoro-phenyl)-2-(4-chloro-3-morpholinosulfonyl-anilino)-N,2-dimethyl-propanamide* (4)

NaH (60% dispersed in mineral oil, 1.3 mg, 0.032 mmol) was added to a stirred solution of **WEHI-991** (7.8 mg, 0.016 mmol) in THF (1 mL) at 20°C and stirred for 10 min. Iodomethane (5 µL, 0.080 mmol) was then added and the reaction stirred for 16 h at this temperature. The reaction was then quenched with 5% citric acid (1 mL) and concentrated. The crude was then suspended between EtOAc (10 ml) and washed with saturated NaHCO3 (10 mL and brine (10 mL), and then dried with anhydrous Na2SO4, filtered and concentrated. The crude material was then purified by reverse phase preparatory HPLC using a gradient of 95% H2O/ACN to 100% ACN to obtain **4** as a solid (1.8 mg, 22%). ^1^H NMR (300 MHz, CDCl3): δ 7.31 - 7.36 (m, 1H), 6.94 - 7.09 (m, 3H), 6.80 - 6.90 (m, 1H), 6.52 - 6.62 (m, 1H), 3.73 - 3.78 (m, 4H), 3.21 - 3.32 (m, 7H), 1.55 (s, 6H). LCMS *m/z* 504.2 [M+1].

### Growth inhibition assays

Synchronised ring-stage parasites (0.3-0.6% parasitemia, 2% haematocrit) were grown for 72 h in a serial-dilution of the compound under investigation: MMV006833 (M-833), compound 2, WEHI-991 (W-991), compound 3 and compound 4. Note that M-833 was first purchased from MolPort (MolPort-002-267-916) then made in-house at WEHI: the differences observed in EC50s across experiments are attributed to the different batches of M-833. For some experiments, 0, 0.25 or 2.5 mM of glucosamine (GlcN) was also added. Experiments were set up in technical duplicates or triplicates. Following a freeze-thaw cycle, lactate dehydrogenase (LDH) activity was assessed as an indicator of parasite growth (Makler and Hinrichs, 1993). 30 μL of parasite culture was incubated for 30 min in the dark with 75 μL Malstat mixture (0.1 M Tris pH 9.0, 20 g/L lactic acid, pH 7.5, 0.2% Triton-X100, 0.5 g/L acetylpyridine adenine dinucleotide (APAD, Sigma), 200 μg/mL nitroblue tetrazolium (NBT; Sigma), 1 μg/mL phenozine ethosulfate (PES; Sigma)). Absorbance was then measured at 650 nm, transformed using the formula below, plotted against the logarithm of the concentrations, and a non-linear regression (log(inhibitor) vs. response, variable slope) was calculated using GraphPad Prism to assess the half-inhibitory concentration (EC50).

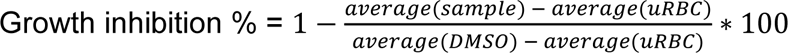

The DMSO samples were also used to assess growth of parasite in the presence of 0.25 and 2.5 mM GlcN (normalised by DMSO in 0 mM GlcN).

### Multiple cycle growth assays

To assess growth of parasites over several cycles, 3D7, PopD-D7, and PopE-F10 parasites were grown for 3 cycles, and growth was measured by LDH assay at the start and end of every 48 h cycle. Cultures at 0.3-1% trophozoite-stage, 2% haematocrit were set up in triplicates in a 96-well plate: duplicates were used for absorbance read-out, while the third replicate was used to make smears to look at the parasite morphology. Every 48 h, a subset of the culture was frozen (growth after 48 h), a subset was diluted and frozen straight away (baseline of the next cycle), and another subset was diluted and left to grow another 48 h. To analyse the result, the absorbance of each cycle at t = 48 h was divided by the absorbance at t = 0 h. The smears were used to check that there was no growth delay (*i.e.*, rings instead of trophozoites).

### Egress, invasion & recovery assay

To measure egress, invasion and follow up recovery in the presence of different compounds, we adapted the method developed in (Dans et al., 2020), using Hyp1-Nluc parasites (Azevedo et al., 2014). These parasites episomally express the exported protein Hyp1 tagged with a NanoLuc luciferase (NLuc) and were selected with 2.5 nM WR99210 (Jacobus). Highly synchronised schizonts were obtained using ML10 (LifeArc) and sorbitol synchronisation, and Percoll-purified (the culture was gently layered over 67% Percoll (Cytiva) in RPMI, span 15 min at 1500 g, the schizonts collected and washed). In a 96-well plate, cultures were set up in 4 replicates (3 replicates for the Nluc readout, and one replicate for smears) with 1-2% parasitemia, 2% haematocrit, with the following treatment: 2 μM M-883, 60 nM W-991, 0.02% DMSO (Sigma), 25 nM ML10 (LifeArc) 100 μg/ml heparin (Sigma) and 10 μM E64 (Sigma). The schizonts were allowed to egress and invade for 4 h at 37°C. A “background” plate without treatment was left in the fridge during this time. After 4 h, cultures were sorbitol-treated to eliminate schizonts that have not egressed. Smears were made every day for the following 3 days, and 5 μL of the resuspended cultures were taken to measure Nluc activity (to assess parasite’s presence). To measure Nluc activity, 5 μL culture were added to 45 μL NanoGlo reagent (1x Promega lysis buffer and NanoGlo reagent; Promega). The average luminescence of the background cultures was subtracted to the average luminescence of each sample.

### Stage arrest and recovery assay

Parasites were synchronised using a Percoll density gradient (Cytiva) combined with 5% sorbitol lysis. Ring-stage parasites at 0-4 hours post invasion at 2% parasitemia and haematocrit were then added to 2 µM M-833, 60 nM W- 991 or 0.02% DMSO and incubated at 37°C with compound replenished every 24 h. Every 12 h for the first lifecycle, Giemsa-stained thin blood smears were made and 3x 50 µL aliquots were taken and fixed with 0.25% gluteraldehyde (Electron Microscope Sciences) and stored at 4°C for remainder of experiment. After 48 h, samples of M-833 and W-991- treated parasites were washed x3 in complete RPMI to remove the compound and put back into culture for the remainder of the experiment. For the following two cycles of growth, samples were taken every 24 h to monitor recovery. Upon completion of time points, fixed cells were stained with 2.5x SYBR Green (Invitrogen) in PBS, washed once in PBS and analysed on the Attune Flow Cytometer (ThermoFisher Scientific). Giemsa-stained blood smears were visualised and imaged using a Nikon Eclipse E600 microscope.

### Parasite reduction ratio assay

This was performed as previously described (Sanz et al., 2012; Walz et al., 2023). Briefly, asynchronous ring-stage parasites at 0.5% parasitemia and 2% haematocrit were exposed to compounds at 10xEC50 of growth for five days with daily drug renewal. After each 24 h, a well from each treatment group was taken and washed three times in complete RPMI and final volume adjusted to 2% haematocrit. The washed culture was then aliquoted into a 96 well U-bottom plate and a 1:3 serial dilution into fresh uninfected RBCs was performed with four technical replicates for each time point. Plates were maintained in normal culturing conditions: media was replaced and fresh RBCs given 1-2 times a week for three weeks, after which parasite growth was measured via lactate dehydrogenase assays. The number of viable parasites was calculated using the formula x^n-1^ +1, where x is the dilution factor and n is the number of wells that contain parasite growth. The viable parasite value was then expressed in log units and parasite reduction ratio was calculated as a decrease in viable parasites over 48 h. The 99.9% parasite clearance rate was determined using a log-linear regression in GraphPad Prism.

### Lattice light sheet imaging

Parasites were sorbitol synchronised at day 5 and 2 prior to filming. To prepare the parasites for filming, culture was loaded on LS columns attached to MACS MultiStand (Miltenyi Biotec) to isolate late-stage parasites. Imaging medium was prepared by adding 10 μM Trolox (Santa Cruz 53188-07-1) to the culture medium. To compare effect of drug treatment on parasite invasion, either 60 nM of W-991 or equivalent DMSO concentration was added to the imaging medium. RBCs were resuspended at 0.5% hematocrit in RPMI-HEPES supplemented with 0.2% sodium bicarbonate and stained with 1.5 μM Di-4-ANEPPDHQ (Invitrogen D36802) membrane marker for 1 h at 37 °C. The stained RBCs were then washed three times and resuspended in imaging medium. Purified schizonts were resuspended in culture medium and incubated with 25 nM Mitotracker Deep Red FM (Invitrogen M22426) for 30 min at 37 °C, 5% CO2. The stained schizonts were then pelleted and the supernatant removed before resuspending the schizonts in imaging medium. Before imaging, imaging medium with drug was loaded to a well and imaging medium with DMSO control was loaded to another well on an 8-well glass bottom plate (Ibidi 80807). Stained RBCs and stained schizonts were then added to each well and let settle for 30 min. The imaging experiments were performed on Zeiss Lattice Lightsheet 7. 488 nm laser was used to excite Di-4- ANEPPDHQ and 640 nm laser was used to excite Mitotracker Deep Red FM. A quad notch filter was used to block 405/488/561/640 nm excitation lights. A 290 μm x 200 μm region was scanned with 0.3-0.4 μm interval for both drug-treated and control wells and a simultaneous timelapse was acquired at 2-3 ms exposure time with 15 s interval for 2 h. Acquired data were deskewed and deconvolved using Lattice Lightsheet processing on ZEISS ZEN Blue 3.4 software, then cropped into smaller regions for analyses.

#### Vacuole tracking

A data subset was used as training dataset for machine learning on Aivia 10.5.1 software. Single 3D frames were chosen and annotated as either ‘Background’ or ‘Want’ on Pixel Classifier analysis tool, where ‘Want’ is the vacuole area. The Pixel Classifier was applied on other data as feedback for training. Once the trained Pixel Classifier reached satisfactory accuracy, it was applied on the whole dataset in batch mode to obtain ‘Want’ channel, which is the vacuole confidence map, as additional channel to each file. 3D Object Tracking was then performed based on the ‘Want’ channel and the vacuoles of interest were isolated from the tracked objects. The sphericity of the vacuoles was extracted from the statistics of the isolated objects. The sphericity data were then plotted on GraphPad Prism 9.5.0 and two-tailed nested t-test was performed on the software to obtain p-value between the vacuole sphericity of drug-treated and DMSO control conditions at initial condition and 15 min after the vacuole is formed.

### Gametocyte culturing and standard membrane feeding assay

NF54 *P. falciparum* parasites (Walter Reed Army Institute of Research, USA), were synchronized and maintained as described above at 5-10% parasitemia at 4% hematocrit (WEHI ethics number HREC86/17). Once parasitemia reached 8-10% rings, parasite were diluted to 0.65% parasitemia and supplied with gametocyte media as outlined for asexual stage parasites and the substitution of Albumax for 10% Heat-inactivated human serum (Australian Red Cross Bloodbank) and no gentamycin. Mature sexual stage gametocytes were induced by allowing continuous growth cultures without the addition of fresh human RBCs to crash asexual stages and media was replaced daily (Vaughan et al., 2012). W- 991 (1 nM, 50 nM or 100 nM) or WM382 (50 nM) was added on day 13-17 with percentage of Stage V gametocytemia evaluated on day 17. Stage V gametocytes were then centrifuged at 13,000 rpm for 1 min to remove gametocyte media, 0.2 % of stage V gametocytes were resuspended with the mixture of fresh uninfected RBCs and heat-inactivated human serum at 2:3 ratio.

*Anopheles stephensi* mosquitoes were reared and maintained in the WEHI insectary. Mosquitoes were the bloodmeal as stated in (Favuzza et al., 2020). On day 7 after feeding, infected mosquitoes were aspirated into a cup and anesthetized by cooling to 4°C until immobilized and kept on ice for the dissection. The mosquitoes were then dissected as previously described (Favuzza et al., 2020) and collected midguts were maintained in 1X PBS (Gibco) until they were stained with mercurochrome for 20 mins. Oocysts were then quantified using light microscopy.

### Recombinant Expression and Purification

Codon optimised (*Spodoptera frugiperda*, Sf) START domain alone (I149-V394 from PF3D7_0104200) was synthesised by Integrated DNA Technologies and cloned into a modified baculovirus transfer vector (pAcGP67-A) with a N-terminal GP67-signal sequence, 8xHis tag, and TEV protease cleavage site. Recombinant baculovirus was generated via the flashBACTM Baculovirus Expression System using Sf21 cells (Life Technologies), amplified to reach a third passage viral stock, and 1.5% used to infect Sf21 for protein expression. After incubation for three days at 28°C, media containing secreted protein was harvested by centrifugation and supplemented with 50 mM Tris pH 7.5, 20 mM MgCl2, 100 mM NaCl and 10-20 mM imidazole. The supplemented protein sample was purified by nickel affinity chromatography (HisTrap Excel 5 mL, Cytiva), and eluted with 20 mM Tris pH 7.5, 500 mM NaCl, 300-500 mM imidazole. Eluted fractions were further purified using size-exclusion chromatography (Superdex S75 16/600 or Superdex 10/300, Cytiva) with the column pre-equilibrated with 20 mM HEPES pH 7.5, 150 mM NaCl, and peak containing fractions concentrated and stored at -80°C until required. If required, prior to ITC, a second size-exclusion chromatography step was performed to remove soluble aggregates.

*Spodoptera frugiperda* codon optimised full-length N309K and N330K PfSTART1 mutants were first synthesised by Integrated DNA Technologies, and then I149 forward and V394 reverse primers utilised to clone the same I149-V394 construct boundaries into the modified pAcGP67a expression vector with a N-terminal GP67-signal sequence, 8xHis tag, and TEV protease cleavage site. As a result of the subcloning, SSG was removed upstream of the TEV cleavage site. Construct boundaries and incorporation of the N309K and N330K mutations was verified by Sanger sequencing. The PfSTART1 mutants were expressed and purified as per wild-type PfSTART1.

### PfSTART1 polyclonal antibody generation and purification

Polyclonal rabbit anti-PfSTART1 antibodies were generated by the WEHI Antibody Facility using recombinant PfSTART1 (I149-V394). We further purified the polyclonal antibodies using a PfSTART1 affinity column that was generated using the AminoLink® Plus Immobilisation Kit (Thermo Scientific), according to the manufacturer’s instructions. For this, we immobilised deglycosylated (PNGaseF, NEB) and TEV cleaved PfSTART1 domain protein that had been further purified by nickel affinity chromatography to remove N-linked glycans and the 8xHis tag. The polyclonal antibody was purified on this matrix according to the manufacturer’s instructions.

### Isothermal titration calorimetry

ITC experiments were performed on a MicroCal PEAQ-ITC calorimeter at 25°C, with a stirring speed of 750 rpm and a reference power of 5 μcal/sec. 500 μM M-833 series inhibitor solutions were diluted to 10 μM in 50 mM NaPO4^-3^ (pH 7.4), 150 mM NaCl with a final DMSO concentration of 2% (v/v) (cell sample). PfSTART1 protein was dialysed extensively against 50 mM NaPO4-3 (pH 7.4), 150 mM NaCl, before diluting to 90 μM in the same buffer with 2% (v/v) DMSO (syringe sample). The first injection was 0.4 μL over a 0.8 s duration, and the remaining 19 injections were 2 μL of 4 s duration, with 150 s injection spacing. Data were collected and analysed using the PEAQ-ITC software (MicroCal) and fit by a single site binding model. A fitted offset (constant control heat) was also applied to the integrated heats.

### Solvent Profiling Western Blot

3D7 parasites at schizont stage were harvested with 10x pellet volume of 0.15% saponin in PBS. To extract soluble material, 10x pellet volume of 0.4% NP40 (IGEPAL CA-630, Sigma-Aldrich) in PBS with 1x Complete Protease cocktail tablet (Sigma-Aldrich) was added to the parasite pellets and 3 cycles of freeze/thawing was performed. Samples were then mechanically sheared by passage through 25G and 30G needles before lysate was cleared via centrifugation at 16,000 g x 30 min at 4°C. Soluble fractions were collected and stored at -80°C until use.

For solvent challenge, lysate was subjected to either W-991 (10 µM) or DMSO (0.1%) treatment for 3 min before aliquoted into final concentrations of 0-25% of Acetic acid/Ethanol/Formic acid (AEF) at a 50:50:0.1 (v/v/v) (Van Vranken et al., 2021; Yu et al., 2023). The treated lysate and AEF mixture was incubated at 37°C for 20 min at 800 rpm before aggregates were pelleted at 17,000 g x 20 min at 4°C. Soluble fractions were removed, added to final concentration of 1x NuPAGE LDS Sample Buffer (Invitrogen) with 1:100 2-mercaptoethanol (Sigma Aldrich) and boiled for 3 min. Proteins were separated on an 4-12% acrylamide gel (NuPAGE, Invitrogen) and subsequently transferred by electroblotting onto nitrocellulose membranes. Blots were probed with primary antibody anti-PfSTART1 (1:1000), followed by secondary antibody anti-rabbit-HRP (1:4000, Merck Millipore). ECL Plus Western blotting reagent (GE Healthcare) was used to visualise bands with the ChemiDoc Imaging System (Biorad).

### Solvent Proteome Profiling (MS)

Saponin-liberated mature parasite stages (30-42 hpi) were resuspended in PBS and lysed by 3x flash freeze/thawing using liquid N2, followed by 10x mechanical sheering with a 29-gauge needle-syringe, and soluble protein isolation through ultracentrifugation (100,000 g; 20 min, 4°C). Protein lysate was exposed to the drug (100 µM of W-991) or the vehicle control (DMSO) for 3 min and subsequently incubated with varying concentration of the solvent mixture ‘AEF’ (50% Acetone, 50% Ethanol, 0.1% Formate) to a final concentration of 7-25% (v/v) with 2% intervals, for 20 min at 37°C at 800 rpm on a Thermomixer (Eppendorf). The experiment was carried out in a biological triplicate. Denatured protein was pelleted through centrifugation (4°C, 18,000 g, 20 min) and the soluble phase was recovered and pulled together in equivolume ratios into two samples; Gradient 1 ‘G1’: 7-15% EAF and Gradient 2 ‘G2’: 17-25% EAF, respectively. Sample preparation for proteomic analysis was carried out using modified SP4 glass beads protocol (Johnston et al., 2022). In brief, protein was reduced (20 mM TCEP, 100 mM TEAB) for 20 min at 55°C and alkylated with 55 mM CAA for 30 min, followed by precipitation on beads in 80% ACN with a 6 min centrifugation at 21,000g and 3x wash with 80% Ethanol. Dried beads were subjected to sequential digestion with LysC (3 h) and trypsin (overnight) and the resulting digest was acidified with 1% TFA and desalted on T3 C18 stage tips (Affinisep).

#### MS data acquisition and data analysis

Peptide samples were analysed on an Orbitrap Eclipse Tribrid Mass Spectrometer (Thermo Scientific) in a DIA mode on a 45 min analytical gradient. Peptide identification was carried out in DIA-NN 1.8.1 using is silico spectral library generated from Uniprot *P. falciparum* (UP000001450) and human (UP000005640) reference proteomes. One missed cleavage and 1 variable modification was allowed (ox(M) and Ac(N-term)). Differential abundance data analysis (moderated t-test) of *P. falciparum* proteins was conducted in the R environment using precursor normalised MaxLFQ data for proteins detected with >³3 peptides. Hit selection criteria included p value <0.01, 0.73 log2 fold change (equivalent to 66% increase) in protein abundance and protein detection across all tested samples.

### Generation of *Plasmodium berghei* sporozoites

*P. berghei* ANKA constitutively expressing mCherry (Rathnapala et al., 2017) was used for the *in vitro* liver stage invasion assay. Animals used for the generation of the sporozoites were 4- to 5-week-old male Swiss Webster mice and were purchased from the Monash Animal Services (Melbourne, Victoria, Australia) and housed at 22 to 25°C on a 12 h light/dark cycle at the School of Biosciences, The University of Melbourne, Australia. All animal experiments were in accordance with the Prevention of Cruelty to Animals Act 1986, the Prevention of Cruelty to Animals Regulations 2008 and National Health and Medical Research Council (2013) Australian code for the care and use of animals for scientific purposes. These experiments were reviewed and permitted by the Melbourne University Animal Ethics Committee (2015123).

Infections of naïve Swiss mice were carried out by intraperitoneal (IP) inoculation obtained from a donor mouse between first and fourth passages from cryopreserved stock. Parasitemia was monitored by Giemsa smear and exflagellation quantified 3 days post-infection. *A. stephensi* mosquitoes were allowed to feed on anaesthetised mice once the exflagellation rate was assessed between about 12 to 15 exflagellation events per 1 × 10^4^ RBCs. Salivary glands of infected mosquitoes (days 17 to 24 post infection) were isolated by dissection and parasites placed into RPMI-1640 media.

### *In vitro* liver invasion assays

This was performed essentially as described (Wilson et al., 2015) with the following variations. *In vitro* human liver HCO4 cells (ATCC) were seeded at 1 × 10^5^ cells/mL, grown for 24 h and then infected for 2 h with 30,000 *Pb* ANKA mCherry expressing sporozoites. Following fixation, an IFA with anti-CSP antibody (1:500) followed by anti-mouse Alexafluor 488 (1:1,000; Invitrogen, Carlsbad, CA, USA) labels uninvaded parasites. Fluorescence signal was assessed using an Olympus CKX41 epifluorescence microscope.

### Dual gamete formation assays

The compounds were tested in the *P. falciparum* Dual Gamete Formation Assay (PfDGFA) (Delves et al., 2016). Briefly, mature stage V gametocytes were exposed to compounds for 48 h at 37°C in 384 well plates in gametocyte culture medium (RPMI 1640 supplemented with 25 mM HEPES, 50 μg/mL hypoxanthine, 4.8 g/L NaHCO3, 2 mM L-glutamine, 5% pooled type AB serum, 0.5% Albumax II (Gibco)) under a 1% O2, 3% CO2, 96% N2 environment. Gametogenesis was then triggered by the addition of 10 μL ookinete medium (gametocyte culture medium supplemented with 100 μM xanthurenic acid and 0.27 µg/mL Cy3-labelled anti-Pfs25 antibody) to each well at room temperature. Plates were then cooled on a metal block at 4°C for four min to ensure even cooling and then stabilised for a further 4 min at 28°C. At 20 min post-induction, male gametogenesis was recorded in each well by automated brightfield microscopy using a x4 objective lens and 1.5x magnifier (x6 effective magnification). Afterward, plates were incubated in the dark at room temperature for 24 h and then female gametogenesis recorded in each well by automated fluorescence microscopy (anti-Pfs25-positive cells). All experiments were performed in quadruplicate with DMSO and Cabamiquine as negative and positive controls respectively. All data was evaluated in comparison to the positive and negative controls to calculate percentage inhibition of male and female gametocytes, and dose response analysis and IC50 calculation performed using GraphPad Prism.

### Protein extraction & Western Blot

Parasites were prepared for the stage of interest and proteins extracted using a saponin-lysis (unless indicated otherwise). For knock-down assays, parasites were exposed to 0 or 2.5 mM GlcN for 48 h (starting from late schizont / early rings): the day before harvesting schizonts, 30 nM ML10 (LifeArc) was also added to the cultures to block egress. Infected red blood cells were lysed 10 min on ice with 0.1% saponin in PBS complemented with 1x Protease Inhibitors Cocktail (Roche; PBS+PI). Parasites were pelleted and washed with PBS+PI to remove haemoglobin. The pellets were resuspended in 10 to 20 x volume of non-reducing sample buffer (NRSB; 50 mM Tris-HCl pH 6.8, 2 mM EDTA, 2% SDS, 10% glycerol, 0.005% phenol blue), sonicated a minimum of 3 x 30 seconds (Diagenode sonicator), were optionally reduced with 100 mM dithiothreitol, and boiled 10 min at 80°C. Protein samples were centrifuged at 15 000 g, then run on a pre-cast 4-12% NuPAGE Bis-Tris gel (Invitrogen) and proteins were transferred onto a nitrocellulose membrane using iBlot (Invitrogen). Membranes were exposed to the primary antibody overnight at 4°C, to the secondary antibody 1 h at room temperature, and fluorescence measured using an Odyssey imaging system, which was also used to measure densitometry. The list of antibodies used, their dilution and origins are described in Table S4.

### Processing of PfSTART1 along schizogony

To pinpoint when PfSTART1 was processed during schizogony, magnet-purified schizonts were harvested after a 4 h treatment with the following: DMSO (0.1%), 49c (10 nM), C1 (1.5 μM) or E64 (10 μM). Parasites were washed in ice-cold PBS+PI (centrifugation steps carried out at 4°C at 3,000 g). Samples were resuspended in NRSB, sonicated, boiled 10 min at 80°C and separated by electrophoresis as explained previously. The parasites used in this experiment were 3D7 transfected with p1.2-ABH(WT)-83-HA-*glm*S, *i.e*. parasites containing a recodonised version of PF3D7_0403800 within its genome.

### Differential lysis assay

To localise PfSTART1, schizont-infected RBCs were sequentially lysed with (1) equinatoxin II (EqtII), which lyses the RBC membrane, (2) saponin, which lyses the PVM, (3) TX-100, which lyses all membranes. All the lysis buffers were made in PBS+PI, supernatants were collected in fresh tubes, and pellets were wash twice in PBS+PI. ML10-treated schizonts were enriched (with Percoll) and lysed in 10x pellet volume of EqtII (at a concentration empirically determined to obtain 100% hemolysis) for 10 min at 37°C. Lysate was centrifuged at 1,000 g for 5 min to collect the supernatant. The pellet was washed then lysed in 10x volume of 0.03% saponin (10 min on ice, centrifuged at 16,000 g for 1 min at 4°C). The supernatant was collected, the pellet was washed, and lysed in 10x volume of 0.25% TX-100 (10 min on ice, centrifuged at 16,000 g for 1 min at 4°C, and the supernatant collected). All supernatants were mixed with NRSB, boiled 10 min at 80°C and separated by electrophoresis as explained previously.

### Carbonate extraction

To determine whether PfSTART1 is soluble, membrane bound or membrane associated, a carbonate extraction was conducted as previously described (Bullen et al., 2022). 3D7 schizonts were lysed in 0.1% saponin as described above. The pellet was subjected to five freeze-thaw cycles in 20x pellet volume of PBS+PI (freeze 1 min in a dry ice-ethanol bath; thaw 2 min at 37°C). The ‘PBS soluble fraction’ was recovered after a 30 min centrifugation step (17,000 g at 4°C). After a wash in PBS+PI, the pellet was incubated 30 min in 0.1 M Na2CO3 (pH 11) at room temperature. The ‘carbonate soluble fraction’ was recovered after another 30 min centrifugation step, the ‘insoluble fraction’ pellet was washed and resuspended in NRSB to a final volume equivalent to the ‘PBS soluble’ and ‘carbonate soluble’ fractions (also complemented with NRSB). Samples were boiled 10 min at 80°C and run on a Western Blot as described previously.

### Proteinase K protection assay

To further determine the localisation of PfSTART1 in schizonts, we conducted a proteinase K protection assay (Jonsdottir et al., 2023). Percoll-purified schizonts were washed in PBS+PI, and lysed in 10x the pellet volume of EqtII (to achieve 100% hemolysis) for 10 min at 37°C. The EqtII supernatant was collected, the pellet was washed 3 times in PBS without protease inhibitors and divided equally between six tubes. The pellets were incubated 20 min on ice with 100 μL: (1-2) PBS -/+ 20 μg/mL proteinase K, (3-4) 0.005% saponin in PBS -/+ 20 μg/mL proteinase K, (5-6) 0.005% saponin + 1% TX-100 in PBS -/+ 20 μg/mL proteinase K. Proteinase K activity was stopped with a final concentration of 10% of trichloroacetic acid (TCA, Sigma). Proteins were pelleted 20 min at 17,000 g at 4°C, the supernatant was discarded, and 500 μL cold acetone (Merck) was added and left overnight at -20°C. Samples were centrifuged another 10 min, the supernatant was discarded, the pellets were left to air-dry and were then resuspended in NRSB.

### Immunofluorescence assay

SLI-WT parasites were allowed to settle onto 12 mm coverslips pre-treated with 10% Poly-L-Lysine (Sigma) in Milli-Q water. Cells were fixed 20 min with 4% paraformaldehyde (Emgrid) and 0.0075% glutaraldehyde (ProSciTech) in PBS at room temperature. Cells were then washed in PBS, and lysed for 10 min with 1 M Glycine, 0.1% TX100 in PBS. Cells were blocked 1 h at room temperature in 3% BSA, 0.01% TX100 in PBS, primary antibody was probed overnight at 4°C, secondary antibody was probed for 1 h at room temperature (with 5% goat serum). Antibodies used, their provider and their concentrations are described in Table S4. VectaShield Antifade Mounting Medium (with DAPI, H-1200) was added before sealing the coverslips onto a slide. Parasites were imaged using a Zeiss Cell Axio Observer (Carl Zeiss), and images were processed with ImageJ. Merozoites from a single experiment (with imaging settings maintained throughout image acquisition) were selected, and co-localisation between PfSTART1-HA and markers PfMSP1-19, PfHSP70-1, PfEXP2 and PfGAPm3 was assessed. To determine the colocalisation assessment of this data the approach outlined by BIOP’s (Bioimaging and Optics Platform) version of JACoP (Just Another Colocalization Plugin) https://c4science.ch/w/bioimaging_and_optics_platform_biop/image-processing/imagej_tools/jacop_b/ (Bolte and Cordelieres, 2006; Cardona et al., 2012) was applied in conjunction with a FIJI script developed to batch process and reduced noisy artefacts in the acquired microscopy images. The script coverts a set of images into 8-bit format, chosen to reduce noise and balance the intensity precision. The conversion reduces inherent noise, simplifies thresholding operations and enhances feature visibility during the image processing process. Images are split into their channels and a background subtraction is conducted to emphasise distinctions between object features and their surrounding background. Once this is complete the BIOP JACoP plugin is called and the Persons correlation coefficient for colocalisation analysis is computed and the resultant data is saved in an output directory

## Acknowledgements

We acknowledge the traditional custodians of the lands on which this project was conducted: the Wurundjeri and the Boon Wurrung people of the Kulin nation. We thank the Lifeblood Biological Resources Australia for providing the human red blood cells and LifeArc for supplying ML10. We thank Danu Marapana for kindly sharing p1.2 CRISPR plasmid. We acknowledge Hayley Bullen for the PfGAPm3 antibody and Leann Tilley for the ERC antibody. We thank the WEHI screening facility for conducting parasite growth assays with newly synthesised compounds. This work was supported by the Victorian Operational Infrastructure Support Program received by the Walter and Eliza Hall and Burnet Institutes. This work was funded by the National Health and Medical Research Council of Australia (Ideas Grant to W.N. and P.G. 2001073; Development Grant 1135421 to B.E.S. and A.F.C.). A.F.C. is a Howard Hughes International Scholar and an Australia Fellow of the NHMRC. B.E.S. is a Corin Centenary Fellow. MTF is supported by a grant from the Medicines for Malaria Venture (RD-21-2003) awarded to MJD. MJD is supported by a UKRI Medical Research Council Career Development Award (MR/V010034/1).

## Supplementary Figure legends

**Supplementary Figure 1. Growth inhibition of M-833 resistant populations, PfSTART1 mutations in PopD-D7, PopA and PopC. (A)** Resistance against M-833 could be generated against three out of five populations of parasites eliciting an 8 to 37 fold increase in EC50. **(B)** *Pfstart1* was PCR amplified from gDNA of the M-833 resistant clone D7 from population D (PopD-D7), and its sequencing indicates that PopD-D7 also contains mutation N309K, similar to other clones from this resistant population. **(C)** Similarly, *pfstart1* from PopC was PCR amplified and sequenced. PopC displayed a I224F mutation.

**Supplementary Figure 2. Engineering PfSTART1 WT, N309K or N330K using Selection Linked Integration (SLI). (A)** Schematic of the cloning strategy. The 5’ arm of native *pfstart1* was amplified, PCR sewn to a recodonised version of the 3’ arm (either WT, N309K or N330K), inserted into a p-HA-2A-Neo-glmS plasmid and transfected into 3D7 parasites, to be integrated into the native gene. **(B)** Transfectant parasites were cloned: SLI-WT clone B4, SLI-N309K clone D6 and SLI-N330K clone G3, thereafter called SLI-WT, SLI-N309K and SLI-N330K. gDNA was extracted and a PCR was conducted to check for correct integration using primers START_5’UTR.F and START_Seq.R (Table S3); the same PCR on 3D7 gDNA was used as a control, as no bands should be observed in the 3D7 samples. **(C)** The sequencing of the PCR amplified *pfstart1* genes shows that SLI-WT has the wild-type version of START, and that SLI- N309K and SLI-N330K contain the correct mutations. **(D)** 3D7, SLI-WT, -N309K or - N330K were exposed for 48 h to 0 or 2.5 mM GlcN, and proteins extracted from schizonts. Western blots were probed with anti-PfSTART1, anti-HA, anti-PfHSP70-1 and anti-PfEXP2 antibodies. The densitometry of 3 biological replicates was measured (the PfSTART1 signal was normalised by the corresponding PfEXP2 signal, and then normalised by the 0 mM GlcN condition within each parasite line). **(E)** Growth of 3D7, SLI-WT, -N309K or -N330K over 72 h in the presence of 0, 0.25 or 2.5 mM GlcN was measured with an LDH assay. Growth was normalised to the 0 mM GlcN condition. n=4 biological replicates. Ordinary one-way ANOVA with Dunnett’s multiple comparison test (comparison to the 0 mM condition). **: p<0.005. ***: p<0.0005. ****: p<0.0001.

**Supplementary Figure 3. Engineering PfSTART1 WT, N309K or N330K using CRISPR-Cas9 (CR). (A)** Schematic of the cloning strategy. The 5’ arm of native *pfstart1* was amplified, sewn to a recodonised version of the 3’ arm (either WT, N309K or N330K), inserted into a p1.2-glmS (Marapana et al., 2018) plasmid, along with the native 3’ arm of *pfstart1*. Note that this construct does not include a HA tag. The plasmid was transfected into 3D7 parasites with Cas9 enzyme complexed with a guide RNA, in order to be integrated into the native gene. **(B)** Transfectant parasites were cloned: CR-WT clone D2, CR-N309K clone C10 and CR-N330K clone B2, thereafter called CR-WT, CR-N309K and CR-N330K. gDNA was extracted and a PCR was conducted to check for correct integration using primers START_5’UTR.F and START_IntR1; the same PCR on 3D7 gDNA was used as a control, as no bands should be observed in the 3D7 samples. **(C)** The PCR product was sequenced and indicated that CR-WT has the wild-type version of *pfstart1*, and that CR-N309K and CR-N330K contain the correct mutations. **(D)** 3D7, CR- WT, -N309K or -N330K were exposed for 48 h to 0 or 2.5 mM GlcN, and proteins extracted from schizonts. Western blots were probed with anti-PfSTART1, anti-PfHSP70- 1 and anti-PfEXP2 antibodies. The densitometry of 3 biological replicates was measured (the PfSTART1 signal was normalised to the corresponding EXP2 signal, and then normalised by the 0 mM GlcN condition within each parasite line). **(E)** Growth of 3D7, CR- WT, -N309K or -N330K over 72 h in the presence of 0, 0.25 or 2.5 mM GlcN was measured by LDH assay. Growth was normalised to the 0 mM GlcN condition. n=6 biological replicates. Ordinary one-way ANOVA with Dunnett’s multiple comparison test (comparison to the 0 mM condition). ***: p<0.0005. ****: p<0.0001.

**Supplementary Figure 4. Growth inhibition assays of M-833 and analogues on M- 833 resistant parasites.** Ring stage drug sensitive 3D7 parasites and M-833-resistant PopD-D7 (N309K mutant), and PopE-F10 (N330K mutant) parasites were exposed 72 h to a dilution series of M-833 **(A)**, compound 2 **(B)**, W-991 **(C)**, compound 3 **(D)**, and compound 4 **(E)**. The structure of the compounds and the EC50 values can be found in Fig 3 and Table S2.

**Supplementary Figure 5. Isothermal titration calorimetry analysis of M-833 series and PfSTART1 and other START-domain containing proteins in solvent proteome profiling assays. (A)** Size-exclusion chromatogram of recombinant WT PfSTART1 (black), N330K (pink), and N309K (teal). Grey box indicates fractions used for ITC experiments. **(B)** SDS-PAGE analysis of recombinant PfSTART1 proteins used for ITC experiments. **(C.i)** Second replicate thermograms of the M-833 series binding to recombinant PfSTART1(WT) or PfSTART1(N330K). The bottom panel comprises the data after integration of the peaks and a fitted offset applied. The binding curve shows the fit to a single-site binding model. DP = differential power. **(ii)** Summary of the individual thermodynamic parameters obtained for n=2. Error represents the error of the fit. **(D)** Replicate western blots of solvent proteome assays (Fig 4B.i). **(E)** No significant difference was found between DMSO or W-991 treated parasite lysate in relative soluble protein abundance of Pf3D7_1351000 or Pf3D7_1463500 after solvent-induced protein precipitation, plotted for Gradient 1 (7-15% AEF) and Gradient 2 (17-25% AEF). Error bars represent the standard deviation of three biological replicates.

**Supplementary Figure 6. PfSTART1 inhibitors have no effect on intraerythrocytic development and exhibit a slow killing profile in a parasite reduction ratio assay. (A)** Highly synchronous ring-stage 3D7 parasites were exposed to M-833 (2 µM), W-991 (60 nM) or DMSO (0.02%) and every 12 h for one growth cycle, Giemsa-stained thin blood smears were taken to visualise parasite morphology. **(B.i)** Ring-stage parasites were exposed to 10xEC50 of chloroquine (CQ) or W-991 for up to five days. An aliquot of the treated culture was removed daily, compounds were washed out and the parasites serially diluted. After 21 days, parasite growth was quantified via LDH assay and log(viable parasites+1) was calculated. W-991 demonstrated a ‘slow’ killing profile when compared to the fast-acting CQ. **(B.ii)** W-991 showed more than 24 h of lag phases (0- 24h, 48-96h) whereby it did not reach its maximum rate of killing. Log parasite reduction ratio (log(PRR)) was calculated by the reduction of three log units between one cell cycle (0-48 h). Parasite clearance time (PCT) was calculated using a linear regression not including lag phases of the compounds. Error bars represent the standard deviation of three biological replicates (values in brackets in table) and linear regressions were calculated using GraphPad Prism. Concentrations used equated to 75 nM and 60 nM for CQ and W-991, respectively.

**Supplementary Figure 7. Measurements of parasite vacuoles after merozoite invasion using lattice light sheet microscopy. (A)** Individual vacuole sphericity quantification directly after merozoite invasion for both DMSO and W-991 treatments. Averages can be seen in Fig 6A.ii. A reduction in both the surface area **(B)** and volume **(C)** of the vacuoles was also observed upon W-991 treatment which was found to not be significant via nested t-tests at 15 mins between DMSO and W-991-treated parasites (p=0.109 and p=0.129, respectively).

**Supplementary Figure 8. W-991 shows no activity against the liver stages or gamete formation. (A)** *In vitro* HC-04 human liver cells were incubated with DMSO, W-991 or cytochalasin D (cytD) in the presence of 30,000 sporozoites expressing mCherry for 2 h, followed by fixation with 4% paraformaldehyde. W-991 treatment showed no defect in invasion as measured by comparing mCherry (all sporozoites) and anti-CSP antibody fluorescence (uninvaded sporozoites) when compared with vehicle control. Statistical analysis performed via an unpaired t-test between DMSO and drug treatments. Error bars represent the standard deviation of two replicates. *** *P* < 0.001. **(B)** Values indicate percent inhibition of 1 µM M-833 and W-991 against female and male gamete formation in *P. falciparum*.

**Supplementary Figure 9. PfSTART1 expression, processing and localisation. (A.i- ii)** Replicates 2 and 3 of the time course of SLI-WT parasites along their 48 h erythrocytic cycle (replicate 1 and densitometry analysis are shown in Fig 7). A saponin-lysis was performed on synchronised SLI-WT young rings, old rings, young trophozoites, and schizonts. **(A.iii)** representative smears of corresponding parasites. **(B.i)** Synchronised 3D7 and SLI-WT schizonts were stalled at different stages, following a 4 h treatment with 25 nM ML10, 10 µM E64, or were allowed to progress (DMSO). Proteins were harvested with a saponin-lysis, and western blot probed with anti-PfSTART1 and anti-HA antibodies. **(B.ii)** Tightly synchronised 3D7 schizonts were magnet-purified, stalled at different stages of egress, following a 4 h treatment with 10 nM 49c (plasmepsin X inhibitor), 1.5 µM compound 1 (C1; PKG inhibitor), 10 µM E64 (cysteine protease inhibitor), or allowed to progress (0.015% DMSO). Whole cells were harvested, loaded onto a Western blot, probed with anti-PfSTART1 antibody. PEXEL-cleaved, and the two further processed forms of PfSTART1 are indicated by solid and empty arrows respectively. **(B.iii)** Densitometry of processed PfSTART1 (processed 1 and processed 2) was normalised to the corresponding full length PfSTART1. n=10 biological replicates, in technical duplicates. Two-way ANOVA with Šídák’s multiple comparisons test (compare each treatment to one another, within each processed PfSTART). *: p<0.05. **: p<0.005. ***: p<0.0005. ****: p<0.0001. **(C)** To localise PfSTART1, Percoll-purified 3D7 schizonts were sequentially lysed with equinatoxin II (Eqt II), saponin (Sap) and Triton-X100 (TX100), and the supernatants (SN) were collected to harvest proteins localising in the red blood cell (RBC) cytosol, the parasitophorous vacuole (PV) and the parasite, respectively. PfGBP130 is exported into the RBC cytosol; PfSERA5 localises in the PV in schizonts; PfHSP70-1 is a parasite cytosolic protein. Another replicate is shown in Fig 7. **(D)** To determine whether the solubility of PfSTART1, saponin-lysed 3D7 schizonts were sequentially lysed in PBS (with five freeze-thaw cycles) and sodium carbonate. The supernatant resulting from the PBS lysis was collected (“PBS soluble”), as well as the supernatant and the pellet of the carbonate lysis (“Carbonate soluble” and “Carbonate insoluble” respectively). Controls are the soluble protein PfHSP70-1, the membrane-associated PfHSP101, and the integral transmembrane protein PfEXP2. Another replicate is shown in Fig 7. **(E)** Proteinase K protection assay was conducted on Percoll-purified 3D7 schizonts. Schizonts were first lysed in EqtII, the supernatant (SN) of which was collected. The remaining parasite and PV were either incubated in PBS (no lysis), saponin (PVM-lysis) or TX-100 (lysis of all membranes) with or without proteinase K. PfGBP130 is RBC cytosolic protein; PfPTEX150 is a PV protein; PfActin-1 is a parasite cytosolic protein. Another replicate is shown in Fig 7.

**Supplementary Figure 10. Immunofluorescence assay of SLI-WT parasites.** Tightly synchronised SLI-WT parasites were probed with different markers of parasite organelles, and parasites of the stage of interest were imaged with a Zeiss Cell Axio Observer (Carl Zeiss). Representative images are shown for schizonts **(A)**, merozoites **(B)** and rings **(C)**. PfSTART1 was detected with an HA antibody (shown in red). The following antibodies were used to mark the following localisations (in green): PfAMA1 – micronemes (Mn); PfEXP2 – dense granules (DG); PfRhopH3 – rhoptries (Rhop); PfERC – endoplasmic reticulum (ER); PfHSP70-1 – cytosol; PfGAPm3 – inner membrane complex (IMC); PfMSP1-19 – merozoite plasma membrane (PM); human Band3 – RBC membrane. Nuclei were stained with DAPI (in blue). Scale bar = 2 µm (schizonts and rings) or 1 µm (merozoites).

## Supplementary Scheme legends

**Scheme S1.** Synthetic route to generate **2**. *Reagents and conditions:* (a) chlorosulfonic acid, 120°C; (b) morpholine, DCM, 0°C; (c) H2O: Zn, saturated NH4Cl (1:1), 40°C; (d) 2- chloroacetyl chloride, TEA, DCM, 20°C; (e) **Int-3**, KI, DMF, 45°C.

**Scheme S2.** Synthetic route to generate **WEHI-991**. *Reagents and conditions:* (a) 2- bromoisobutyryl bromide, TEA, DCM, 20°C; (b) **Int-3**, NaH, THF, 40°C.

**Scheme S3.** Synthetic route to generate **3**. *Reagents and conditions:* (a) sodium nitrite, H2SO4, H2O, 0°C →65°C →100°C; (b) **Int-3**, NaH, THF, 40°C.

**Scheme S4.** Synthetic route to generate **4**. *Reagents and conditions:* (a) MeI, NaH, THF, 20°C.

## Supplementary Table legends

**Supplementary Table 1. M-833-resistant genomes pooled variant summary.** Table depicts genes in which non-synonymous SNPs were identified that passed quality filtration and were detected in at least one resistant clone. Highlighted rows indicate putative targets.

**Supplementary Table 2. EC50 values of M-833 analogues on M-833-resistant clones.** PopD-D7 contains the PfSTART1 mutation N309K; PopE-F10 contains the PfSTART1 mutation N330K. CI: 95% confidence interval. Fold-change compared to the EC50 on 3D7 parasites is also indicated.

**Supplementary Table 3. List of primers used in this project.**

**Supplementary Table 4. List of antibodies used in this project.**

## Supplementary Video legends

**Supplementary movie 1. RBC invasion with DMSO control.** Merozoites are stained in cyan (with Mitotracker Deep Red CMXRos) and the RBC membrane is stained in magenta (with Di-4-ANEPPDHQ)

**Supplementary movie 2. RBC invasion with 60 nM W-991 control.** Merozoites are stained in cyan (with Mitotracker Deep Red CMXRos) and the RBC membrane is stained in magenta (with Di-4-ANEPPDHQ)

## References

Akula, N., Midzak, A., Lecanu, L., and Papadopoulos, V. (2012). Identification of small-molecule inhibitors of the steroidogenic acute regulatory protein (STARD1) by structure-based design. Bioorg Med Chem Lett 22, 4139–4143.

Asif, K., Memeo, L., Palazzolo, S., Frión-Herrera, Y., Parisi, S., Caligiuri, I., Canzonieri, V., Granchi, C., Tuccinardi, T., and Rizzolio, F. (2021). STARD3: A Prospective Target for Cancer Therapy. In Cancers.

Azevedo, M.F., Nie, C.Q., Elsworth, B., Charnaud, S.C., Sanders, P.R., Crabb, B.S., and Gilson, P.R. (2014). *Plasmodium falciparum* Transfected with Ultra Bright NanoLuc Luciferase Offers High Sensitivity Detection for the Screening of Growth and Cellular Trafficking Inhibitors. PLoS ONE 9, e112571.

Bailly, E., Jambou, R., Savel, J., and Jaureguiberry, G. (1992). *Plasmodium falciparum*: Differential Sensitivity In Vitro to E-64 (Cysteine Protease Inhibitor) and Pepstatin A (Aspartyl Protease Inhibitor). The Journal of Protozoology 39, 593–599.

Barnes, C.B.G., Dans, M.G., Jonsdottir, T.K., Crabb, B.S., and Gilson, P.R. (2022). PfATP4 inhibitors in the Medicines for Malaria Venture Malaria Box and Pathogen Box block the schizont-to-ring transition by inhibiting egress rather than invasion. Frontiers in Cellular and Infection Microbiology 12.

Bergmann, C., van Loon, W., Habarugira, F., Tacoli, C., Jäger, J.C., Savelsberg, D., Nshimiyimana, F., Rwamugema, E., Mbarushimana, D., Ndoli, J., et al. (2021). Increase in Kelch 13 Polymorphisms in *Plasmodium falciparum*, Southern Rwanda. Emerg Infect Dis 27, 294–296.

Birnbaum, J., Flemming, S., Reichard, N., Soares, A.B., Mesén-Ramírez, P., Jonscher, E., Bergmann, B., and Spielmann, T. (2017). A genetic system to study *Plasmodium falciparum* protein function. Nature methods 14, 450–456.

Blackman, M.J. (2008). Malarial proteases and host cell egress: an ‘emerging’ cascade. Cell Microbiol 10, 1925–1934.

Bolte, S., and Cordelieres, F.P. (2006). A guided tour into subcellular colocalization analysis in light microscopy. Journal of Microscopy 224, 213–232.

Bopp, S., Pasaje, C.F.A., Summers, R.L., Magistrado-Coxen, P., Schindler, K.A., Corpas-Lopez, V., Yeo, T., Mok, S., Dey, S., Smick, S., et al. (2023). Potent acyl-CoA synthetase 10 inhibitors kill *Plasmodium falciparum* by disrupting triglyceride formation. Nature Communications 14, 1455.

Burns, A.L., Dans, M.G., Balbin, J.M., deKoning-Ward, T.F., Gilson, P.R., Beeson, J.G., Boyle, M.J., and Wilson, D.W. (2019). Targeting malaria parasite invasion of red blood cells as an antimalarial strategy. FEMS Microbiol Rev 43, 223.

Bozdech, Z., Llinás, M., Pulliam, B.L., Wong, E.D., Zhu, J., and DeRisi, J.L. (2003). The Transcriptome of the Intraerythrocytic Developmental Cycle of *Plasmodium falciparum*. PLoS Biol 1, e5.

Bullen, H.E., Sanders, P.R., Dans, M.G., Jonsdottir, T.K., Riglar, D.T., Looker, O., Palmer, C.S., Kouskousis, B., Charnaud, S.C., Triglia, T., et al. (2022). The *Plasmodium falciparum* parasitophorous vacuole protein P113 interacts with the parasite protein export machinery and maintains normal vacuole architecture. Molecular Microbiology 117, 1245–1262.

Cardona, A., Saalfeld, S., Schindelin, J., Arganda-Carreras, I., Preibisch, S., Longair, M., Tomancak, P., Hartenstein, V., and Douglas, R.J. (2012). TrakEM2 software for neural circuit reconstruction. PLoS One 7, e38011.

Chitrala, K.N., and Yeguvapalli, S. (2014). Computational screening and molecular dynamic simulation of breast cancer associated deleterious non-synonymous single nucleotide polymorphisms in TP53 gene. PLoS One 9, e104242.

Clark, B.J. (2020). The START-domain proteins in intracellular lipid transport and beyond. Molecular and Cellular Endocrinology 504, 110704.

Dans, M.G., Weiss, G.E., Wilson, D.W., Sleebs, B.E., Crabb, B.S., de Koning-Ward, T.F., and Gilson, P.R. (2020). Screening the Medicines for Malaria Venture Pathogen Box for invasion and egress inhibitors of the blood stage of *Plasmodium falciparum* reveals several inhibitory compounds. International Journal for Parasitology 50, 235–252.

Dans, M.G., Piirainen, H., Nguyen, W., Khurana, S., Mehra, S., Razook, Z., Geoghegan, N.D., Dawson, A.T., Das, S., Parkyn Schneider, M., et al. (2023). Sulfonylpiperazine compounds prevent *Plasmodium falciparum* invasion of red blood cells through interference with actin-1/profilin dynamics. PLOS Biology 21, e3002066.

Delves, M.J., Miguel-Blanco, C., Matthews, H., Molina, I., Ruecker, A., Yahiya, S., Straschil, U., Abraham, M., León, M.L., Fischer, O.J., et al. (2018). A high throughput screen for next-generation leads targeting malaria parasite transmission. Nat Commun 9, 3805.

Delves, M.J., Straschil, U., Ruecker, A., Miguel-Blanco, C., Marques, S., Dufour, A.C., Baum, J., and Sinden, R.E. (2016). Routine in vitro culture of *P. falciparum* gametocytes to evaluate novel transmission-blocking interventions. Nature Protocols 11, 1668–1680.

Derré, I., Swiss, R., and Agaisse, H. (2011). The Lipid Transfer Protein CERT Interacts with the Chlamydia Inclusion Protein IncD and Participates to ER-Chlamydia Inclusion Membrane Contact Sites. PLOS Pathogens 7, e1002092.

Dietrich, M.H., Chan, L.J., Adair, A., Boulet, C., O’Neill, M.T., Tan, L.L., Keremane, S., Mok, Y.F., Lo, A.W., Gilson, P., et al. (2022). Structure of the Pf12 and Pf41 heterodimeric complex of *Plasmodium falciparum* 6-cysteine proteins. FEMS Microbes 3, xtac005.

Doerr, A. (2015). DIA mass spectrometry. Nature methods 12, 35–35.

Dondorp, A.M., Nosten, F., Yi, P., Das, D., Phyo, A.P., Tarning, J., Lwin, K.M., Ariey, F., Hanpithakpong, W., Lee, S.J., et al. (2009). Artemisinin Resistance in *Plasmodium falciparum* Malaria. N Engl J Med 361, 455–467.

Dziekan, J.M., Wirjanata, G., Dai, L., Go, K.D., Yu, H., Lim, Y.T., Chen, L., Wang, L.C., Puspita, B., Prabhu, N., et al. (2020). Cellular thermal shift assay for the identification of drug–target interactions in the *Plasmodium falciparum* proteome. Nature Protocols 15, 1881–1921.

Dziekan, J.M., Yu, H., Chen, D., Dai, L., Wirjanata, G., Larsson, A., Prabhu, N., Sobota, R.M., Bozdech, Z., and Nordlund, P. (2019). Identifying purine nucleoside phosphorylase as the target of quinine using cellular thermal shift assay. Science Translational Medicine 11, eaau3174.

Eastman, R.T., and Fidock, D.A. (2009). Artemisinin-based combination therapies: a vital tool in efforts to eliminate malaria. Nat Rev Microbiol 7, 864–874.

Favuzza, P., de Lera Ruiz, M., Thompson, J.K., Triglia, T., Ngo, A., Steel, R.W.J., Vavrek, M., Christensen, J., Healer, J., Boyce, C., et al. (2020). Dual Plasmepsin-Targeting Antimalarial Agents Disrupt Multiple Stages of the Malaria Parasite Life Cycle. Cell Host Microbe 27, 642–658.e612.

Garbarino, J., Pan, M., Chin, H.F., Lund, F.W., Maxfield, F.R., and Breslow, J.L. (2012). STARD4 knockdown in HepG2 cells disrupts cholesterol trafficking associated with the plasma membrane, ER, and ERC. J Lipid Res 53, 2716–2725.

Garnham, P.C., Bird, R.G., and Baker, J.R. (1962). Electron microscope studies of motile stages of malaria parasites. III. The ookinetes of Haemamoeba and Plasmodium. Trans R Soc Trop Med Hyg 56, 116–120.

Geoghegan, N.D., Evelyn, C., Whitehead, L.W., Pasternak, M., McDonald, P., Triglia, T., Marapana, D.S., Kempe, D., Thompson, J.K., Mlodzianoski, M.J., et al. (2021). 4D analysis of malaria parasite invasion offers insights into erythrocyte membrane remodeling and parasitophorous vacuole formation. Nature communications 12, 3620–3620.

Ghosh, A.K., and Jacobs-Lorena, M. (2009). *Plasmodium* sporozoite invasion of the mosquito salivary gland. Current Opinion in Microbiology 12, 394–400.

Gilson, P.R., and Crabb, B.S. (2009). Morphology and kinetics of the three distinct phases of red blood cell invasion by *Plasmodium falciparum* merozoites. Int J Parasitol 39, 91–96.

Goldberg, D.E., and Zimmerberg, J. (2020). Hardly Vacuous: The Parasitophorous Vacuolar Membrane of Malaria Parasites. Trends in Parasitology 36, 138–146.

Gómez-Díaz, E., Yerbanga, R.S., Lefèvre, T., Cohuet, A., Rowley, M.J., Ouedraogo, J.B., and Corces, V.G. (2017). Epigenetic regulation of *Plasmodium falciparum* clonally variant gene expression during development in Anopheles gambiae. Sci Rep 7, 40655.

Gurnett, A.M., Liberator, P.A., Dulski, P.M., Salowe, S.P., Donald, R.G.K., Anderson, J.W., Wiltsie, J., Diaz, C.A., Harris, G., Chang, B., et al. (2002). Purification and Molecular Characterization of cGMP-dependent Protein Kinase from Apicomplexan Parasites: A NOVEL CHEMOTHERAPEUTIC TARGET. J Biol Chem 277, 15913–15922.

Hill, R.J., Ringel, A., Knuepfer, E., Moon, R.W., Blackman, M.J., and van Ooij, C. (2016). Regulation and Essentiality of the StAR-related Lipid Transfer (START) Domain-containing Phospholipid Transfer Protein PFA0210c in Malaria Parasites. The Journal of biological chemistry 291, 24280–24292.

Iyer, L.M., Koonin, E.V., and Aravind, L. (2001). Adaptations of the helix-grip fold for ligand binding and catalysis in the START domain superfamily. Proteins 43, 134–144.

Jonsdottir, T.K., Counihan, N.A., Modak, J.K., Kouskousis, B., Sanders, P.R., Gabriela, M., Bullen, H.E., Crabb, B.S., de Koning-Ward, T.F., and Gilson, P.R. (2021). Characterisation of complexes formed by parasite proteins exported into the host cell compartment of Plasmodium falciparum infected red blood cells. Cellular Microbiology *n/a*, e13332.

Jonsdottir, T.K., Elsworth, B., Cobbold, S., Gabriela, M., Ploeger, E., Parkyn Schneider, M., Charnaud, S.C., Dans, M.G., McConville, M., Bullen, H.E., et al. (2023). PTEX helps efficiently traffic haemoglobinases to the food vacuole in *Plasmodium falciparum*. PLOS Pathogens 19, e1011006.

Jumper, J., Evans, R., Pritzel, A., Green, T., Figurnov, M., Ronneberger, O., Tunyasuvunakool, K., Bates, R., Žídek, A., Potapenko, A., et al. (2021). Highly accurate protein structure prediction with AlphaFold. Nature 596, 583–589.

Lapillo, M., Salis, B., Palazzolo, S., Poli, G., Granchi, C., Minutolo, F., Rotondo, R., Caligiuri, I., Canzonieri, V., Tuccinardi, T., et al. (2019). First-of-its-kind STARD(3) Inhibitor: In Silico Identification and Biological Evaluation as Anticancer Agent. ACS Med Chem Lett 10, 475–480.

Lautu-Gumal, D., Razook, Z., Koleala, T., Nate, E., McEwen, S., Timbi, D., Hetzel, M.W., Lavu, E., Tefuarani, N., Makita, L., et al. (2021). Surveillance of molecular markers of *Plasmodium falciparum* artemisinin resistance (kelch13 mutations) in Papua New Guinea between 2016 and 2018. Int J Parasitol Drugs Drug Resist 16, 188–193.

Lidumniece, E., Withers-Martinez, C., Hackett, F., Blackman, M.J., and Jirgensons, A. (2022). Subtilisin-like Serine Protease 1 (SUB1) as an Emerging Antimalarial Drug Target: Current Achievements in Inhibitor Discovery. Journal of Medicinal Chemistry 65, 12535–12545.

Lindner, S.E., Swearingen, K.E., Shears, M.J., Walker, M.P., Vrana, E.N., Hart, K.J., Minns, A.M., Sinnis, P., Moritz, R.L., and Kappe, S.H.I. (2019). Transcriptomics and proteomics reveal two waves of translational repression during the maturation of malaria parasite sporozoites. Nat Commun 10, 4964.

Llinás, M., Bozdech, Z., Wong, E.D., Adai, A.T., and DeRisi, J.L. (2006). Comparative whole genome transcriptome analysis of three *Plasmodium falciparum* strains. Nucleic Acids Res 34, 1166–1173.

López-Barragán, M.J., Lemieux, J., Quiñones, M., Williamson, K.C., Molina-Cruz, A., Cui, K., Barillas-Mury, C., Zhao, K., and Su, X.Z. (2011). Directional gene expression and antisense transcripts in sexual and asexual stages of *Plasmodium falciparum*. BMC Genomics 12, 587.

Makler, M.T., and Hinrichs, D.J. (1993). Measurement of the Lactate Dehydrogenase Activity of Plasmodium falciparum as an Assessment of Parasitemia. The American Journal of Tropical Medicine and Hygiene 48, 205–210.

Marapana, D.S., Dagley, L.F., Sandow, J.J., Nebl, T., Triglia, T., Pasternak, M., Dickerman, B.K., Crabb, B.S., Gilson, P.R., Webb, A.I., et al. (2018). Plasmepsin V cleaves malaria effector proteins in a distinct endoplasmic reticulum translocation interactome for export to the erythrocyte. Nat Microbiol 3, 1010–1022.

Margarida, R., Aline, F., and Christiaan van, O. (2023). Identification of a non-exported Plasmepsin V substrate that functions in the parasitophorous vacuole of malaria parasites. bioRxiv, 2023.2005.2015.540838.

Mazhari, R., Takashima, E., Longley, R.J., Ruybal-Pesantez, S., White, M.T., Kanoi, B.N., Nagaoka, H., Kiniboro, B., Siba, P., Tsuboi, T., et al. (2023). Identification of novel *Plasmodium vivax* proteins associated with protection against clinical malaria. Frontiers in Cellular and Infection Microbiology 13.

McHugh, E., Bulloch, M., Batinovic, S., Sarna, D., and Ralph, S. (2021). A divergent nonsense-mediated decay machinery in *Plasmodium falciparum* is inefficient and non-essential (bioRxiv).

Meis, J.F.G.M., and Ponnudurai, T. (1987). Ultrastructural studies on the interaction of *Plasmodium falciparum* ookinetes with the midgut epithelium of *Anopheles stephensi* mosquitoes. Parasitology research 73, 500–506.

Meis, J.F.G.M., Verhave, J.P., Jap, P.H.K., Sinden, R.E., and Meuwissen, J.H.E.T. (1983). Malaria parasites—discovery of the early liver form. Nature 302, 424–426.

Mikolajczak, S.A., Sacci, J.J.B., De La Vega, P., Camargo, N., VanBuskirk, K., Krzych, U., Cao, J., Jacobs-Lorena, M., Cowman, A.F., and Kappe, S.H.I. (2011). Disruption of the *Plasmodium falciparum* liver-stage antigen-1 locus causes a differentiation defect in late liver-stage parasites. Cellular Microbiology 13, 1250–1260.

Mok, S., Stokes, B.H., Gnädig, N.F., Ross, L.S., Yeo, T., Amaratunga, C., Allman, E., Solyakov, L., Bottrill, A.R., Tripathi, J., et al. (2021). Artemisinin-resistant K13 mutations rewire *Plasmodium falciparum’s* intra-erythrocytic metabolic program to enhance survival. Nature Communications 12, 530.

Mota, M.M., Pradel, G., Vanderberg, J.P., Hafalla, J.C., Frevert, U., Nussenzweig, R.S., Nussenzweig, V., and Rodríguez, A. (2001). Migration of *Plasmodium* sporozoites through cells before infection. Science 291, 141–144.

Mukherjee, S., Nguyen, S., Sharma, E., and Goldberg, D.E. (2022). Maturation and substrate processing topography of the *Plasmodium falciparum* invasion/egress protease plasmepsin X. Nature Communications 13, 4537.

Müller-Sienerth, N., Shilts, J., Kadir, K.A., Yman, V., Homann, M.V., Asghar, M., Ngasala, B., Singh, B., Färnert, A., and Wright, G.J. (2020). A panel of recombinant proteins from human-infective *Plasmodium* species for serological surveillance. Malar J 19, 31.

Nakao, N., Ueno, M., Sakai, S., Egawa, D., Hanzawa, H., Kawasaki, S., Kumagai, K., Suzuki, M., Kobayashi, S., and Hanada, K. (2019). Natural ligand-nonmimetic inhibitors of the lipid-transfer protein CERT. Communications Chemistry 2, 20.

Norcross, N.R., Wilson, C., Baragaña, B., Hallyburton, I., Osuna-Cabello, M., Norval, S., Riley, J., Fletcher, D., Sinden, R., Delves, M., et al. (2019). Substituted Aminoacetamides as Novel Leads for Malaria Treatment. ChemMedChem 14, 1329–1335.

Perkins, M. (1988). Stage-dependent processing and localization of a *Plasmodium falciparum* protein of 130,000 molecular weight. Experimental Parasitology 65, 61–68.

Pimenta, P.F., Touray, M., and Miller, L. (1994). The Journey of Malaria Sporozoites in the Mosquito Salivary Gland. Journal of Eukaryotic Microbiology 41, 608–624.

Pino, P., Caldelari, R., Mukherjee, B., Vahokoski, J., Klages, N., Maco, B., Collins, C.R., Blackman, M.J., Kursula, I., Heussler, V., et al. (2017). A multi-stage antimalarial targets the plasmepsins IX and X essential for invasion and egress. Science 358, 522–528.

Rathnapala, U.L., Goodman, C.D., and McFadden, G.I. (2017). A novel genetic technique in *Plasmodium berghei* allows liver stage analysis of genes required for mosquito stage development and demonstrates that de novo heme synthesis is essential for liver stage development in the malaria parasite. PLoS Pathog 13, e1006396.

Ressurreição, M., Fréville, A., and van Ooij, C. (2023). Identification of a non-exported Plasmepsin V substrate that functions in the parasitophorous vacuole of malaria parasites. bioRxiv, 2023.2005.2015.540838.

Ressurreição, M., and van Ooij, C. (2021). Lipid transport proteins in malaria, from *Plasmodium* parasites to their hosts. Biochim Biophys Acta Mol Cell Biol Lipids 1866, 159047.

Risco-Castillo, V., Topçu, S., Marinach, C., Manzoni, G., Bigorgne, Amélie E., Briquet, S., Baudin, X., Lebrun, M., Dubremetz, J.-F., and Silvie, O. (2015). Malaria Sporozoites Traverse Host Cells within Transient Vacuoles. Cell Host & Microbe 18, 593–603.

Rodriguez, M.H., and Hernández-Hernández, F.D.L.C. (2004). Insect-malaria parasites interactions: The salivary gland. Insect Biocehm Mol. Biol 34(7), 615–24.

Sanz, L.M., Crespo, B., De-Cózar, C., Ding, X.C., Llergo, J.L., Burrows, J.N., García-Bustos, J.F., and Gamo, F.-J. (2012). *P. falciparum* in vitro killing rates allow to discriminate between different antimalarial mode-of-action. PloS one 7, e30949–e30949.

Spielmann, T., Montagna, G.N., Hecht, L., and Matuschewski, K. (2012). Molecular make-up of the *Plasmodium* parasitophorous vacuolar membrane. International Journal of Medical Microbiology 302, 179–186.

Teuscher, F., Gatton, M.L., Chen, N., Peters, J., Kyle, D.E., and Cheng, Q. (2010). Artemisinin-Induced Dormancy in *Plasmodium falciparum*: Duration, Recovery Rates, and Implications in Treatment Failure. The Journal of Infectious Diseases 202, 1362–1368.

Trager, W., and Jensen, J.B. (1976). Human malaria parasites in continuous culture. Science 193, 673.

van Ooij, C., Withers-Martinez, C., Ringel, A., Cockcroft, S., Haldar, K., and Blackman, M.J. (2013). Identification of a *Plasmodium falciparum* phospholipid transfer protein. The Journal of biological chemistry 288, 31971–31983.

Van Vranken, J.G., Li, J., Mitchell, D.C., Navarrete-Perea, J., and Gygi, S.P. (2021). Assessing target engagement using proteome-wide solvent shift assays. eLife 10, e70784.

Varadi, M., Anyango, S., Deshpande, M., Nair, S., Natassia, C., Yordanova, G., Yuan, D., Stroe, O., Wood, G., Laydon, A., et al. (2022). AlphaFold Protein Structure Database: massively expanding the structural coverage of protein-sequence space with high-accuracy models. Nucleic Acids Res 50, D439–d444.

Vaughan, A.M., Mikolajczak, S.A., Wilson, E.M., Grompe, M., Kaushansky, A., Camargo, N., Bial, J., Ploss, A., and Kappe, S.H. (2012). Complete *Plasmodium falciparum* liver-stage development in liver-chimeric mice. J Clin Invest 122, 3618–3628.

Walz, A., Duffey, M., Aljayyoussi, G., Sax, S., Leroy, D., Besson, D., Burrows, J.N., Cherkaoui-Rbati, M.H., Gobeau, N., Westwood, M.-A., et al. (2023). The Parasite Reduction Ratio (PRR) Assay Version 2: Standardized Assessment of *Plasmodium falciparum* Viability after Antimalarial Treatment In Vitro. In Pharmaceuticals.

Weiss, G.E., Crabb, B.S., and Gilson, P.R. (2016). Overlaying Molecular and Temporal Aspects of Malaria Parasite Invasion. Trends Parasitol 32, 284–295.

Weiss, G.E., Gilson, P.R., Taechalertpaisarn, T., Tham, W.-H., de Jong, N.W.M., Harvey, K.L., Fowkes, F.J.I., Barlow, P.N., Rayner, J.C., Wright, G.J., et al. (2015). Revealing the Sequence and Resulting Cellular Morphology of Receptor-Ligand Interactions during *Plasmodium falciparum* Invasion of Erythrocytes. PLoS Pathog 11, e1004670.

Wilson, D.W., Goodman, C.D., Sleebs, B.E., Weiss, G.E., de Jong, N.W., Angrisano, F., Langer, C., Baum, J., Crabb, B.S., Gilson, P.R., et al. (2015). Macrolides rapidly inhibit red blood cell invasion by the human malaria parasite, *Plasmodium falciparum*. BMC Biol 13, 52–52.

Witkowski, B., Lelièvre, J., Barragán, M.J., Laurent, V., Su, X.Z., Berry, A., and Benoit-Vical, F. (2010). Increased tolerance to artemisinin in *Plasmodium falciparum* is mediated by a quiescence mechanism. Antimicrob Agents Chemother 54, 1872–1877.

Witmer, K., Dahalan, F.A., Delves, M.J., Yahiya, S., Watson, O.J., Straschil, U., Chiwcharoen, D., Sornboon, B., Pukrittayakamee, S., Pearson, R.D., et al. (2020). Transmission of Artemisinin-Resistant Malaria Parasites to Mosquitoes under Antimalarial Drug Pressure. Antimicrob Agents Chemother 65.

World Health Organization (2022). World Malaria Report 2022.

Yasuda, S., Kitagawa, H., Ueno, M., Ishitani, H., Fukasawa, M., Nishijima, M., Kobayashi, S., and Hanada, K. (2001). A novel inhibitor of ceramide trafficking from the endoplasmic reticulum to the site of sphingomyelin synthesis. J Biol Chem 276, 43994–44002.

Yoshida, N., Yamauchi, M., Morikawa, R., Hombhanje, F., and Mita, T. (2021). Increase in the proportion of *Plasmodium falciparum* with kelch13 C580Y mutation and decline in pfcrt and pfmdr1 mutant alleles in Papua New Guinea. Malaria Journal 20, 410.

Yu, C., Chen, X., Xu, W., Li, S., Chai, Q., and Zhang, Y. (2023). Solvent-induced proteome profiling for proteomic quantitation and target discovery of small molecular drugs. PROTEOMICS 23, 2200281.

Zenonos, Z.A., Rayner, J.C., and Wright, G.J. (2014). Towards a comprehensive *Plasmodium falciparum* merozoite cell surface and secreted recombinant protein library. Malaria Journal 13, 93.

Zhang, X., Lytovchenko, O., Lundström, S.L., Zubarev, R.A., and Gaetani, M. (2022). Proteome Integral Solubility Alteration (PISA) Assay in Mammalian Cells for Deep, High-Confidence, and High-Throughput Target Deconvolution. Bio Protoc 12.

Zhang, X., Wang, Q., Li, Y., Ruan, C., Wang, S., Hu, L., and Ye, M. (2020). Solvent-Induced Protein Precipitation for Drug Target Discovery on the Proteomic Scale. Anal Chem 92, 1363–1371.

Zieler, H., and Dvorak, J.A. (2000). Invasion in vitro of mosquito midgut cells by the malaria parasite proceeds by a conserved mechanism and results in death of the invaded midgut cells. Proc Natl Acad Sci U S A 97, 11516–11521.

