## Supplementary Figures and Tables for "Aryl amino acetamides prevent the development of *Plasmodium falciparum* rings via inhibition of the lipid transfer protein PfSTART1"

**Figure S1.**

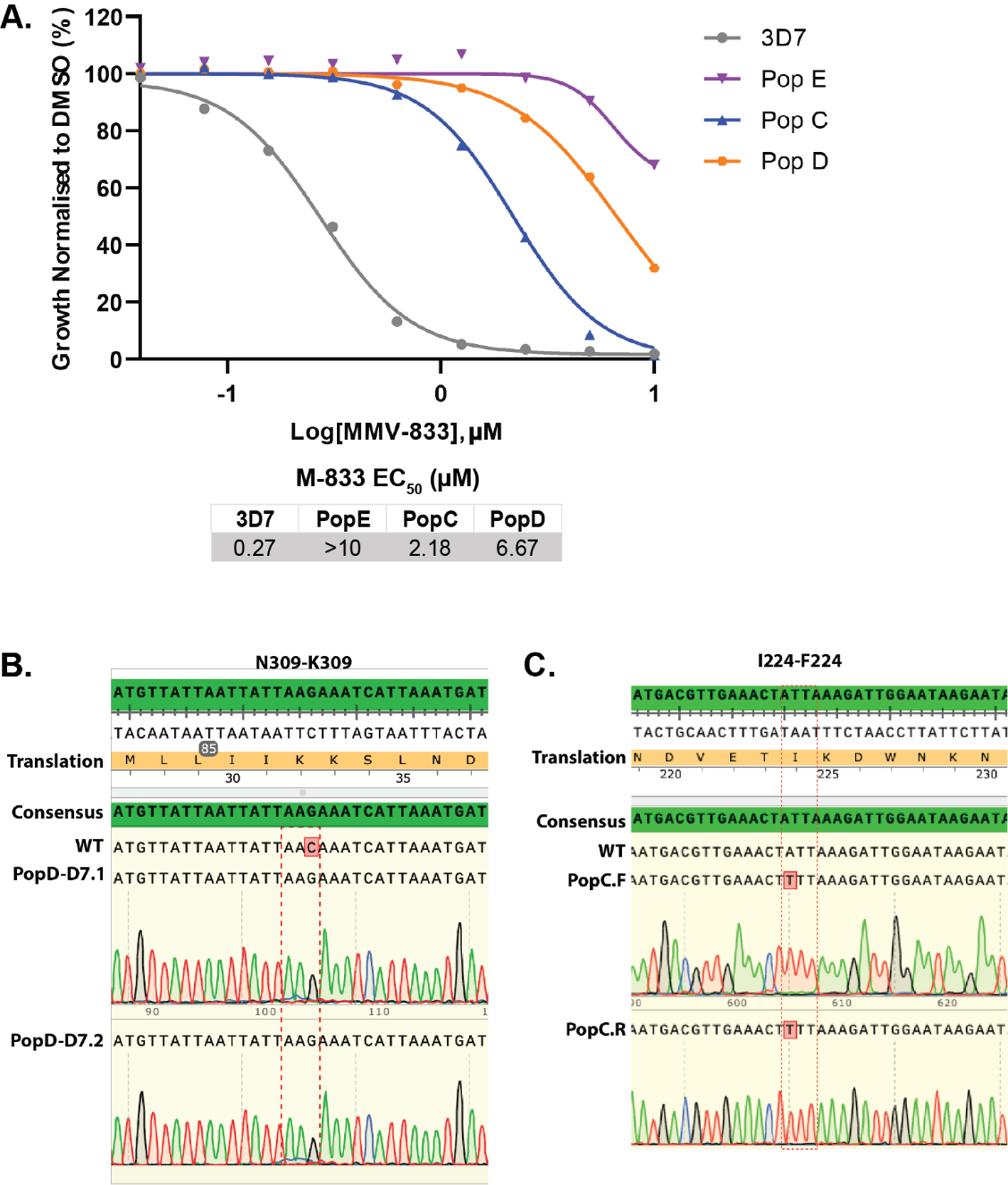

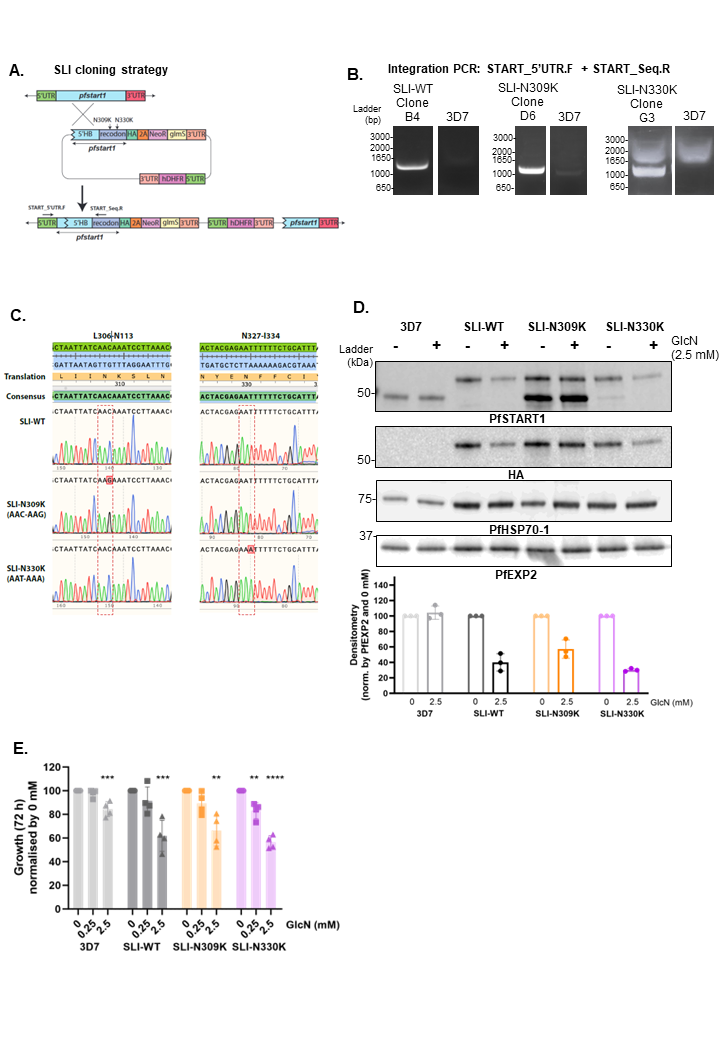
**Figure S2.**

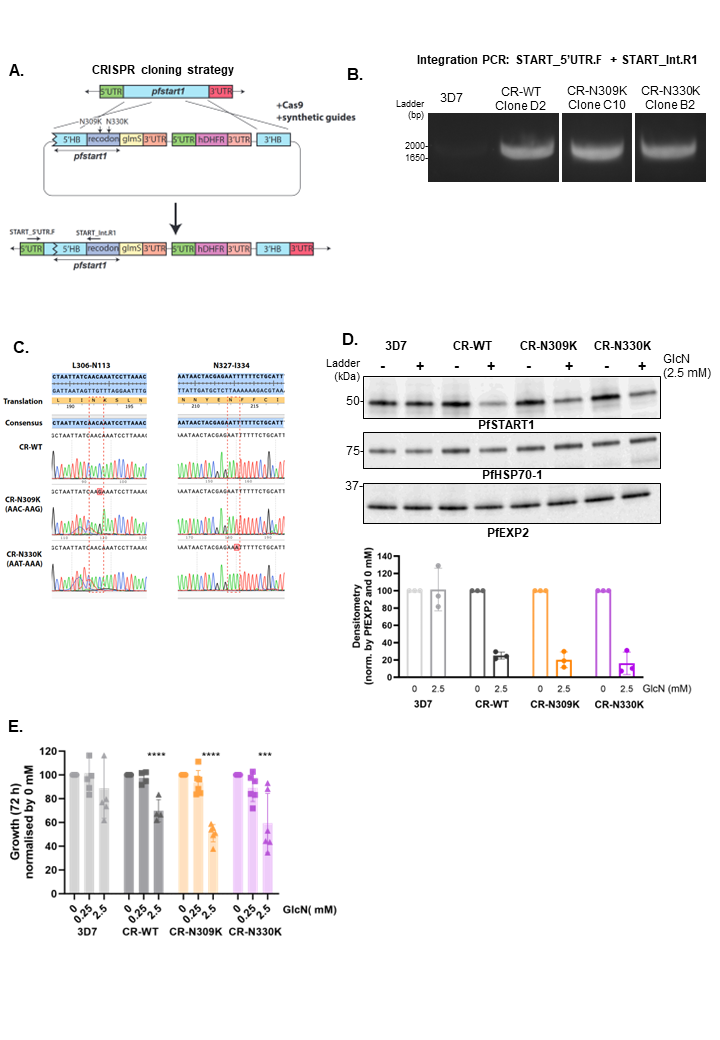
**Figure S3.**

**Figure S4.**

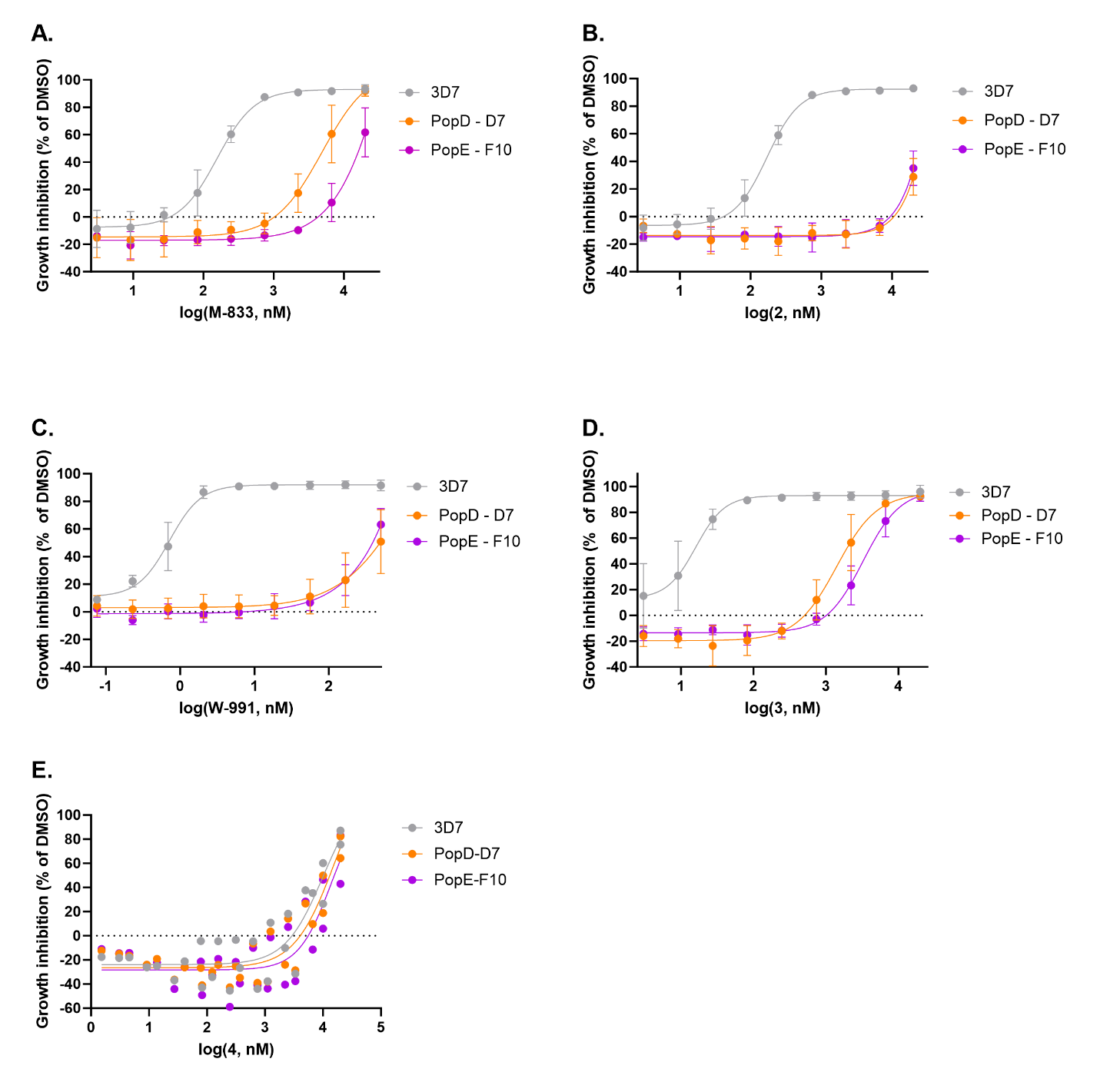

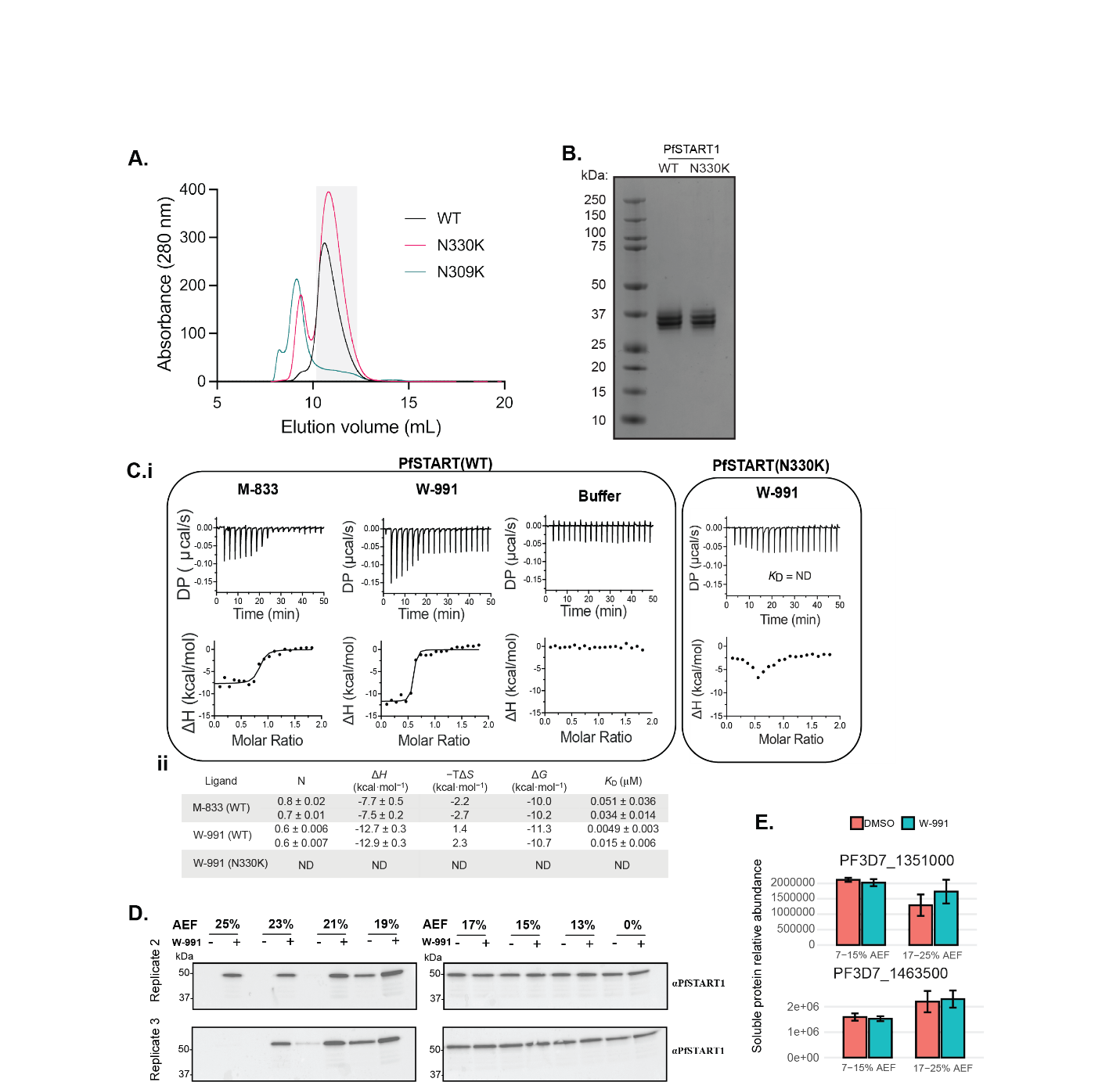
**Figure S5.**

**Figure S6.**

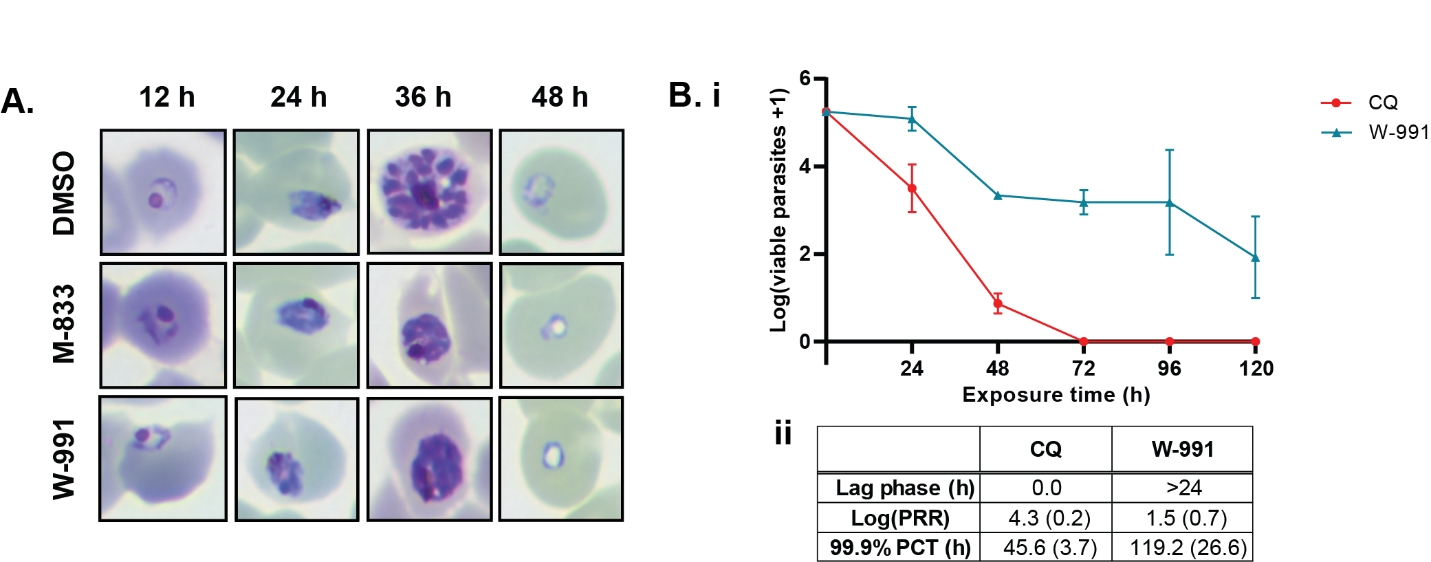

**Figure S7.**

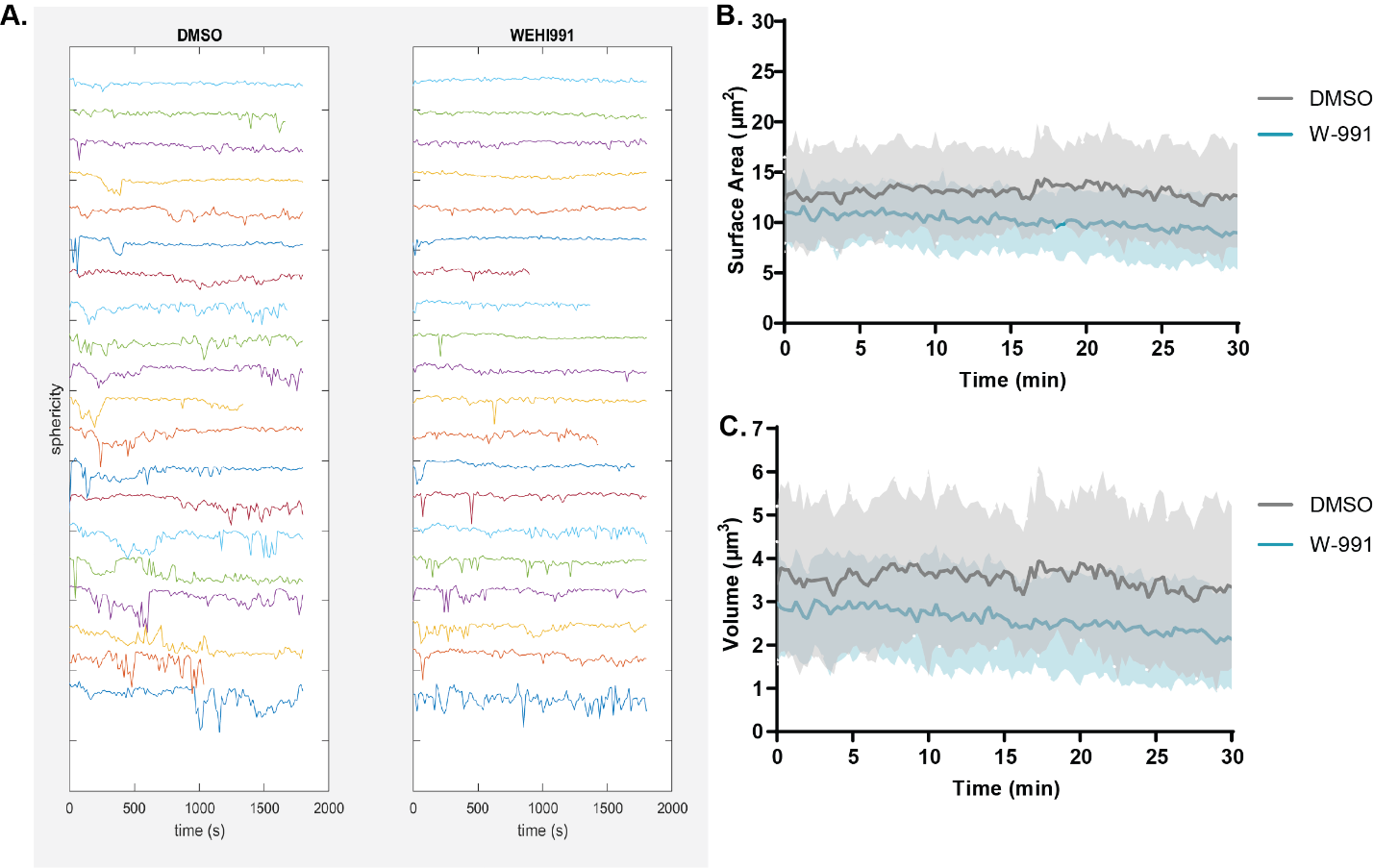

**Figure S8.**

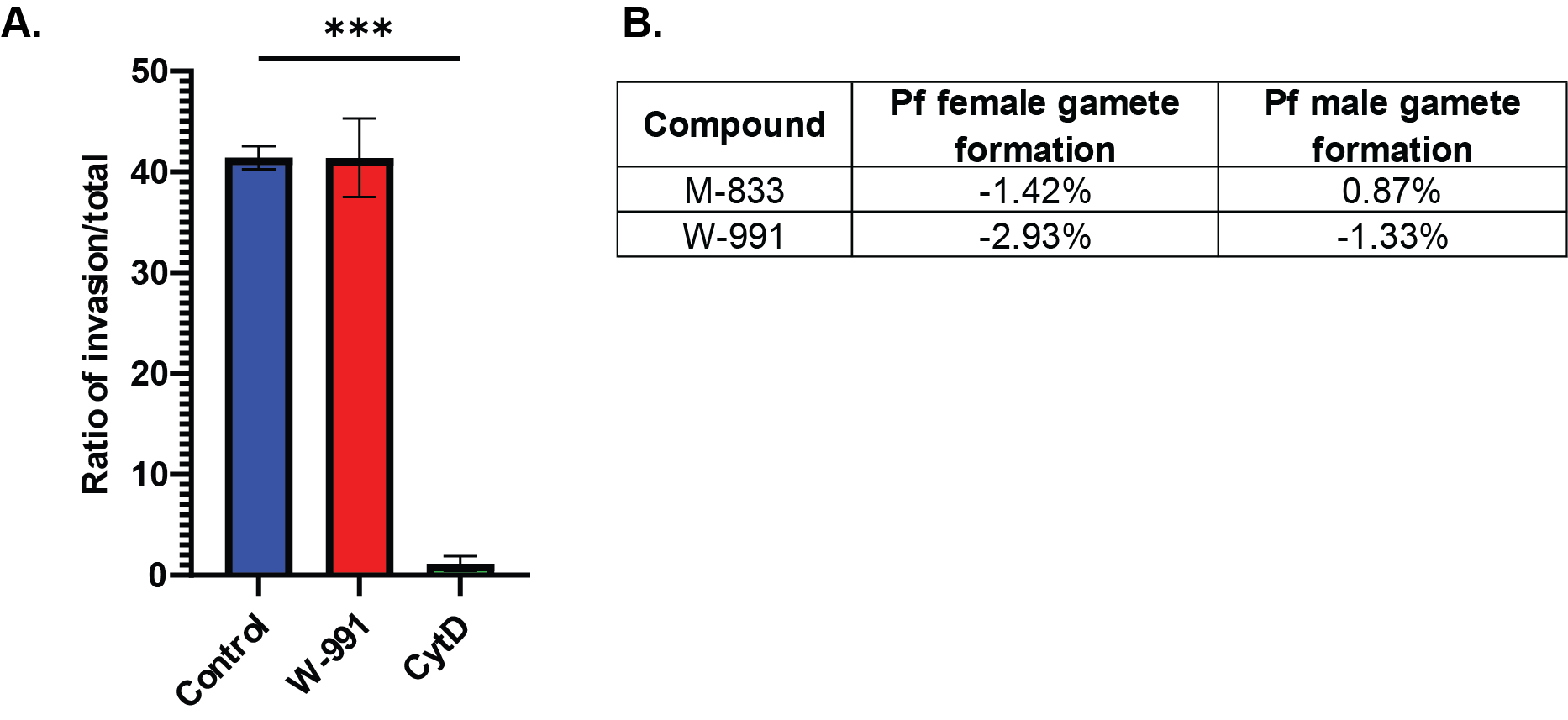

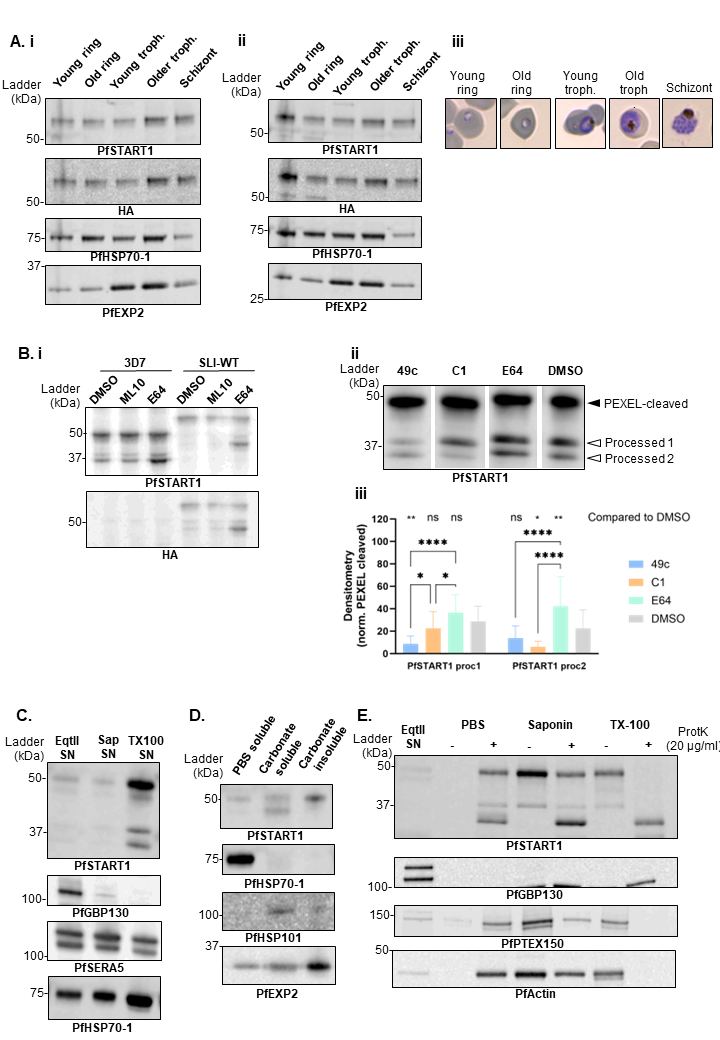
**Figure S9.**

**Figure S10.**

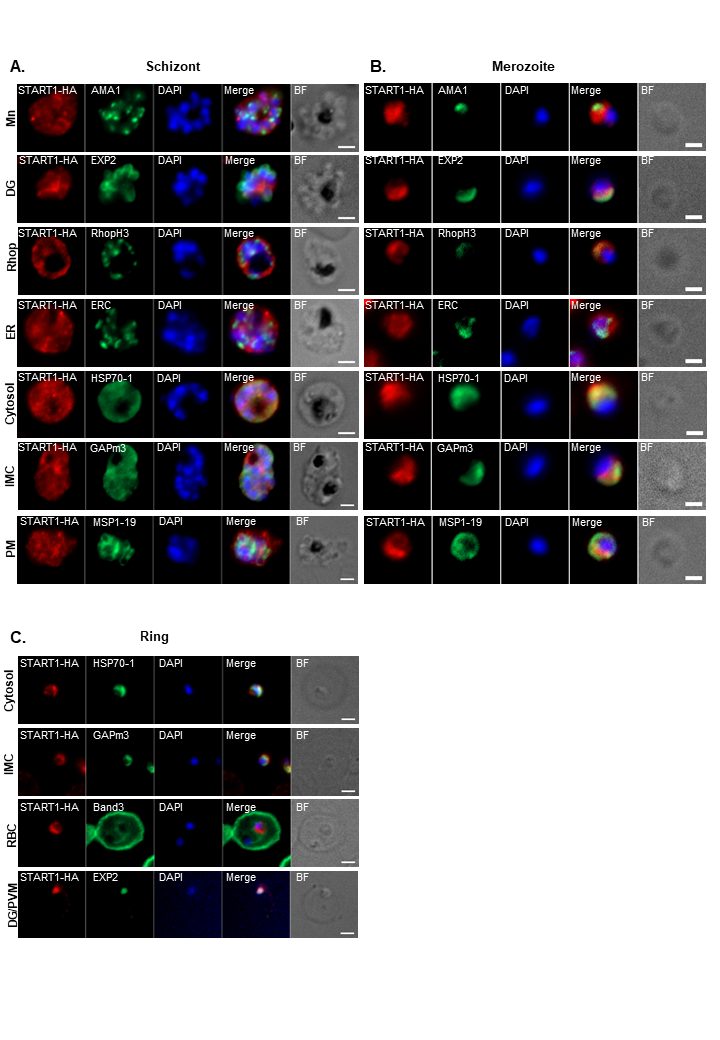

**Supplementary tables**

**Table S2.**

|  |  | **3D7** | **PopD – D7** | **PopE – F10** |
| --- | --- | --- | --- | --- |
| **M-833** | **EC_50_**  **(CI)** | 156 nM  (121.8 to 198.4) | 4.88 μM  (2.889-128.972) | >20 μM  (N/A) |
|  | Fold change compared to 3D7 | - | x 31 | > x 128 |
| **2** | **EC_50_**  **(CI)** | 171 nM  (142.7-202.8) | >20 μM  (N/A) | >20 μM  (N/A) |
|  | Fold change compared to 3D7 | - | > x 117 | > x 117 |
| **W-991** | **EC_50_**  **(CI)** | 0.72 nM  (0.5361-0.8852) | >200 nM  (N/A) | >200 nM  (N/A) |
|  | Fold change compared to 3D7 | - | > x 278 | > x 278 |
| **3** | **EC_50_**  **(CI)** | 16 nM  (6.950-?) | 1.4 μM  (1.016-2.019) | 3.25 μM  (2.575-4.443) |
|  | Fold change compared to 3D7 | - | x 88 | x 203 |
| **4** | **EC_50_**  **(CI)** | 10.03 μM  (3.685-?) | 15.73 μM  (5.059-?) | 13.84 μM  (9.919-?) |
|  | Fold change compared to 3D7 | - | x 1.6 | x 1.4 |

**Table S3.**

| **Primer name** | **Sequence** | **Purpose** |
| --- | --- | --- |
| SLTP_1F | AGATCTTTGTCTTTTGTTTTCTCTGTTGTGGAGCT | Amplify *pfstart* 5’ homology flank from 3D7 gDNA |
| SLTP_2R | CCAAGGAAGTCCATTAATAAGATATAAATATTTTCT |  |
| SLTP_3F | ATATCTTATTAATGGACTTCCTTGGCCaTTcAAgAGCCAGGAC | Amplify recodonised *pfstart* and add overlap with native 5’ flank and Pst cutting site |
| SLTP_PstR | CTGCAGcATCTTTATTGAAGAAGATACCGAATATTTTCTTGAAGAAGTTGA |  |
| SLTP_EcoR1 | GAATTCCCTTTTAAAAGTCAGGATACTATATATGAAGTGTATCA | Amplify *pfstart* 3’ homology flank from 3D7 gDNA |
| SLTP_KasR | GGCGCCTTAGTCCTTATTAAAAAATATACCAAATATTTTTTTAAAAAAGTTAACGT |  |
| START_5’UTR.F | TGTAATAATTTATACATTTTTATATCAGTTTATTTATTTTTGAGAAGGA | Amplify DNA only in parasites that have integrated the construct (used in SLI constructs) |
| START_Seq.R | ACTCATAACATATTTCACGCCTTTGTCATAAGAGT |  |
| START_Int.R1 | TCAAACACATCGTTTAAGGATTTGTTGATAATTAGCAGCA | With START_5’UTR.F: amplify DNA in parasites that have integrated the CRISPR construct |
| START_Int.R2 | ACCATATGAGTTTTCTGGGTCAGTCTGCACGTGC | With START_3F: used to amplify the *pfstart* locus to be sequenced in CRISPR parasites |
| Recodonised 3’ flank of START:   - WT (N309 and N330) - N309K mutant (orange, in brackets) - N330K mutant (purple, in brackets) | CCaTTcAAgAGCCAGGACACCATATACGAAGTGTACCAAAAATACTACAACAACAAGAATATGCTGCTAATTATC***AAC(AAG)***AAATCCTTAAACGATGTGTTTGATAATAATAGTAGTTATGCCAGGATAAATAACTACGAG***AAT(AAA)***TTTTTCTGCATTTACCCGAAATCAAAAAACTCTTACGACAAAGGTGTCAAGTACGTTATGAGCATCATATATGATGTGAATATCCCTAAGTTCATACAAAACAATATCCTAAACCAGATaTTCCCGGATCTTATATTCAATTTGCATAATACATCTATAGCTATAACTAATAAGACGGTGGGGACAGTGGTAGACTTATCCAAAAATGAACAGAATGCTTGGCACGCGCACAGTCTAAAAAACGTGAAACCAGAAGATACTCCAAATACTGAGCACGTGCAGACTGACCCAGAAAACTCATATGGTTTGGGGTTCATAAAGATGATATTTGTCGACGGTCCCTATAACCTGTGGATAATAAATGTCAACTTCTTCAAGAAAATATTCGGTATCTTCTTCAATAAAGAT | |
| gRNA | /AltR1/rArUrArGrUrArUrCrCrUrGrArCrUrU | Guides the recombinant Cas9 enzyme to cut the native *pfstart* locus (but not the recodonised *pfstart* DNA template) |

**Table S4.**

| **Antibody** | **Raised in:** | **Provider** | **Dilution** (WB: western blot; IFA: immunofluorescent assay) |
| --- | --- | --- | --- |
| HA | Mouse | Sigma | 1:1000 (WB)  1:500 (IFA) |
| PfSTART | Rabbit | WEHI (see Methods) | 1:1000 (WB) |
| PfEXP2 | Mouse | WEHI (PMID: 19536257) | 1:2000 (WB) |
| PfEXP2 | Rabbit | WEHI (PMID: 19536257) | 1:2000 (WB)  1:500 (IFA) |
| PfHSP70.1 | Rabbit | WEHI (PMID: 28732045) | 1:2000 (WB)  1:500 (IFA) |
| PfGBP130 | Mouse | WEHI | 1:1000 (WB) |
| PfSERA5 | Rabbit | WEHI (PMID: 13679369) | 1:1000 (WB) |
| PfHSP101 | Rabbit | WEHI (PMID: 19536257) | 1:1000 (WB) |
| PfActin | Mouse | WEHI (PMID: 22389687) | 1:500 (WB) |
| PfAMA1 | Rabbit | PMID: 29165651 | 1:500 (IFA) |
| PfRhopH3 | Rabbit | PMID: 28252383 | 1:500 (IFA) |
| PfERC | Rabbit | PMID: 9364972 | 1:500 (IFA) |
| PfMSP1-19 | Rabbit | PMID: 19561073 | 1:500 (IFA) |
| PfGAPm3 N-term | Rabbit | Made by Genscript to C-term peptide. | 1:500 (IFA) |
| Band3 | Rabbit | Abcam | 1:500 (IFA) |
| Rabbit-Alexa Fluor Plus 680 | Goat | Invitrogen | 1:10000 (WB) |
| Mouse-Alexa Fluor Plus 800 | Goat | Invitrogen | 1:10000 (WB) |
| Rabbit-Alexa Fluor 488 | Goat | Invitrogen | 1:2000 (IFA) |
| Mouse-Alexa Fluor 594 | Goat | Invitrogen | 1:2000 (IFA) |
